# CellRank for directed single-cell fate mapping

**DOI:** 10.1101/2020.10.19.345983

**Authors:** Marius Lange, Volker Bergen, Michal Klein, Manu Setty, Bernhard Reuter, Mostafa Bakhti, Heiko Lickert, Meshal Ansari, Janine Schniering, Herbert B. Schiller, Dana Pe’er, Fabian J. Theis

## Abstract

Computational trajectory inference enables the reconstruction of cell-state dynamics from single-cell RNA sequencing experiments. However, trajectory inference requires that the direction of a biological process is known, largely limiting its application to differentiating systems in normal development. Here, we present CellRank (https://cellrank.org) for mapping the fate of single cells in diverse scenarios, including perturbations such as regeneration or disease, for which direction is unknown. Our approach combines the robustness of trajectory inference with directional information from RNA velocity, derived from ratios of spliced to unspliced reads. CellRank takes into account both the gradual and stochastic nature of cellular fate decisions, as well as uncertainty in RNA velocity vectors. On data from pancreas development, we show that it automatically detects initial, intermediate and terminal populations, predicts fate potentials and visualizes continuous gene expression trends along individual lineages. CellRank also predicts a novel dedifferentiation trajectory during regeneration after lung injury, which we follow up experimentally by confirming the existence of previously unknown intermediate cell states.

## Introduction

Cells undergo state transitions during many biological processes, including development, reprogramming, regeneration, cell cycle and cancer, and they often do so in a highly asynchronous fashion^1^. Single-cell RNA sequencing (scRNA-seq) has been very successful at capturing the heterogeneity of these processes as they unfold in individual cells. However, lineage relationships are lost in scRNA-seq due to its destructive nature—cells cannot be measured multiple times. Experimental approaches have been proposed to mitigate this problem; scRNA-seq can be combined with lineage tracing methods^2–5^ that use heritable barcodes to follow clonal evolution over long time scales, and metabolic labeling^6–9^ uses the ratio between nascent and mature RNA molecules to statistically link observed gene expression profiles over short time windows. Yet both strategies are mostly limited to *in vitro* applications, prompting the development of computational approaches to reconstruct pseudotime trajectories^1,10–16^. These approaches are based on the observation that developmentally related cells tend to share similar gene expression profiles, and they have been used extensively to compute pseudotemporal orderings of cells along differentiation trajectories and to study cell-fate decisions.

Computational trajectory inference typically demands the use of prior biological knowledge to determine the directionality of cell-state changes, often by specifying an initial cell^17^. This has largely limited the success of such approaches to normal developmental scenarios with known cell hierarchies. The recently introduced concept of RNA velocity^18^ alleviates this problem by reconstructing the direction of state-change trajectories based on the ratio of spliced to unspliced mRNA molecules. The approach has been generalized to include transient cell populations and protein kinetics^19,20^; however, velocity estimates are noisy and the interpretation of high-dimensional velocity vectors has mostly been limited to low-dimensional projections, which do not easily reveal long-range probabilistic fates or allow quantitative interpretation.

Here, we present CellRank, a method that combines the robustness of previous similarity-based trajectory inference methods with directional information given by RNA velocity to learn directed, probabilistic state-change trajectories under either normal or perturbed conditions. CellRank infers initial, intermediate and terminal populations of an scRNA-seq dataset and computes fate probabilities, which we use to uncover putative lineage drivers and to visualize lineage-specific gene expression trends. For these computations, we take into account both the stochastic nature of cellular fate decisions as well as uncertainty in the velocity estimates. We demonstrate CellRank’s capabilities on pancreatic endocrine lineage development, where we identify established terminal and initial states and correctly recover lineage bias and key driver genes for somatostatin-producing delta cell differentiation. Further, we apply CellRank to lung regeneration, where we predict a novel dedifferentiation trajectory and experimentally validate the existence of previously unknown intermediate cell states. We show that CellRank outperforms methods that do not include velocity information. CellRank is available as a scalable, user friendly open source software package with documentation and tutorials at https://cellrank.org.

## Results

### CellRank combines cell-cell similarity with RNA velocity to model cellular state transitions

The CellRank algorithm aims to model the cell state dynamics of a system. First, CellRank detects the initial, terminal and intermediate cell states of the system and computes a global map of fate potentials, assigning each cell the probability of reaching each terminal state. Based on the inferred potentials, CellRank can chart gene expression dynamics as cells take on different fates, and it can identify putative regulators of cell-fate decisions. The algorithm uses an scRNA-seq count matrix and a corresponding RNA velocity vector matrix as input. Note that while we use RNA velocity here to approximate the direction of cellular dynamics, CellRank generalizes to accommodate any directional measure which gives a vector field representation of the process, such as metabolic labeling^6–9^ or real time information^21,22^.

The main assumption underlying all pseudotime algorithms that faithfully capture trajectories^10–14^ is that cell states change in small steps with many transitional populations^1^. CellRank uses the same assumption to model state transitions using a Markov chain, where each state in the chain is given by one observed cellular profile, and edge weights denote the probability of transitioning from one cell to another. The first step in chain construction is to compute an undirected k-nearest neighbor (KNN) graph representing cell-cell similarities in the phenotypic manifold (Fig. 1a,b, Supplementary Fig 1a and Online Methods). Each node in the graph represents an observed cellular profile, and edges connect cells which are most similar.

**Figure 1:**
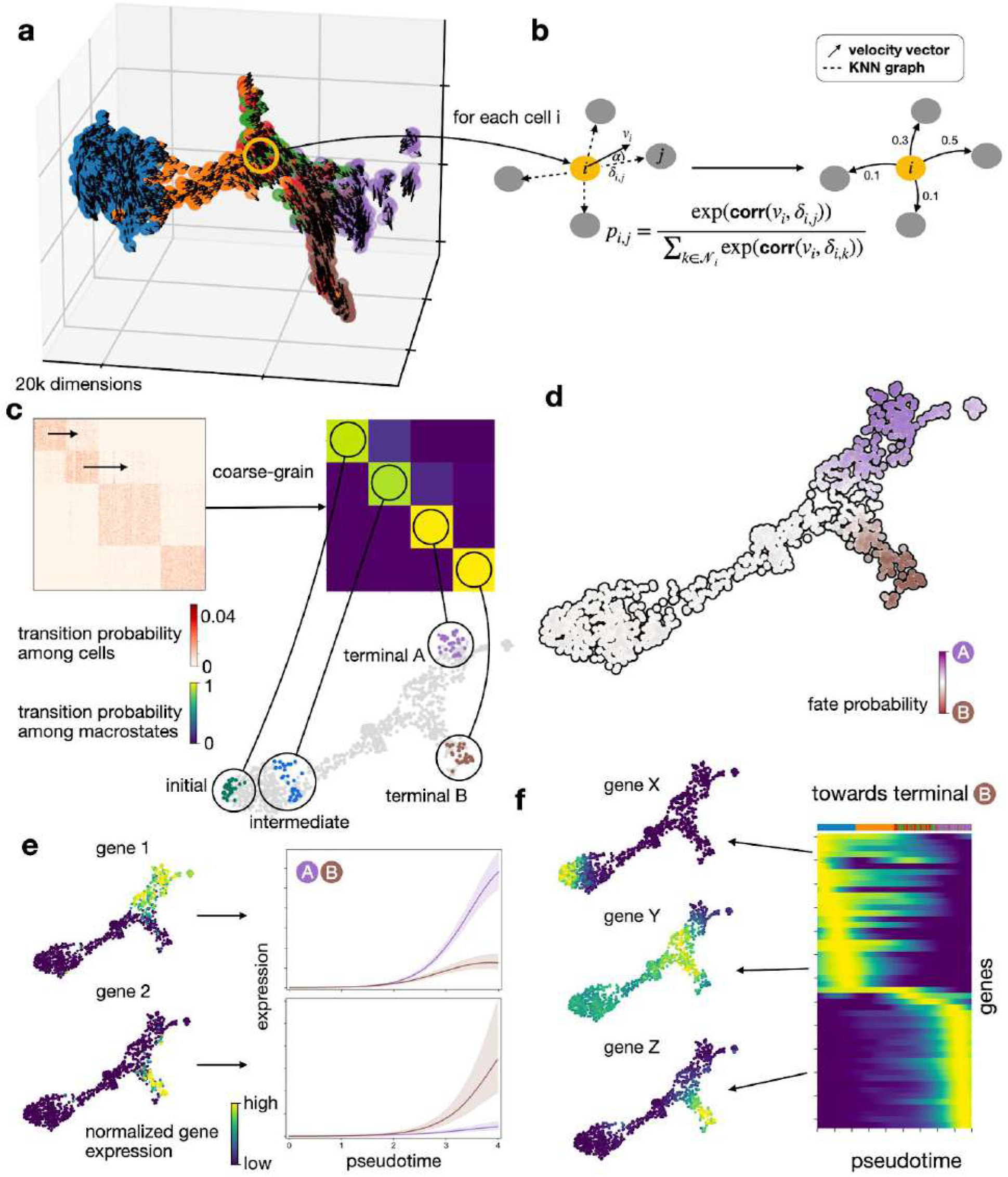
Combining RNA velocity with cell-cell similarity to determine initial and terminal states and compute a global map of cellular fate potential. **a.** 3D UMAP of 1000 simulated cells with their velocity vectors, using DynGen^84^. Colors reflect DynGen ground truth branch assignment. CellRank models cell state transitions directly in high dimensional gene expression space. **b.** A reference cell *i* with velocity vector *v_i_* and its nearest neighbors. The vector *δ_ij_* is the difference in gene expression between cells *j* and *i*. To assign probability *p_ij_*. to cell *i* transitioning to cell *j* in the neighborhood *N_t_* of cell *i*, we transform correlations between the transcriptomic difference vectors *δ_ij_* and the velocity vector *v_i_*, essentially considering the angle *α* between these vectors. **c**. The directed transition matrix is coarse-grained into 4 macrostates. Heat maps show transition probabilities among cells (left) and macrostates (right); sorting cells according to macrostate membership recovers block structure in the cell-cell transition matrix. We recover initial, intermediate and two terminal states. The 30 colored cells are mostly likely to belong to each macrostate in the UMAP. **d.** For each cell not assigned to either A or B, we compute its probability of reaching A or B. We show these fate probabilities in a fate map where each cell is colored according to the terminal state it is most likely to reach. Color intensity reflects the degree of lineage priming. **e.** Using these fate probabilities and a pseudotime, we plot gene expression trends which are specific to either A or B. Left each cell is colored based on the expression of the indicated genes and the respective trends along pseudotime towards each fate is shown on the right. **f**. We show expression trends in pseudotime of the top 50 genes whose expression correlates best with the probability of reaching B in a heatmap. Genes have been sorted according to their smoothed peak in pseudotime. We highlight one early gene (X), one intermediate gene (Y) and one late gene (Z) by showing expression in the UMAP.

Unlike pseudotime algorithms, we infuse directionality by using RNA velocity to direct Markov chain edges. The RNA velocity vector of a given cell predicts which genes are currently being up- or downregulated and points towards the likely future state of that cell. The more a neighboring cell lies in the direction of the velocity vector, the higher its transition probability (Online Methods). We compute a second set of transition probabilities based on gene expression similarity between cells and combine it with the first set via a weighted mean (Online Methods). The resulting matrix of directed transition probabilities is independent of any low-dimensional embedding and reflects transcriptional similarity as well as directional information given by RNA velocity.

The transition matrix may be extremely large, noisy and difficult to interpret. We alleviate these problems by summarizing individual gene expression profiles into macrostates, regions of the phenotypic manifold that cells are unlikely to leave (Fig. 1c, Supp Fig 1c-e). CellRank decomposes the dynamics of the Markov chain into these macrostates and computes coarse-grained transition probabilities among them. The number of macrostates is a model parameter that can be chosen using knee-point heuristics or prior knowledge about the biological system (Supplementary Fig 1b and Online Methods). Individual cells are assigned to macrostates via a soft assignment. To compute macrostates and the induced coarse-grained transition probabilities, we adapt the *Generalized Perron Cluster Cluster Analysis* (G-PCCA)^23,24^ to the single-cell context (Online Methods).

Viewing the biological system at coarse resolution allows us to identify populations based on transition probabilities: terminal macrostates will have high self-transition probability, initial macrostates will have low incoming transition probability, and remaining macrostates will be intermediate. We automate the identification of terminal states through a stability index (SI) between zero and one, indicating self-transition probability (Online methods); macrostates with an SI of 0.96 or greater are classified as terminal. We automate the identification of initial states though the coarse-grained stationary distribution (CGSD), which describes the long-term evolution of the coarse-grained Markov chain (Online methods). The CGSD assigns small values to macrostates that the process is unlikely to revisit after leaving; these macrostates are classified as initial. The number of initial states is a parameter that is set to one by default.

Finally, CellRank uses the directed single-cell transition matrix to compute fate probability, the likelihood that a given cell will ultimately transition towards each terminal population defined in the previous step (Fig. 1d and Supplementary Fig. 1f). These probabilities can be efficiently computed for all cells by solving a linear system in closed form (Online Methods). Fate probabilities extend the short-range fate prediction given by RNA velocity to the global structure spanning initial to terminal states. The stochastic Markov chain-based formulation allows us to overcome noise in individual velocity vectors and cell-cell similarities by aggregating many of these into our final fate prediction.

We combine fate probability estimates with a pseudotemporal ordering to visualize gene expression programs executed by cells along trajectories leading to terminal states (Fig. 1e and Online Methods). Pseudotime orders a progression of cell states from the initial state, while CellRank fate probabilities indicate how committed each cell is to every trajectory. By softly assigning cells to trajectories via fate probabilities, we capture the effect of gradual lineage commitment, whereby cells transition from an uncommitted state (contribution to several trajectories) to a committed state (contribution to a single trajectory)^25–28^. Palantir^25^, which is based on an iteratively refined shortest path in the space of diffusion components, is used for pseudotime ordering by default, where Palantir is provided with CellRank’s computed initial state. By correlating gene expression with fate probabilities, CellRank enhances the ability to uncover putative trajectory-specific regulators (Fig. 1f). By sorting putative regulators according to their peak in pseudotime, we visualize gene expression cascades specific to their cellular trajectory while accounting for the continuous nature of cellular fate commitment.

### Macrostates resolve initial and terminal states of pancreatic endocrine lineage formation

We applied CellRank to an scRNA-seq dataset of E15.5 murine pancreatic development^29^. A UMAP^30^ representation with original cluster annotations and scVelo-projected velocities recapitulated the main developmental trends^19^ (Fig 2a); from an initial cluster of endocrine progenitors (EPs) expressing low levels of the transcription factor neurogenin 3 *(Neurog3* or *Ngn3*), cells traverse trajectories towards alpha, beta, epsilon and delta cell fates.

**Figure 2:**
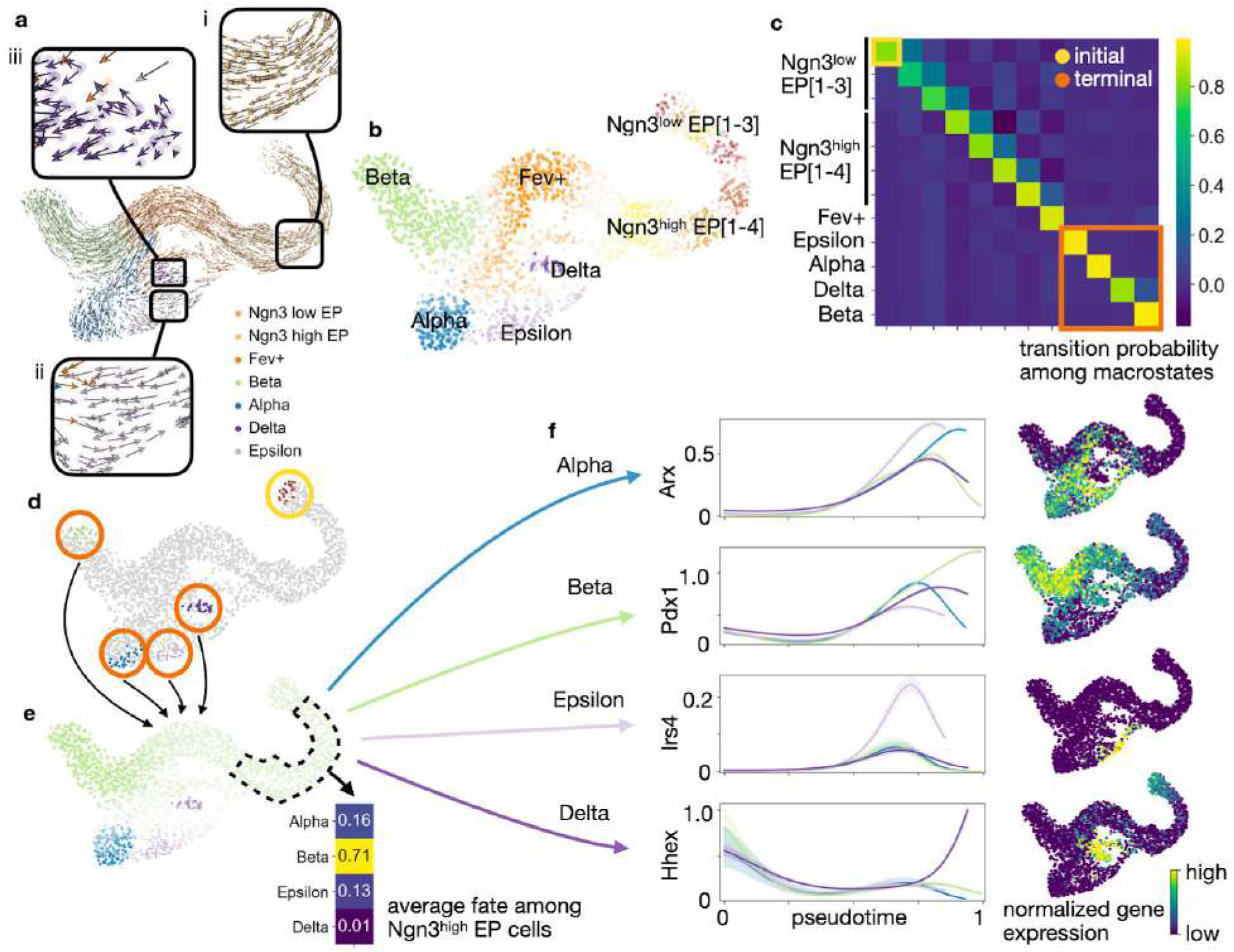
Delineating fate choice in pancreatic development. **a.** UMAP of murine pancreatic development at E15.5 with scVelo projected velocities. Colors correspond to published cluster annotations^29^. CellRank provides additional insights regarding (i) the fate of early cells, (ii) the identification of terminal states and (iii) likely progenitors of terminal fates (boxed insets). **b**. Soft assignment of cells to macrostates. Cells colored by most likely macrostate; color intensity reflects degree of confidence, and grey cells reside between multiple macrostates. **c**. Coarse-grained transition probabilities among macrostates. Terminal macrostates are outlined in red and the initial Ngn3^low^ EP_1 macrostate is outlined in yellow. **d**. Highlight of the 30 cells most confidently assigned to each initial and terminal macrostate, colored as in (b). e. UMAP displaying probabilities for reaching alpha, beta, epsilon and delta terminal fates. Fates colored as in (b), with darker color indicating higher probability. Inset shows average fate probabilities of cells in the Ngn3^high^ EP cluster marked with a dashed line. **f**. Smoothed gene expression trends in pseudotime for the lineage determinants Arx^39^ (alpha), Pdx1^40^ (beta) and Hhex^41^ (delta) as well as the lineage associated gene Irs4^42^ (epsilon). The trend for each gene is shown for each trajectory leading to the indicated terminal population. Right column, expression values for the corresponding gene in the UMAP.

To investigate specific questions such as the onset of lineage bias, precise location of initial and terminal states, and likely progenitors of any terminal state, we argue against basing hypotheses purely on the projected velocity vectors for three reasons. First, the vectors are projected onto only 2 or 3 dimensions, which may over-regularize the true velocities and lead to overly smooth vector fields. Interpreting cellular trajectories in 2D or 3D embeddings is often misleading, as high-dimensional distances cannot be fully preserved in lower dimensions; this is why most neighborhood-based dimensionality reduction techniques such as t-SNE^31,32^ or UMAP^30,33^ do not conserve global relationships well^34–36^. Second, visual interpretation of projected vectors ignores uncertainty in RNA velocity and therefore leads to overconfidence in the inferred trajectories. Third, velocities are only available locally, whereas CellRank aggregates these local signals globally, computing longer range trends. Similar to the consensus reached by the single cell field to avoid clustering cells in 2D or 3D representations^37^, we argue that velocity vectors projected onto two or three dimensions must not be used to address detailed questions of trajectory inference. CellRank overcomes these limitations and allows us to model global trajectories, as we demonstrate on pancreas data below.

We computed CellRank’s directed transition matrix and then coarse-grained it into 12 macrostates (Fig. 2b) based on eigenvalue gap analysis (Supplementary Fig. 2a,b), revealing a block-like structure in the transition matrix (Fig. 2c and Supplementary Fig. 3a-c). Macrostates, annotated according to their overlap with the underlying gene expression clusters (Online Methods), comprised all developmental stages in this dataset, from an initial Ngn3^low^ EP state, to intermediate Ngn3^high^ EP and Fev+ states, to terminal hormone-producing alpha, beta, epsilon and delta cell states.

The three most stable states according to the coarse-grained transition matrix were alpha (SI 0.97), beta (SI 1.00) and epsilon (SI 0.98) macrostates, which were accordingly labeled as terminal by CellRank, consistent with known biology (Fig. 2c). Additionally, we recovered one relatively stable (SI 0.84) macrostate which largely overlapped with delta cells. We identified the Ngn3^low^ EP_1 state as initial because it was assigned the smallest CGSD value (2 x 10^-6^). The location of initial and terminal states agrees with the expression of well-known marker genes, including *Ins1* and *Ins2* for beta, *Gcg* for alpha, *Ghrl* for epsilon, *Sst* for delta cells and ductal cell markers *Sox9, Anxa2* and *Bicc1* for the initial state^29,38^ (Fig. 2d and Suppl. Fig. 4a,b).

We computed fate probabilities and summarized them in a fate map (Fig. 2e). This analysis correctly identified the beta cell fate as dominant within the Ngn3^high^ EP cluster at E15.5, consistent with known biology^29^ (Fig. 2e, inset), as also visualized with pie charts on a directed implementation of partition-based graph abstraction^11^ (PAGA) (Online Methods and Supplementary Fig. 8a,b). Using a cell within the Ngn3^low^ EP_1 macrostate as the starting state for Palantir^25^, we ordered cells in pseudotime (Supplementary Fig. 6a-c) and overlaid the expression of master regulators *Arx^39^* (alpha), *Pdx1^40^* (beta) and *Hhex^41^* (delta), and the lineage-associated gene *Irs4^42^* (epsilon) (Fig. 2f) to visualize trends. All of these genes were correctly upregulated when approaching their associated terminal populations.

All components of CellRank are extremely robust to parameter variation, based on sensitivity analysis for the number of macrostates (Supplementary Fig. 7a-f), number of neighbors in the KNN graph, scVelo minimal gene counts, number of highly variable genes and number of principal components. Additionally, CellRank is robust to random subsampling of cells (Supplementary Figs. 8a-e and 9a-e).

### CellRank identifies putative gene programs driving delta cell differentiation

Delta cells highlight how CellRank’s global approach overcomes limitations in RNA velocity. Delta cells are very rare in our data (70 cells or 3% of total, Supplementary Fig. 10a,b) and more importantly, no known drivers of delta cell development were among scVelo’s 30 genes with highest likelihoods (Supplementary Fig. 11a). Moreover, genes implicated in delta cell development were not captured well by scVelo’s model of splicing kinetics (Supplementary Fig. 11b,c). We hypothesize that splicing kinetics fail to capture delta cell differentiation because these cells appear late in pancreatic development and thus are very rare in our data^43^.

The development of delta cells is not well understood^38^. Mature delta cells can be identified by *Sst* expression (Supplementary Fig. 13a,b), but immature cells are much more difficult to identify. *Hhex* is the only widely accepted lineage marker^41^, and *Cd24a* has recently been implicated in human delta cell development^44,45^. To learn more about delta cell development, we focused on CellRank fate probabilities towards the relatively stable delta macrostate (SI 0.84) which was not automatically classified as terminal^38^ (Fig. 3a). Velocities projected onto the UMAP show multiple possible paths towards delta (Supplementary Fig. 12a), but CellRank fate probabilities reveal one path with highest likelihood, through cells that were annotated as delta precursors in a study involving subclustering of the Fev+ population (Fig. 3c and Supplementary Fig. 10c)^29^. Therefore, while RNA velocity fails to capture the dynamics of delta cell development, they can be successfully recovered by CellRank because it constrains velocities to the phenotypic manifold via the KNN graph, incorporates cell-cell similarly and models long-range trends.

**Figure 3:**
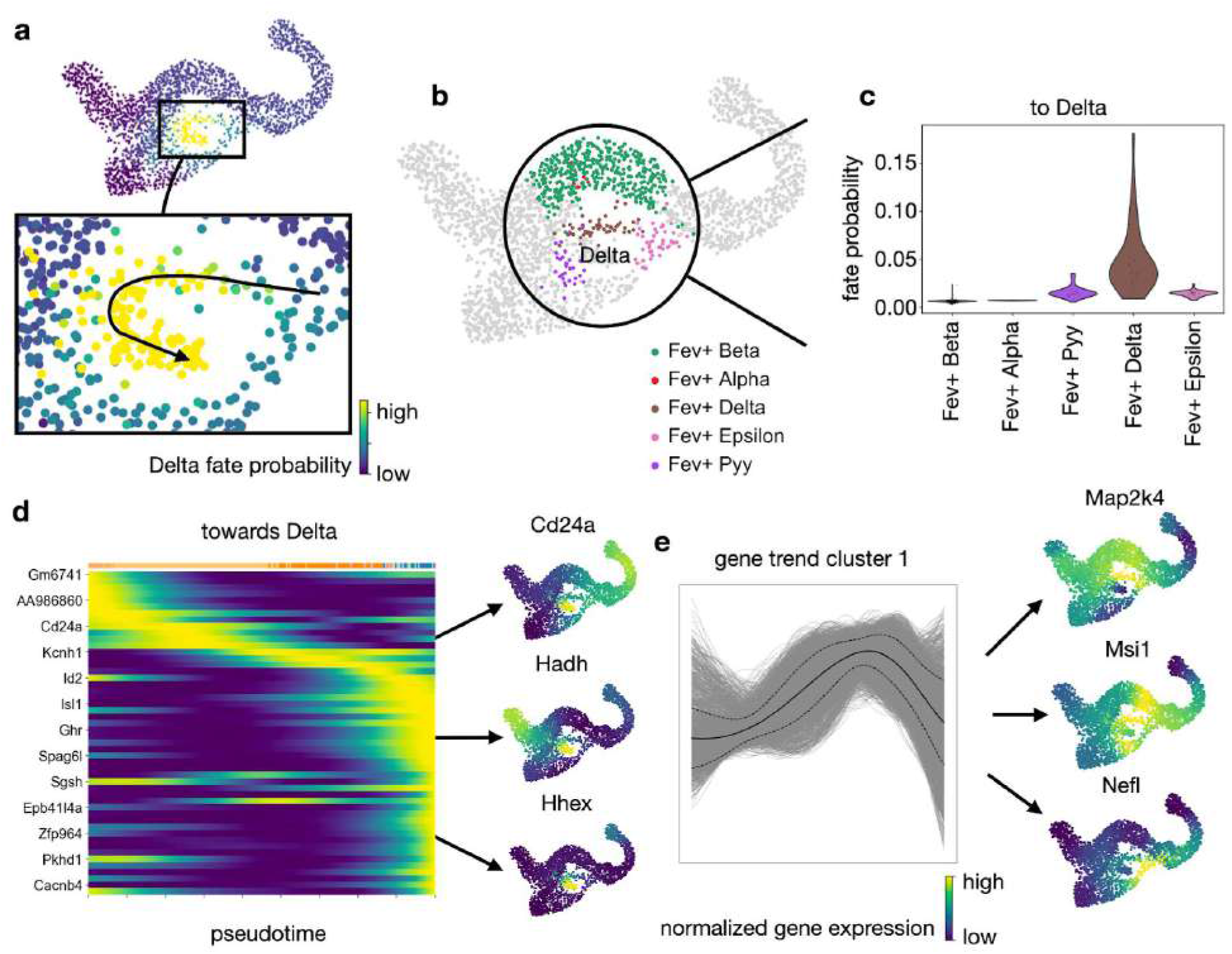
Zooming into the delta state to elucidate differentiation paths. **a**. CellRank probabilities for acquiring the terminal delta cell fate (see Fig. 2d). Cells are colored by the probability of reaching the delta state. Inset, group of cells likely to become delta, showing differentiation path predicted by CellRank (arrow). **b**. Orthogonal sub-clustering of Fev+ hormone-negative endocrine cells from ref.29. CellRank’s predicted differentiation path visually aligns with the Fev+ delta sub-cluster. **c**. Delta fate probabilities within each Fev+ sub-cluster. Cells in the Fev+ delta sub-cluster are assigned significantly higher probability by CellRank (two-sided Welch’s t-test, 51 Fev+ Delta cells vs. 536 other Fev+ cells, t = 8.6, P = 1.7 x 10^-11^, Online Methods). **d.** Smoothed gene expression trends of the top 50 genes whose expression values correlate best with delta fate probabilities, sorted according to peak in pseudotime. Not all gene names are shown (see Supplementary Fig. 13a for full list). Right, UMAP projected MAGIC^85^ imputed expression values of *Hhex* and *Cd24a,* examples of known regulators which were identified automatically, as well as of *Hadh.* **e**. Smoothed gene expression trends along the delta lineage for all 12,987 genes which are expressed in at least ten cells, clustered using louvain^86^. Cluster 1 contains transiently upregulated genes. Solid line denotes mean trend, dashed lines denote 1 SD. Genes within cluster 1 are sorted according to their correlation with delta fate probabilities. Right, expression on the UMAP of *Map2k4, Msi1* and *Nefl,* among the genes that correlated best.

To discover more delta genes, we correlated the expression values of all genes against CellRank delta fate probabilities. Smoothed gene expression trends for the 50 genes with highest correlation revealed a cascade of gene activation events (Fig. 3d). Among the top 50 genes are *Hhex* and *Cd24,* as well as *Sst,* the hormone produced by mature delta cells^38^. Genes with no previously described role in delta cell differentiation include *Hadh* (a target of *Foxa2^46^*, implicated in pancreatic differentiation^47^), *Isl1* (a transcription factor involved in pancreatic differentiation^48–50^) and *Pkhd1* (a target of *Hnf1a/b^51–52^*, transcription factors involved in pancreatic differentiation^53^). Next, we focused on a cluster of transiently upregulated genes (Fig. 3e). When ranked by their correlation with delta fate, we identified *Map2k4, Msi1* and *Nefl* as novel candidate regulators. *Msi1* is regulated by *Rfx4^54^*, which is a paralog of *Rfx6* that is structurally related to *Rfx3^55^*, both of which are involved in endocrine differentiation^50,56–60^.

### Propagating uncertainty rectifies noise in RNA velocity

CellRank’s success with the delta cell fate is in part due to its ability to properly account for uncertainty in noisy RNA velocity vectors. Both the original velocyto and generalized scVelo models compute velocity vectors on the basis of spliced to unspliced count ratios^18,19^. However, these counts are influenced by many sources of biological and technical noise, such as ambient RNA, sparsity, doublets, bursting kinetics and low capture efficiency. Unspliced RNA in particular is rarer in the cell and suffers from low detection rates. The uncertainty in molecule counts translates into uncertainty in RNA velocity vectors, which can be estimated in scVelo (Fig. 4a and Online Methods). CellRank accounts for these sources of uncertainty by propagating the estimated distribution over velocity vectors (Fig. 4b,c). By default, it uses an analytical approximation which computes the expected value of the transition probabilities towards nearest neighbors, given the distribution over velocity vectors (Online Methods). The analytical approximation is very efficient and ensures that we can handle uncertainty even for large datasets. Alternatively, CellRank has an option for far slower, more accurate computation of fate probabilities via Monte Carlo (MC) sampling (Online methods). We confirm that this gives similar results to our analytical approximation (Supplementary Fig. 14a-c).

**Figure 4:**
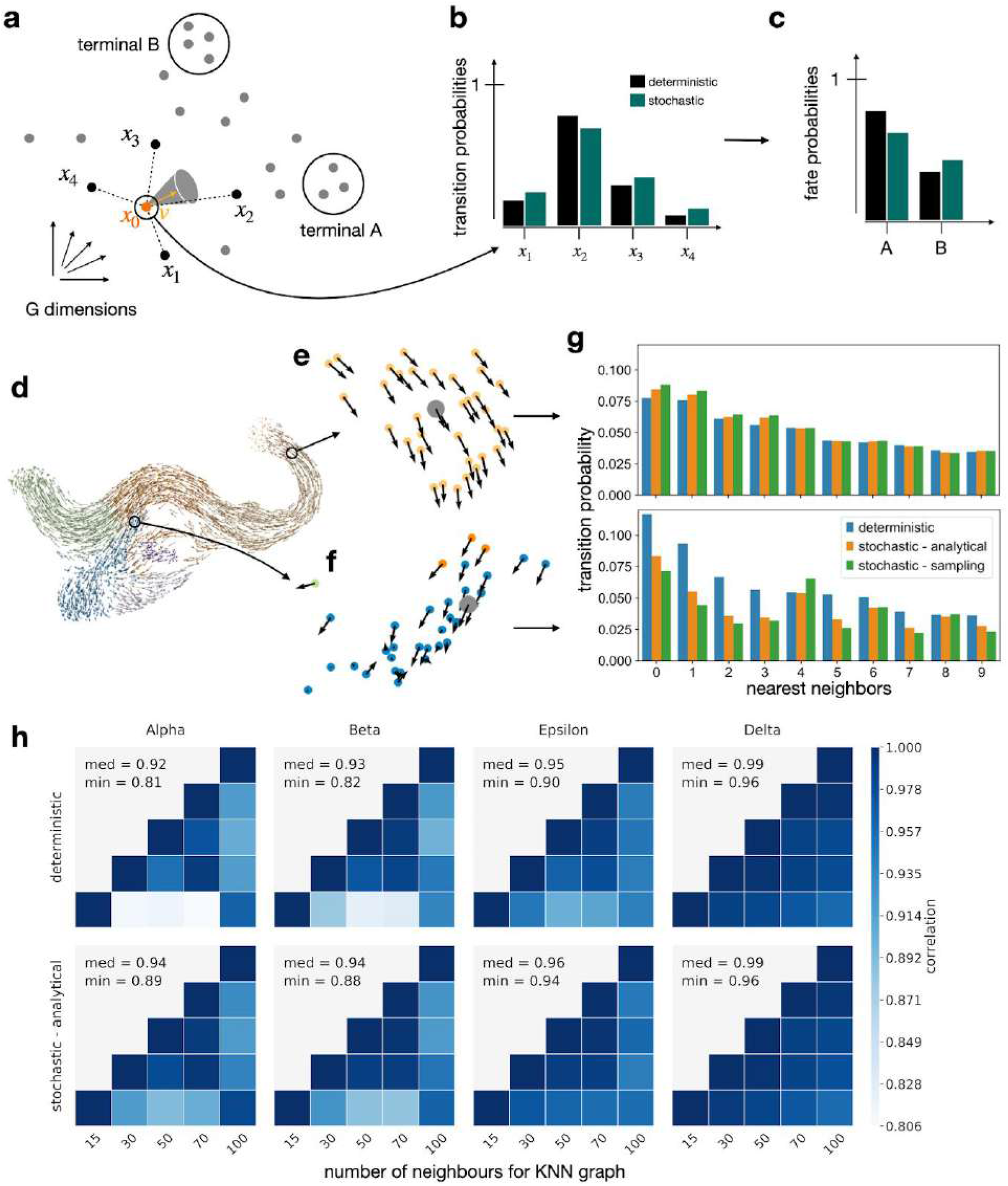
Uncertainty propagation adjusts for noise in RNA velocity vectors. **a.** When predicting the future state of cell *x*_0_, CellRank takes uncertainty in the velocity vector *v* in the high-dimensional gene space into account. **b**. Propagating noise changes the transition probabilities from one cell to its nearest neighbors. c. The adjusted transition probabilities agglomerate over longer paths to result in adjusted fate probabilities. **d-f.** Effect of noise propagation, illustrated using pancreas data. One cell from a low noise region, where velocity vectors from neighboring cells tend to point in the same direction (e), and one from a high noise region, where vectors from neighboring cells point in different directions (f), are highlighted. **g**. Transition probabilities from the reference cell to its 10 nearest neighbors using a deterministic or stochastic (analytical approximation or Monte Carlo sampling-based) formulation, for both the low and high noise cell. Corrections applied by stochastic approaches are larger in the high noise region. h. Propagating uncertainty increases robustness of fate probabilities with respect to parameter changes. This plot shows variation in the number of neighbors during KNN graph construction (see Supplementary Figs. 15 and 16 for more parameters).

We used the full pancreas dataset to investigate the effects of uncertainty propagation (Fig. 4d). We selected two cells, one from a low noise region where velocity vectors of neighboring cells tend to agree (Fig. 4e) and one from a high noise region (Fig. 4f). To compute transition probabilities towards nearest neighbors, we used a deterministic approach that does not propagate uncertainty, as well as our analytical approximation and MC sampling. Differences between deterministic and stochastic transition probabilities were greatest in the high noise region, highlighting that uncertainty propagation automatically down-weights transitions towards cells in noisy areas where individual velocity vectors are less trustworthy (Fig. 4h). Overall, this leads to increased robustness of fate probabilities (Fig. 4i and Supplementary Figs. 15a-e and 16a-e).

### CellRank outperforms competing similarity-based methods

To evaluate the impact of including velocity information, we compared CellRank with other methods that provide cell-fate probabilities: Palantir^25^, STEMNET^61^ and FateID^62^ on the pancreas data. Only CellRank correctly identified both initial and terminal states (Fig. 5a). Palantir requires user-provided initial states and only identified 2 out of 4 terminal states, and STEMNET and FateID cannot determine either initial or terminal states. Next, we supplied all methods with CellRank terminal states and tested cell fate probabilities, finding that only CellRank and Palantir correctly identified beta as the dominant fate among Ngn3^high^ EP cells (Fig. 5b). For gene expression, CellRank and Palantir correctly predicted trends for key lineage drivers, whereas FateID failed to predict (transient) upregulation of *Pdx1* and *Pax4* along the beta lineage^39,40^ as well as upregulation of *Arx* along the alpha lineage^39^, and STEMNET does not visualize expression trends (Fig. 5c and Supplementary Figs. 17a-c, 18a-f and 19a-f).

**Figure 5:**
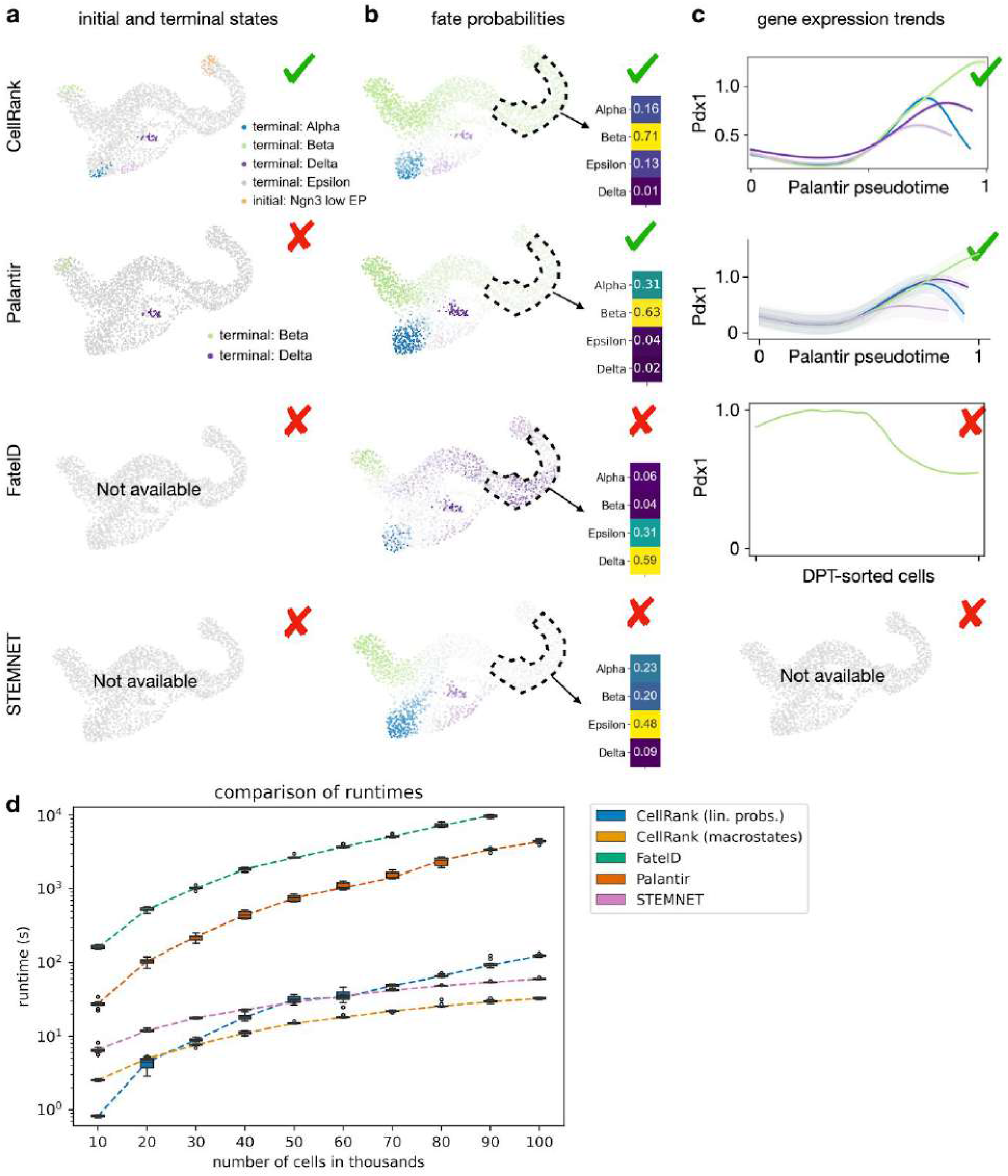
CellRank outperforms methods that do not include RNA velocity. (**a-c**) Methods were compared on pancreas data. **a.** Only CellRank correctly identifies initial and terminal states. **b.** CellRank and Palantir correctly predict beta to be the dominant fate among Ngn3^high^ EP cells. **c.** Gene expression trends for the beta-regulator *Pdx1^40,87,88^*. On the x-axis is the pseudotime used by the corresponding method, on the y-axis is gene expression. For FateID, the x-axis is given by the cell indices which are assigned to the beta lineage, sorted by diffusion pseudotime^10^ (DPT). We show one smoothed trend per lineage for CellRank and Palantir and a smoothed trend along just the beta lineage for FateID because it does not allow one gene to be visualised simultaneously along several lineages. CellRank and Palantir correctly identify upregulation of *Pdx1* along the beta lineage. FateID fails to do so while STEMNET does not offer an option to visualise gene expression trends (Online methods). **d.** Boxplot comparing methods in terms of computational runtime on a 100k cell reprogramming dataset^63^ (Online methods). We split the datasets into 10 subsets of increasing size and run each method 10 times on each subset. Box plots show the median, the box covers the 25 to 75% quantiles, whiskers extend up to 1.5 times the interquartile range above and below the box and the dashed lines connect the medians. Outliers are shown as dots. For CellRank, we separately recorded the time it takes to compute macrostates (and therewith terminal states) and fate probabilities.

We also benchmarked runtime and memory usage on an scRNA-seq dataset of 100k cells undergoing reprogramming from mouse embryonic fibroblasts to induced endoderm progenitor cells^63^ (Fig. 5d, Supplementary Fig. 20a and Online Methods). It took CellRank about 33 sec to compute macrostates from this large dataset (Supplementary Table 1). For fate probabilities, the (generalized) linear model STEMNET was fastest as expected, taking only 1 min, while CellRank took about 2 min and Palantir took 1h 12 min. FateID on 90k cells took even longer and failed on 100k cells due to memory constraints. For peak memory usage, results looked similar with CellRank requiring 3 respectively 5 times less memory than Palantir and FateID on 100k cells to compute fate probabilities (Supplementary Fig. 20a and Supplementary Table 2). Only STEMNET required even less memory. On 100k cells without parallelisation, CellRank had a peak memory usage of less than 23 GiB, making it possible to run such large cell numbers on a modern laptop (Suppl. Tab. 3).

### Fate probabilities predict a novel dedifferentiation trajectory in lung regeneration

To demonstrate CellRank’s ability to generalize beyond development, we applied it to murine lung regeneration in response to acute injury^64^. The scRNA-seq dataset comprised 24,051 lung airway and alveolar epithelial cells, sequenced at 13 time points spanning days 2-15 after bleomycin injury (Supplementary Fig. 21a,b). A high degree of plasticity between epithelial cell types has been observed when homeostasis is perturbed and the tissue environment changes^65^, including injury-induced reprogramming of differentiated cell types to bona fide long-lived stem cells in the lung^66^ and other organs^67^. In the current airway cell lineage model, multipotent basal cells give rise to club cells, which in turn can give rise to secretory goblet and ciliated cells^68^. Interestingly, it has been shown that upon ablation of basal stem cells, luminal secretory cells can dedifferentiate into fully functional basal stem cells^66^. Here, we applied CellRank for unbiased discovery of unexpected regeneration trajectories among airway cells.

We computed scVelo velocities and applied CellRank to identify nine macrostates (Fig. 6a and Supplementary Fig. 22a). Focusing our analysis on airway cells, we identified three macrostates in ciliated cells, one in basal cells and one in goblet cells. In agreement with lineage tracing experiments^69^, we observed a high probability for club cells to give rise to ciliated cells (Supplementary Fig. 22b-e). The goblet cell macrostate was distinguished from club cells by the expression of specific mucin genes such as *Muc5b* and *Muc5ac,* as well as secreted proteins involved in innate immunity, such as *Bpifb1* (Supplementary Fig. 21c). Analysis of fate probabilities towards basal and goblet states revealed that, surprisingly, goblet cells are likely to dedifferentiate towards Krt5+/Trp63+ basal cells (Fig. 6b,c and Supplementary Fig. 23a-d).

**Figure 6:**
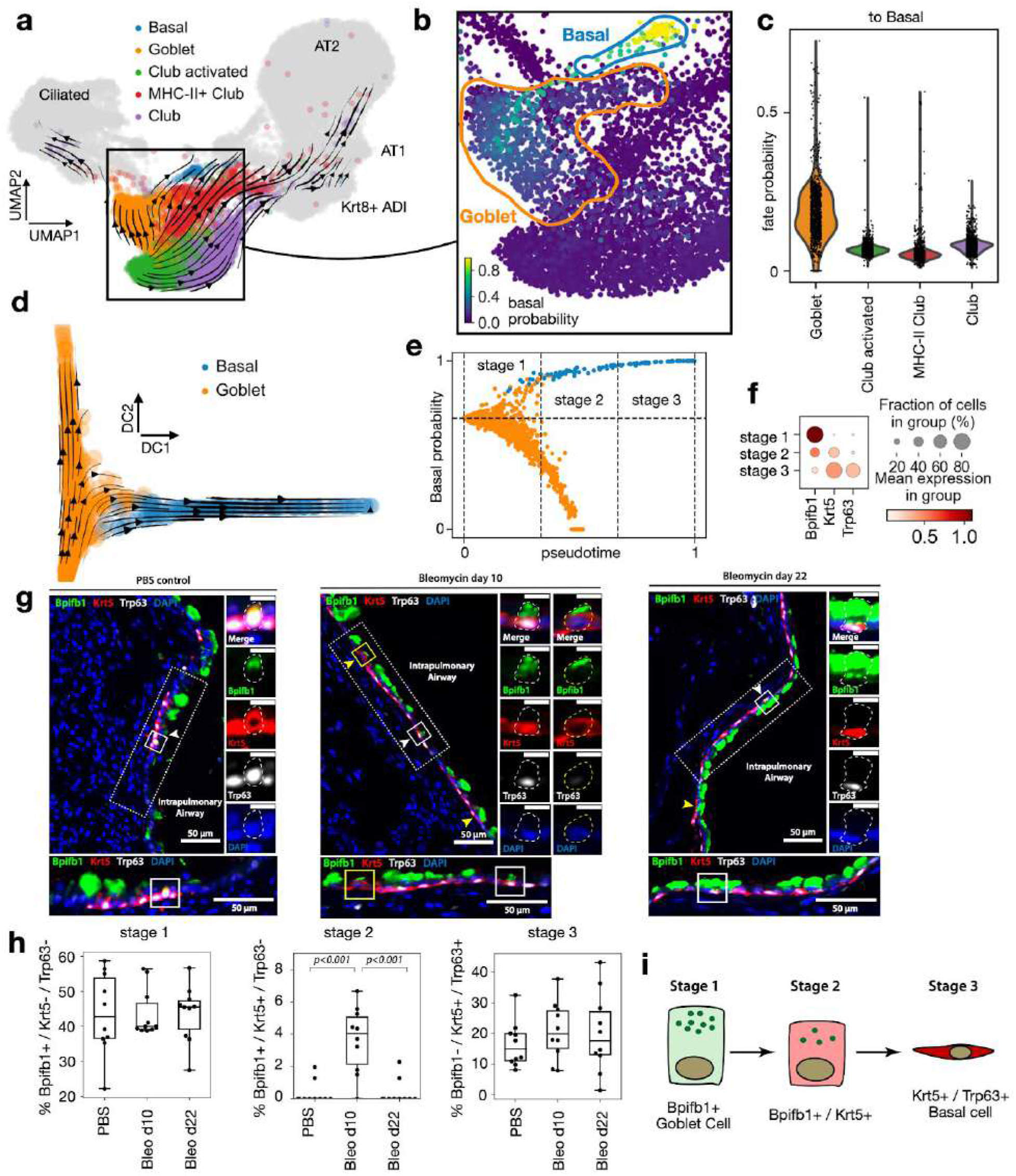
CellRank predicts a novel de-differentiation trajectory in murine lung regeneration. **a**. UMAP of 24,051 epithelial cells from 13 time points, spanning days 2-15 after lung injury by bleomycin treatment in mice. Airway cells from published clusters and annotations^64^ are highlighted. Streamlines show averaged and projected scVelo velocities. **b**. Cells in UMAP colored by CellRank-computed fate probabilities towards the basal cell macrostate, showing a route from goblet to basal cells **c**. Basal cell fate probabilities for secretory airway clusters, colored as in a. Goblet cells have significantly higher probability to transition towards the basal state (two-sided Welch’s t-test, 1116 Goblet cells vs. 4668 other secretory airway cells, t = 42.2, P = 1.4 x 10^-236^, Online methods). **d**. Diffusion map computed on the subset of goblet and basal cells, with projected streamline-aggregated scVelo velocity vectors. **e**. CellRank fate probabilities and Palantir pseudotime^25^ are used to define three stages of the dedifferentiation trajectory (Online methods) f. Dedifferentiation stages are characterized by expression of *Bpifb1* (goblet), *Krt5* (early basal) and *Trp63* (late basal); stage 1 corresponds to goblet, stage 2 to intermediate, and stage 3 to basal cells. **g**. Immunofluorescence stainings for Bpifb1 (green), Krt5 (red), Trp63 (white) and DAPI (blue) in mouse lung tissue sections. We find triple positive Bpifb1+/Krt5+/Trp63+ cells (white squares and arrow heads) and cells from the intermediate stage 2 (Bpfib1+/Krt5+/Trp63-) in bleomycin-injured lungs (yellow squares and arrow heads) and only very rarely in phosphate-buffered saline (PBS)-treated controls (Suppl. Fig. 25a). Scale bars = 50 μm, 10 μm for zoom-in images. In each panel, dotted-line boxes are magnified at bottom, and solid-line boxed cells are magnified at right, showing individual and merged channels. **h**. Quantification of cell abundances by stage in wild type (PBS, n=2), 10 days post bleomycin injury (bleo d10, n=2), and 22 days post injury (bleo d22, n=2) mice, in n = 5 intrapulmonary airway regions per mouse. Bleo d10 is significantly enriched for stage 2 cells (Nested One-Way ANOVA with Tukey’s multiple comparison test, P < 10^-3^) **i**. Proposed model for the dedifferentiation trajectory.

We computed a diffusion map^70^ on basal and goblet cells alone to study the trajectory at higher resolution (Fig. 6d). Using CellRank and the CGSD, we identified early cells in the transition, from which we computed a pseudotime using Palantir (Supplementary Fig. 24a-e). We combined pseudotime with the probability of transitioning towards the basal fate to define stages within the dedifferentiation trajectory in the data subset (Fig. 6e), splitting cells with at least 66% probability of reaching the basal state into three equal pseudotime bins. Stage 1 consists of goblet cells characterized by high expression of goblet marker *Bpifb1.* Stage 2 comprises an intermediate set of cells that express both *Bpifb1* and basal marker *Krt5.* Stage 3 consists of terminal basal cells, characterized by basal markers *Krt5* and *Trp63,* and no expression of *Bpifb1*.

Our novel goblet cell dedifferentiation model predicts that after injury, the frequency of stage 2 cells should increase. To validate this prediction, we assessed *Bpifb1, Krt5* and *Trp63* expression by immunofluorescence of mouse airway epithelial cells at days 10 and 21 post-bleomycin treatment, as well as in untreated animals (Fig. 6g). Cells from stage 1 (goblet) and stage 3 (basal) were found in both control and treated mice. However, intermediate stage 2 cells were only found in 10-day post-treatment mice (Fig. 6h,i). Furthermore we also found triple positive cells, which however only appeared after injury (Suppl. Fig. 25a). Goblet cell hyperplasia, an increase in the number of mucous secreting cells in the airways, is a prominent feature in several chronic inflammatory conditions^71^. The novel dedifferentiation trajectory to basal stem cells that CellRank analysis discovered is unexpected, suggesting a route for generating multipotent stem cells in the resolution phase of the regenerative response to injury.

## Discussion

We have shown that CellRank combines gene-expression similarity with RNA velocity to robustly estimate directed trajectories of cells in development and regeneration. Applied to pancreatic development, CellRank correctly recovered initial and terminal states, fate potentials and gene expression trends, outperforming competing methods that do not use RNA velocity information, while only taking seconds to compute terminal states and a few minutes to compute fate potentials on 100k cells.

Although we identified alpha, beta and epsilon states automatically, the delta macrostate required us to manually assign its terminal status. We believe that the cutoff for terminal assignment was not reached because delta cells are rare in this dataset and their regulation is not detected correctly by velocities. To overcome this problem, it may be possible to extend the CellRank model to include epigenetic information such as chromatin accessibility. Many regulatory processes are initiated at the epigenetic level and only become visible transcriptionally after a delay, or not at all^3,72,73^. Including such information in the CellRank model, possibly by introducing limited memory to the Markov chain, could therefore increase its applicability.

For delta cell development, we showed how clustering gene expression trends within one lineage and correlating with fate probabilities can identify putative driver genes. Alternatively, drivers could be identified directly through statistical tests on the parameters of the generalized additive models used for fitting these trends. Such models already exist and could benefit from CellRank’s fate probabilities for assigning cells to lineages^74,75^.

RNA velocity vectors are noisy estimates of the current state of gene regulation. CellRank takes care of uncertain velocity vectors by propagating their distribution. We showed how this correction automatically scales with the local noise level and increases robustness. A current limitation is that we need to approximate the distribution over velocity vectors by computing moments over velocity vectors in the local neighborhood. In the future, we envisage an end-to-end framework where uncertainty is propagated all the way from raw spliced and unspliced counts via velocities into the final quantities of interest, i.e. initial and terminal state assignments as well as fate probabilities.

CellRank is a method to quantitatively analyze RNA velocity-induced vector fields in high dimensions. While other approaches have started to address this, they either ignore the stochastic nature of cellular fate decisions and velocity uncertainty^76,77^, or they focus on questions other than trajectory reconstruction^78^. The original velocyto^18^ model proposed an idea to find initial and terminal states that was based on simulating a Markov process forwards or backwards in time, however, their implementation relied on a 2D t-SNE embedding, was computationally expensive because of the sampling, ignored uncertainty in velocity vectors and did not allow the separation into individual initial and terminal states. CellRank overcomes these problems through a principled approach that is simulation free, independent of any low-dimensional embedding, takes into account velocity uncertainty and is able to identify individual initial and terminal states.

In contrast to previous Markov-chain based methods^10,70^, our approach is based on a directed non-symmetric transition matrix. This implies that straight-forward eigendecomposition of the transition matrix to learn about aggregate dynamics is not possible, as eigenvectors of non-symmetric transition matrices are in general complex and do not permit a physical interpretation. To overcome this, one possibility would be to revert to computationally expensive simulation-based approaches^79^. In CellRank, we took a more principled approach based on the real Schur decomposition, a generalization of the eigendecomposition to non-diagonalizable matrices. A current limitation of our implementation is that it uses a derivative-free method^80,81^ to solve the resulting constrained optimization problem, which may get computationally expensive as the number of macrostates increases. A possible solution we would like to explore is to use a Gauss-Newton-based optimizer^24^.

Similarity-based methods have shown great success, but their application has been mostly limited to normal development, because the starting cell and direction of the process are often established. Here, we show how CellRank generalizes beyond normal development, by discovering a novel goblet to basal cell dedifferentiation trajectory upon lung injury. The dataset consists of 13 time points^64^; however, it is unclear how to incorporate temporal information into the estimation of transition probabilities. Including this information could help to regularize the model, by only allowing transitions consistent with experimental time^82^. Further, where available, lineage tracing information could be included to further regularize the model to obey clonal dynamics^83^.

CellRank could be easily applied to data from metabolic labeling^6–9^. As a general framework for interpreting high dimensional vector fields, we anticipate that CellRank will be useful to describe complex trajectories in regeneration, reprogramming and cancer, where determining the direction of the process is often challenging.

## Acknowledgements

We thank S. Tritschler for helping us with the biological interpretation of results, F. Paul for guidance regarding PCCA and GPCCA, G. Diez for pointing us to literature in the conformational protein dynamics field, J. L. R. López for his input to the implementation and T. Walzthoeni (Bioinformatics Core Facility, Institute of Computational Biology, Helmholtz Zentrum München) for bioinformatics support. We would further like to thank M. Weber for valuable discussions regarding irreversible Markov processes, M. Weber and A. Sikorski for pointing us to SLEPc for partial Schur decompositions of sparse matrices, J. Chan, G. Palla and L. Hubert for stimulating discussions and T. Nawy for helping to write the manuscript. This work was supported by the BMBF (grant# 01IS18036B and grant# 01IS18053A) and by the Helmholtz Association (Incubator grant sparse2big, grant # ZT-I-0007). M. Lange further acknowledges financial support by the DFG through the Graduate School of QBM (GSC 1006), by the Joachim Herz Stiftung and by the Bayer Foundations.

## Author’s contributions

ML designed and developed the method, implemented CellRank, analyzed the data and wrote the online methods. MK contributed to the implementation and to the data analysis. BR developed the original GPCCA code, integrated it into the MSMTools package and helped to integrate it into CellRank. VB contributed to the implementation and helped to harmonize CellRank with scVelo and SCANPY. MS helped with the design of the method and the presentation of results. MB and HL interpreted the relevance of the method for inferring developmental trajectories in the pancreas data. HS and JS interpreted the relevance of the method for inferring regeneration trajectories in the lung and experimentally validated the dedifferentiation hypothesis. FJT and DP supervised the project and contributed to the conception of the project. ML, FJT and DP wrote the manuscript with contributions from the coauthors. All authors read and approved the final manuscript.

## Data availability

Raw published data for the pancreas^29^, lung^64^ and reprogramming^63^ examples is available from the Gene Expression Omnibus (GEO) under accession codes GSE132188, GSE141259 and GSE99915, respectively. Processed data, including spliced and unspliced count abundances, is available from figshare under https://doi.org/10.6084/m9.figshare.c.5172299.

## Code availability

The CellRank software package is available at https://cellrank.org including documentation, tutorials and examples. Jupyter notebooks to reproduce our analysis and figures are available at https://github.com/theislab/cellrank_reproducibility.

## Supplementary data

Supplementary figures: Supplementary Figs. 1 – 25

Supplementary tables: Supplementary Tables 1 – 3

## Ethics statement

Pathogen-free female C57BL/6J mice were purchased from Charles River Germany and maintained at the appropriate biosafety level at constant temperature and humidity with a 12h light cycle. Animals were allowed food and water ad libitum. All animal experiments were performed in accordance with the governmental and international guidelines and ethical oversight by the local government for the administrative region of Upper Bavaria (Germany), registered under 55.2-1-54-2532-130-2014 and ROB-55.2-2532.Vet_02-16-208.

## CellRank – Online Methods

### 1 The CellRank algorithm

The aim of the CellRank algorithm is to detect the initial, terminal and intermediate states of a cellular system and to define a global map of fate potentials that assigns each cell to these states in a probabilistic manner. Given our inferred fate potentials, we compute gene expression trends along trajectories in the fate map and provide several possibilities for visualizing these. The inputs to CellRank are a count matrix *X* ∈ ℝ^*N×G*^ where *N* is the number of cells and *G* is the number of genes as well as a velocity matrix *V* = ℝ^*N×G*^, defining a vector field representing RNA velocity^1,2^ for each cell and gene. Note that CellRank can be generalized to any kind of vector field, i.e. *V* could equally represent directed information given by e.g. metabolic labeling^3–6^. There are three main steps to the CellRank algorithm:

1. Compute transition probabilities among observed cells. These reflect how likely a cell with a given cell state, defined by its gene expression profile, is to change its profile to that of a target cell. We compute these probabilities by integrating two sources of evidence: (1) transcriptomic similarity between the source and target cells and (2) an extrapolation of a cell’s current gene expression profile into the near future using RNA velocity. We aggregate these transition probabilities in the transition matrix *P* and use it to model cell-state transitions as a Markov chain.
2. Coarse-grain the Markov chain into a set of initial, terminal and intermediate macrostates of cellular dynamics. Each cell is assigned to each macrostate via a membership matrix *χ*. The assignment is soft, i.e. each cell has a certain degree of confidence of belonging to each macrostate. We compute transition probabilities among macrostates in the matrix *P_c_*. This matrix allows us to identify whether macrostates are initial, terminal or intermediate.
3. Compute fate probabilities towards a subset of the macrostates. This will typically include the terminal states, but can also include intermediate states, depending on the biological question. We compute how likely each cell is to transition into each of the selected macrostates and return these probabilities in a fate matrix *F*.

#### CellRank extracts the essence of cellular state transitions

The principle of the CellRank algorithm is to decompose the dynamics of the biological system into a set of dynamical macrostates. We target macrostates that are associated with regions in the phenotypic manifold which cells are unlikely to leave once they have entered them. For each observed cell, we compute how likely it is to belong to each of these macrostates. We accumulate these soft assignments in a membership matrix *χ* ∈ ℝ^*N×n*_*s*_^. Further, we compute a coarse-grained transition matrix *P_c_* ∈ ℝ*^n_s_×n_s_^* which specifies transition probabilities among macrostates. The coarse-grained transition matrix allows us to reduce the biological system to its essence: dynamical macrostates of observed cell-state transitions and their relationship to one another. Based on the coarse-grained transition matrix, we classify macrostates as either initial, intermediate or terminal. Initial states will be macrostates that have very small incoming but large outgoing transition probability. Intermediate states will be macrostates that have both incoming and outgoing transition probability. Terminal states will be macrostates that have large incoming but very little outgoing and large self-transition probability.

#### CellRank computes probabilistic fate potentials

Each macrostate is associated with a subset of the observed cells via the membership matrix *χ*. Once we classified macrostates as either initial, intermediate or terminal using the coarse-grained transition matrix *P_c_*, we may ask how likely each cell is to transition to each of the *n_t_* terminal states. CellRank efficiently computes these probabilities and returns a fate matrix *F* ∈ ℝ^*N×n_t_*^. The matrix *F* extends the short-range fate relationships given by RNA velocity to the global scale: from initial to terminal states along the phenotypic manifold. We account for high noise levels in the velocity vectors via a stochastic Markov chain formulation, by restricting predicted transitions to align with the phenotypic manifold and by propagating velocity uncertainty into the Markov chain.

#### CellRank uncovers gene expression trends towards specific terminal populations

The outputs of the CellRank algorithm are

- a membership matrix *χ* ∈ ℝ^*N×n_s_*^ where *n_s_* is the number of macrostates. Row *i* in *χ* softly assigns cell *i* to any of the macrostates.
- a coarse-grained transition matrix *P_c_* ∈ ℝ^*n_s_*×*n_s_*^ that describes how likely these macrostates are to transition into one another. The matrix *P_c_* allows macrostates to be classified as either initial, intermediate or terminal.
- a fate matrix *F* ∈ ℝ^*N*×*n_t_*^ where *n_t_* is the number of terminal states. Row *i* in *F* specifies how likely cell *i* is to transition towards any of the terminal states.

We use the fate matrix *F* to model gradual lineage commitment. Fate biases can be aggregated to the cluster level and visualized as pie charts on a new directed version of PAGA graphs^7^ (Section 2). Further, we use the fate matrix *F* to uncover gene expression trends towards the identified terminal states (Section 3). Once the trends have been fit, they can be clustered to discover the main regulatory dynamics towards different terminal states (Section 4). For the identification of putative regulators towards specific terminal states, we correlate gene expression values with fate probabilities (Section 5).

##### 1.1 Modelling approach

Similarly to other methods^8–10^, CellRank models cell state transitions among observed cellular profiles. Unlike other velocity based methods, following the success of pseudotime methods, key to our model is that we restrict possible state changes to those consistent with the global structure of the phenotypic manifold via a KNN graph computed based on similarities in gene expression space. Our approach then biases the likely future state of an observed cell within its local graph neighborhood based on RNA velocity, by combining transcriptional similarity with RNA velocity to direct edges in the graph and to assign a probability to each cell state transition. When computing these probabilities, we take into account uncertainty in the velocity vectors. By aggregating individual, stochastic transitions within the global structure of the phenotypic manifold, we uncover the fate bias for individual cells. We make the following assumptions:

- state transitions are gradual, daughter cells are in general transcriptomically similar to their mother cells. Cells traverse a low-dimensional phenotypic manifold from initial to terminal states via a set of intermediate states.
- the set of sampled cellular profiles spans the entire state change trajectory, i.e. intermediate states have been covered, there are no ‘gaps’ in the trajectory.
- while for an individual cell, its past history is stored in epigenetic modifications, we model average cellular dynamics where state transitions occur without memory.
- RNA velocity approximates the first derivative of gene expression. This must not precisely hold for every gene in each individual cell as we treat state transitions as a stochastic process, enforce alignment with the manifold and propagate uncertainty, but it should hold in expectation for enough cells so that we are able to estimate the overall directional flow.

Based on these assumptions, we model cellular state transitions using a Markov chain: a stochastic process *X* = (*X_t_*)_*t∈T*_ – a sequence of random variables *X_t_*: *Ω* → *E* on a probability space 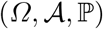 over a countable set *Ω* mapping to a measurable state space (*E, Σ*) – that describes the evolution of a probability distribution over time where the future distribution only depends on the current distribution and not on the past, i.e. Pr(*X*_*t*_n+1__ = *x* |*X*_*t*_1__ = *x*|, *X*_*t*_2__ = *x*_2_, …,*X_t_n__* = *x_n_*) = Pr(*X*_*t*_*n*+1__ = *x* | *X_tn_* = *x_n_*). We use a Markov chain over a discrete and finite state space *Ω*, where each state in the chain is given by an observed cellular transcriptional profile. To define the Markov chain, we need to compute a transition matrix *P* ∈ ℝ^*N×N*^ which describes how likely one cell is to transition into another. We construct *P* ∈ ℝ^*N×N*^ using a KNN graph based on transcriptional similarity between cells and a given vector field. While CellRank generalizes to any given vector field, we demonstrate it using RNA-velocities, based on unspliced to spliced read ratios, computed with scVelo^9^.

#### Defining initial, intermediate and terminal states in biological terms

We define an initial (terminal) state as an ensemble of measured gene expression profiles which, when taken together, characterize the starting (end) point of one particular cell-state change. We define an intermediate state as an ensemble of gene expression profiles which, when taken together, characterize a point on the cell-state transition trajectory which lies in between one or several initial and terminal states.

#### Translating initial, intermediate and terminal states into mathematical terms

To translate the above terms into mathematics, we make use of the coarse-graining given by the membership matrix *χ* and the coarse-grained transition matrix *P_c_*. We show below that our assignment of cells to macrostates maximizes a quantity called *crispness*^11^: we obtain macrostates which have little overlap and large self-transition probability. In other words: we recover the kinetics of the Markov chain on slow-time scales, i.e. macrostates and their transitions reflect the limiting behavior of the Markov chain. Among the set of macrostates, we identify initial states as those which have little incoming large but large outgoing transition probability in *P_c_*. Intermediate states will have both incoming and outgoing transition probability in *P_c_*. Terminal states will have large incoming but little outgoing and large self-transition probability in *P_c_*. An important term in the mathematical framework is *metastability*: a process starting in a metastable state will stay there with high probability for a long time. Accordingly, we define a metastable state of cellular dynamics as an area in phenotypic space that cells are unlikely to leave again once they have entered. A metastable state will typically correspond to a terminal state, while an intermediate state is typically only weakly metastable. Initial states can constitute weakly metastable states, if the probability of leaving them is small, potentially because of heavily cycling populations.

#### Reversing the Markov chain to recover initial states

Initial states may not be picked up as macrostates during coarse-graining of the Markov chain because they are not stable enough, i.e. cells in the initial state have very little probability of transitioning into one another and rapidly start traversing their state change trajectory. In these cases, we reverse the Markov chain, i.e. we flip the arrows in the velocity vector field *V*. The initial state now constitutes a terminal (i.e. metastable) state of the reversed dynamics and may be recovered by coarse-graining and interpreting the reversed Markov chain.

#### Defining fate probabilities towards macrostates

Biologically, we define the fate probability of cell *i* to reach macrostate *j* ∈ 1,…,*M* as the probability of cell *i* executing a series of gene expression programs which change its phenotype to match the phenotype of cells in macrostate *j*. Within the context of fate probabilities, we will typically be interested in macrostates which are either terminal or intermediate states. Mathematically, we translate this to the probability of a random walk on the Markov chain initialized in cell *i* to reach any cell belonging to macrostate *j* before reaching any cell belonging to another macrostate. CellRank efficiently computes these probabilities in closed form using absorption probabilities (Subsection 1.4).

##### 1.2 Computing the transition matrix

We model each observed cell by one microstate in the Markov chain. To compute transition probabilities among cells, we make use of transcriptomic similarity to define the global topology of the phenotypic manifold and of RNA velocity to direct local movement on the manifold. To model the global topology of the phenotypic manifold, the first step of the CellRank algorithm is to compute a KNN graph.

#### Computing a KNN graph to align local transitions with global topology

We compute a KNN graph to constrain the set of possible transitions to those that are consistent with the global topology of the phenotypic manifold: Each cell is thus only allowed to transition into one of its *K* nearest neighbors. While CellRank can generalize to any reasonable similarity kernel, here, we compute the KNN graph as follows:

1. project the data onto the first *L* principal components to obtain a matrix *X_pc_* ∈ ℝ^*N×L*^, where rows correspond to cells and columns correspond to PC features.
2. for each cell *i*, compute distances to its *K*-nearest neighbors based on euclidean distance in *X_PC_*. Accumulate distances in a matrix *D* ∈ ℝ^*N×N*^.
3. the KNN relationship will lead to a directed graph because it is not a symmetric relationship. Symmetrize the KNN relations encoded by *D*, such that cells *i* and *j* are nearest neighbors if either *i* is a nearest neighbors of *j*, or *j* is a nearest neighbors of *i*. This will yield an undirected symmetric version *D_sym_* of *D*, where each cell has at least *K* nearest neighbors.
4. compute a symmetric adjacency matrix *A* based on *D_sym_* containing similarity estimates between neighboring cells according to the manifold structure. To approximate cell similarities, we use the method implemented in the UMAP algorithm, which adapts the singular set and geometric realization functors from algebraic topology to work in the context of metric spaces and fuzzy simplicical sets^12,13^.

We choose *K* = 30 to be the number of nearest neighbors by default. We show in Supplementary Fig. 8a that CellRank is robust to the choice of *K*. To compute the similarity metric, the option presented is the default in SCANPY^14^. Alternatively, similarity may be computed using a Gaussian kernel with density-scaled kernel width as introduced by ref.^15^ and adapted to the single cell context in ref.^10^. We choose *L* = 30 to be the number of principal components by default. This can be adapted based on knee-point heuristics or the percentage of variance explained, however, we show in Supplementary Fig. 8d that CellRank is robust to the exact choice of *L*.

#### Directing the KNN graph based on RNA Velocity

Next, we direct the edges of the KNN graph using RNA velocity information, giving higher probability to those neighbors whose direction best aligns with the direction of the velocity vector. Specifically, for cell *i* with gene expression profile *x_i_* ∈ ℝ^*G*^ and velocity vector *V_i_* ∈ ℝ^*G*^, consider its neighbors *j* = 1,2,…,*K_i_* with gene expression profiles {*x*_1_, *x*_2_, …,*x_K_*}. Note that the graph construction outlined above leads to a symmetric KNN graph, where *K_i_* is not constant across all cells, but *K_i_* ≥ *K*∀*i* ∈ {1, …,*N*}. For each neighboring cell *k*, compute the corresponding state-change vector with cell *i, s_ik_* = *x_k_* — *x_i_* ∈ *R*^*G*^. Next, we compute Pearson correlations *c_i_* ∈ *R*^*K*^ of *v_i_* with all state change vectors via

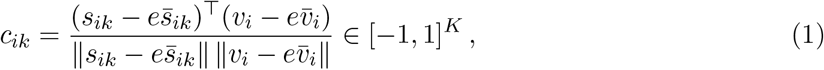

where *e* is a constant vector of ones and 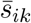 and 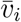 are averages over the state change vector and the velocity vector, respectively. Intuitively, *c_i_* contains the cosines of the angles that the mean-centered *v_i_* forms with the mean-centered state-change vectors *s_ik_*. A value of 1 means perfect correlation between the gene expression changes predicted by the local velocity vector and the actual change observed when going from the reference cell to any of its nearest neighbors. Pearson correlations have been computed in similar ways by scvelo^9^ and velocyto^1^ to project the velocity vectors into a given embedding. In Subsection 1.5 below, we show how their ideas can be formalized and extended to account for uncertainty in the velocity vector.

#### Transforming correlations into transition probabilities

To use the vector *c_i_* as a set of transition probabilities to neighboring cells, we need to make sure it is positive and sums to one. For cell *i*, define a set of transition probabilities *p_i_* ∈ ℝ^*K*^ via

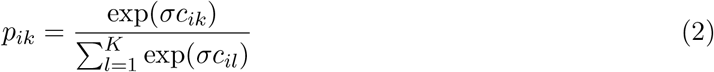

where *σ* > 0 is a scalar constant that controls how centered the categorical distribution will be around the most likely value, i.e. around the state-change transition with maximum correlation (see below). We repeat this for all (*i,k*) which are nearest neighbors to compute the transition matrix *P_v_* ∈ ℝ^*N×N*^. This scales linearly in the number of cells *N*, the number of nearest neighbors *K* and the number of genes *G* as the KNN graph is sparse.

#### Automatically determine

*σ* We reasoned that the value of *σ* should depend on typical Pearson correlation’s between velocity vectors and state change vectors observed in the given data-set. For this reason, we use the following heuristic:

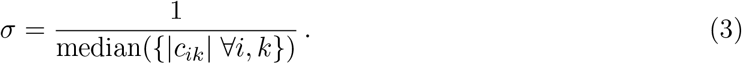

This means that if the median absolute Pearson correlation observed in the data is large (small), we use a small (large) value for σ. The intuition behind this is that for sparsely sampled data-sets where velocity vectors only roughly point into the direction of neighboring cells, we upscale all correlations a bit. Typical values for σ we compute this way range from 1.5 (lung example^16^) to 3.8 (pancreas example^17^).

#### Coping with uncertainty in the velocity vectors

scRNA-seq data is a noisy measurement of the underlying gene expression state of individual cells. RNA velocity is computed on the basis of these noisy measurements and is therefore itself a substantially noisy quantity. In particular the unspliced reads required by velocity and scVelo to estimate velocities are very sparse and their abundance varies depending on the amount of relevant intronic sequence of different genes. Besides this inherent noise, preprocessing decisions in the alignment pipeline of spliced and unspliced reads have been shown to impact the final velocity estimate^18^. Further uncertainty in the velocity estimate arises because assumptions have to be made which may not always be satisfied in practice:

- the original velocyto^1^ model assumes that for each gene, a steady state is captured in the data. The scVelo^9^ model circumvents this assumption by dynamic modeling, extending RNA velocity to transient cell populations, however there is often only a sparsity of transitional cells to estimate these dynamics.
- both models assume that the key biological driver genes for a given cell-state transition are intron rich and may therefore be used to estimate spliced to unspliced ratios. This has been shown to be the case in many neurological settings, however, in other systems such as hematopoeisis, it remains unclear whether this assumption is met.
- both models assume that per gene, a single set of kinetic parameters *α* (transcription rate), *β* (splicing rate) and *γ* (degratation rate) may be used across all cells. However, we know that in many settings, this assumption is violated because of alternative splicing or cell-type specific regulation^19–22^.
- both models assume that there are no batch effects present in the data. To date, to the best of our knowledge, there are no computational tools to correct for batch effects in velocity estimates.
- both models assume that the cell-state transition captured in the data is compatible with the time scale of splicing kinetics. However, this is often not known a priori and may explain the limited success of RNA velocity in studying hematopoeisis to date.

The points outlined above highlight that RNA velocity is a noisy, uncertain estimate of the likely direction of the future cell state. To cope with the uncertainty present in RNA velocity, we adapt four strategies:

- we restrict the set of possible transitions to those consistent with the global topology of the phenotypic manifold as described by the KNN graph.
- we use a stochastic formulation based on Markov chains to describe cell-state transitions. For cell *i* with velocity vector *v_i_*, we allow transitions to each nearest neighbor *j* with transition probability *p_j_*. This means that we even allow transitions backwards, against the flow prescribed by the velocity vector field, with small probability. This reflects our uncertainty in *v_i_*.
- we combine RNA velocity information with trancriptomic similarity, see below.
- we propagate uncertainty in *v_i_* into the downstream computations (Subsection 1.5).

#### Emphasizing transcriptomic similarity

Thus far, we have combined RNA velocity with transcriptomic similarity by computing a similarity-based KNN graph to restrict the set of possible transitions. To further take advantage of the information captured by the KNN graph and to increase robustness of the algorithm with respect to noisy velocity vectors, we combine the velocity based transition matrix *P_v_* with a similarity based transition matrix *P_s_* via

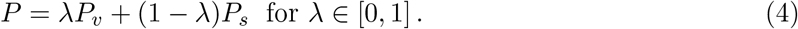

The matrix *P_s_* is computed by row-normalizing the adjacency matrix *A*. In practical applications, we have found that using values around λ = 0.2 increase robustness with respect to noisy velocity estimates. The matrix *P* is the final transition matrix estimated by the CellRank algorithm.

##### 1.3 Coarse-graining the Markov chain

The transition matrix *P* defines a Markov chain among the set of all observed cells, where each cell constitutes a microstate of the Markov chain. However, it is difficult to directly use *P* to interpret the cellular trajectory because *P* is a fine-grained, noisy representation of cell state transitions. Therefore, we seek to reduce *P* to its essence: macrostates representing key biological states and their transition probabilities among each other. We accomplish this using GPCCA^23^, which utilizes the *Generalized Perron Cluster Cluster Analysis* (G-PCCA)^24–26^, a method originally developed to study conformational dynamics in proteins. We adapt it to the single cell setting and utilize it to project the large transition matrix *P* onto a much smaller coarse-grained transition matrix *P_c_* that describes transitions among a set of macrostates. A macrostate is associated with a subset M of the state space *M* ⊂ *Ω*. The macrostates are defined through a so-called membership matrix *χ*. Rows of *χ* contain the soft assignment of cells to macrostates.

#### Generalized Perron Cluster Cluster Analysis (G-PCCA)

The aim of the G-PCCA method is to project the large transition matrix *P* onto a much smaller coarse-grained transition matrix *P_c_*, which describes transitions between macrostates of the biological system^24,25^. For the projected or embedded dynamics to be Markovian, we require the projection to be based on an invariant subspace of *P* (invariant subspace projection), i.e. a subspace *W* for which

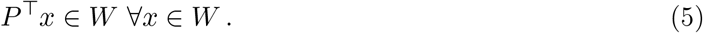

In case of a reversible *P*, invariant subspaces are spanned by the eigenvectors of *P*^11^. In our case however, *P* is non-reversible and the eigenvectors will in general be complex. Since the G-PCCA method can not cope with complex vectors, we rely on real invariant subspaces of the matrix *P* for the projection. Such subspaces are provided by the real Schur decomposition of *P*^24,25,27^,

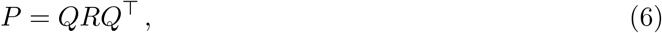

where *Q* ∈ ℝ^*N×N*^ is orthogonal and *R* ∈ ℝ^*N×N*^ is quasi-upper triangular^28^. *R* has 1-by-1 or 2-by-2 blocks on the diagonal, where the former are given by the real eigenvalues and the latter are associated with pairs of complex conjugate eigenvalues.

#### Invariant subspaces of the transition matrix

Columns of *Q* corresponding to real eigenvalues span real invariant subspaces. Columns of *Q* corresponding to pairs of complex conjugate eigenvalues span real invariant subspaces when kept together, but not if they are separated. Particularly, for columns *q_j_* and *q_k_* of *Q* belonging to a pair of complex conjugate eigenvalues, the space *W*_0_ = span(*q_j_, q_k_*) is invariant under P, but the individual *q_j_* and *q_k_* are not^29^. Depending on the constructed subspace, different dynamical properties of *P* will be projected onto *P_c_*. Choosing Schur vectors belonging to real eigenvalues close to 1, metastabilities are recovered, while for Schur vectors with complex eigenvalues close to the unit circle, cyclic dynamics are recovered^24,25^. Both options are available in CellRank, defaulting to the recovery of metastabilities.

#### Projecting the transition matrix

Let 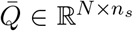 be the matrix formed by selecting *n_s_* columns from *Q* according to some criterion (metastability or cyclicity). Let *χ* ∈ ℝ^*N×n_s_*^ be a matrix obtained via linear combinations of the columns in 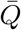, i.e.

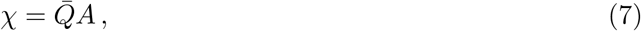

for an invertible matrix *A* ∈ ℝ^*n_s_×n_s_*^. We obtain the projected transition matrix via a Galerkin projection^24,25^,

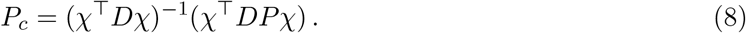

Here, the matrix *D* is the diagonal matrix of a weighted scalar product. The Schur vectors must be orthogonal with respect to this weighted scalar product, i.e. *Q*^T^*DQ = I* with the *n_s_*-dimensional unit matrix *I*, to yield the required invariant subspace projection. The diagonal elements of *D* are in principle arbitrary, but a convenient choice would be the uniform distribution or some distribution of the cellular states of interest. Choosing the uniform distribution, as is the default in CellRank, would result in a indiscriminate handling (without imposing any presumptions about their distribution) of the cellular states. Note that the matrix inversion in Equation (8) is performed on a very small matrix of size *n_s_ × n_s_*.

#### Computing the membership vectors

In principle, we could use any invertible *A* in Equation (7). However, we would like to interpret the rows of of *χ* as membership vectors that assign cells to macrostates. For this reason, we seek a matrix *A* that minimizes the overlap between the membership vectors *χ*, i.e. a matrix *A* that minimizes off-diagonal entries in *χ*^T^*D_χ_*. This is equivalent to maximizing

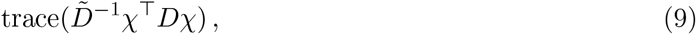

where 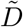 is chosen to row-normalize 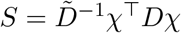,

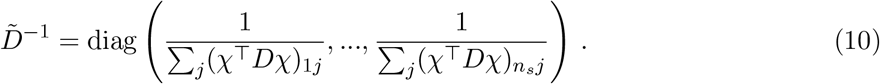

Choosing Schur vectors with real eigenvalues close to one, thus recovering metastability, maximizing Equation (9) can be interpreted as maximizing the metastability of the macrostates in the system. In practice, we use

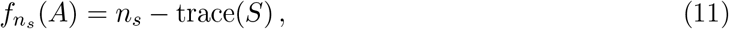

as our objective function, which is bounded below by zero and convex on the feasible set defined through the linear constraints^11^. We must minimize *f_n_s__* with respect to the constraints

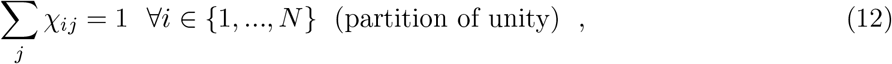

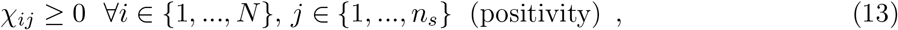

which is not a trivial task. Among the several possibilities to solve the minimization problem, a convenient choice is to perform unconstrained optimization on *A*(2: *n_s_*, 2: *n_s_*) using a trick: to impose the constraints after each iteration step, thus transforming the (unfeasible) solution into a feasible solution^11^. The drawback of this approach is that this routine is non-differentiable. Thus a derivative-free method like the Nelder-Mead method, as implemented in the Scipy routine `scipy.optimize.fmin`, should be used for the optimization.

#### Positivity of the projected transition matrix

Note that the projected transition matrix Pc may have negative elements if macrostates share a large overlap. In practice, this is caused by a subop-timal number of macrostates ns and can be resolved by changing that number. We may interpret Pc as the transition matrix of a Markov chain between the set of macrostates if it is non-negative within numerical precision^24^.

#### Tuning the number of macrostates

The number of macrostates ns can be chosen in a number of different ways:

- using the eigengap heuristic for the real part of the eigenvalues close to one.
- define the crispness *ξ* of the solution as the value of trace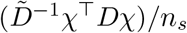^11^. The larger this value, the smaller the overlap between the macrostates, and in turn, the sharper or “crisper” the recovered macrostates. The crispness can be computed for different numbers of macrostates *n_s_* and the number *n_s_* with the largest value of *ξ* should be selected.
- to avoid having to solve the full problem for too many values of *n_s_*, do a pre-selection using the minChi criterion^11^: Based on an initial guess for *A*, compute a membership matrix *χ* and calculate minChi = min_*i,j*_ (*χ_ij_*). In general, this value will be negative because the starting guess is infeasible. The closer to zero the value of minChi is, the more we can expect *n_s_* to yield a crisp decomposition of the dynamics.
- combining the minChi criterion and the crispness, to avoid solving the full problem for many *n_s_*, but still select the *n_s_* with the crispest decomposition. This is done by first selecting an interval of potentially good numbers of macrostates ns via the minChi criterion and afterwards using the crispness to select the best ns from the preselected macrostate numbers.

All of the above are available through CellRank.

#### Scalable Python implementation of GPCCA

Following the original MATLAB implementation^23^, we wrote up GPCCA as a general algorithm in Python and included it in the `MSMTools`^30^ package, which is widely used for studying protein folding kinetics. From CellRank, we interface to `MSMTools` for the GPCCA algorithm. A naive implementation of the Schur decomposition would scale cubically in cell number. We alleviate this problem by using `SLEPSc` to compute a partial real Schur decomposition using an iterative, Krylov-subspace based algorithm that optimally exploits the sparsity structure of the transition matrix^31,32^. Overall, this reduces the computational complexity of our algorithm to be almost linear in cell number (Fig. 5d and Supplementary Table 1). This allows CellRank to scale well to very large cell numbers.

#### Automatically determine terminal states

We use the coarse-grained dynamics given by *P_c_* to automatically identify terminal states. The idea is to look for the most stable macrostates according to the coarse-grained transition matrix *P_c_*. Define the *stability index* (SI) of a macrostate *m* ∈ {1,…,*n_s_*} through its corresponding diagonal value in *P_c_*, i.e. through its self-transition probability *P_c_* (*m,m*). The intuition behind this is that cells in terminal populations should have very little probability to transition to cells in other populations and should distribute most (if not all) of their probability mass to cells from the same terminal population. To identify the number of terminal states, we set a threshold on SI, i.e. we classify all states as terminal for which SI ≥ *ϵ*_SI_ with ∈ *ϵ*_SI_ = 0.96 by default.

#### Automatically determine initial states

To identify the initial states automatically, we introduce the *coarse-grained stationary distribution* (CGSD) *π_p_*, given by

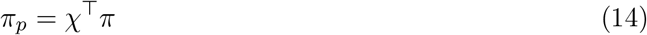

where *π* is the stationary distribution of the original transition matrix *P*. The stationary distribution satisfies

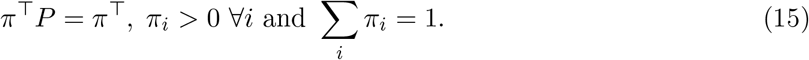

In other words, the stationary distribution *π* is an invariant measure of *P* and can be computed by normalizing the top left eigenvector of *P* (corresponding to the eigenvalue 1). Under certain conditions (ergodicity, see ref. ^33^) imposed on the Markov chain, the stationary distribution is the distribution that the process converges to, if it evolves long enough, i.e. it describes the long-term evolution of the Markov chain. In the same vein, the CGSD *π_c_* describes the long-term evolution of the Markov chain given by the coarse-grained transition matrix *P_c_*. The CGSD *π_c_* assigns large (small) values to macrostates that the process spends a large (little) amount of time in, if it is run infinitely long. As such, we may use it to identify initial states by looking for macrostates which are assigned the smallest values in *π_c_*. The intuition behind this is that initial states should be states that the process is unlikely to visit again once it has left them. The number of initial states is a method parameter set to one by default which can be modified by the user to detect several initial states.

#### Automatically determine intermediate states

All remaining macrostates, i.e. macrostates which have neither been classified as terminal nor as initial, are classified as intermediate. Biologically, these correspond to intermediate, transient cell populations on the state change trajectory.

##### 1.4 Computing fate probabilities

Given the soft assignment of cells to macrostates by *χ* and the identification of terminal states through *P_c_*, we compute how likely each cell is to transition towards these terminal states. Let *n_t_* be the number of terminal states. For the sake of clarity, we assume that we are only interested in fate probabilities towards terminal states, however, the below computations apply just as well to intermediate states, depending on the biological question. For each terminal macrostate *t* for *t* ∈ {1,…,*n_t_*}, we choose f cells which are strongly assigned to t according to *χ*. That is, for terminal macrostate *t*, we extract the corresponding column from *χ* and we calculate the terminal index set 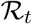 of cells which have the largest values in this column of *χ*. If cell *i* is part of terminal index set 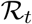, we assume cell *i* is among the *f* most eligible cells to characterize the terminal macrostate *t* in terms of gene expression. We store the indices of the remaining cells in the transient index set 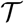. The index sets 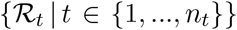 and 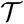 form a disjoint partition of the state space, which means they do not overlap and they cover the entire state space. For each cell *i* in 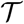, we would like to compute a vector of probabilities *f_i_* ∈ ℝ^*n_t_*^ which specifies how likely this cell is to transition into any of the terminal sets 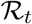. To interpret *f_i_* as a categorical distribution over cell fate, we require 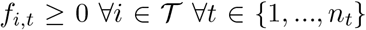 and 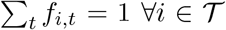. We accumulate the *f_i_* column-wise in the fate matrix *F ∈ R^N×n_t_^*.

#### Absorption probabilities reveal cell fates

We could approximate the *f_i_* based on sampling: initialise a random walk on the Markov chain in cell *i*. Continue to simulate the random walk until any cell from a terminal set 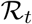 is reached. Record *t* and repeat this many times. Finally, count how often random walks initialized in cell i terminated in any of the terminal index sets 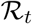. In the limit of repeating this infinitely many times, the normalized frequencies over reaching either terminal set will be equal to the desired fate probabilities for cell *i*, under reasonable assumptions on the Markov chain (irreducibility). Luckily, we do not have to do this in a sampling based approach, we can exploit the fact that a closed form solution exists for this problem: absorption probabilities.

#### Computing absorption probabilities

Key to the concept of absorption probabilities are *recurrent* and *transient* classes, which we will define here for the present case of a finite and discrete state space. Let *i* ∈ Ω and *j* ∈ Ω be two states of the Markov chain. In our case, *i* and *j* are cells. We say that *i* is *accessible* from *j*, if and only if there exists a path from *j* to *i* according to the transition matrix *P*. A *path* is a sequence of transitions which has non-zero transition probability. Further, *i* and *j communicate,* if and only if *i* is accessible from *j* and *j* is also accessible from *i*. Communication defines an equivalence relation on the state space *Ω*, i.e. it is a reflexive, symmetric and transitive relation between two states^33^. It follows that the state space *Ω* can be partitioned into its *communication classes* 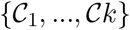. The communication classes are mutually disjoint non-empty and their union is *Ω*. In other words: any two states from the same communication class communicate, states from different communication classes never communicate. We call a communication class 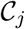 *closed* if the submatrix of *P* restricted to 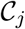 has all rows sum to one. Intuitively, if 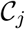 is closed, then a random walk which enters 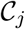 will never leave it again. Closed communication classes are also called *recurrent classes*. If a communication class is not recurrent, we call it *transient*. In Theorem 1, we reproduce the statement of Thm. 28 in ref.^33^ to compute absorption probabilities towards states that belong to recurrent classes on the Markov chain.

#### Theorem 1 (Absorption Probabilities)

*Consider a MC with transition matrix P* ∈ ℝ^*N×N*^. *We may rewrite P as follows*:

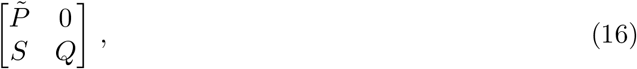

*where 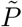 and Q are restrictions of* P *to recurrent and transient states, respectively, and S is the restriction of P to transitions from transient to recurrent states. The upper right 0 is due to the fact that there are no transitions back from recurrent to transient states. Define the matrix M ∈ ℝ^N×N^ via*

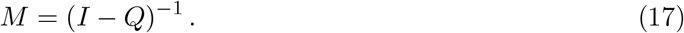

*Then, the ij-th entry of M describes the expected number of visits of the process to state j before absorption, conditional on the process being initialised in state i. M is often referred to as the fundamental matrix of the MC. Further, the matrix*

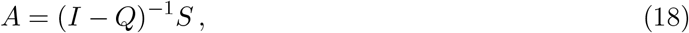

*in the ij-th entry contains the probability of j being the first recurrent state reached by the MC, given that it was started in i.*

For a proof, see See Thm. 26 in ref.^33^. To compute fate probabilities towards the terminal index sets 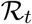 defined above, we approximate these as recurrent classes, i.e. we remove any outgoing edges from these sets. We then apply Theorem 1, which, for each cell 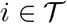 yields absorption probabilities towards each of the *f* cells in each of the *n_t_* recurrent index sets. We aggregate these to yield absorption probabilities towards the recurrent index sets themselves by summing up absorption probabilities towards individual cells in these sets.

#### CellRank provides an efficient implementation to compute absorption probabilities

A naive implementation of absorption probabilities scales cubically in the number of transient cells due to the matrix inversion in Equation (18). The number of transient cells is smaller than the total cell number only by a small constant, so the naive approach can be considered cubic in cell number. This will inevitably fail for large cell numbers. We alleviate this by re-writing Equation (18) as a linear problem,

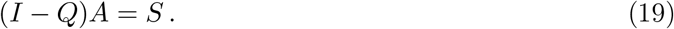

Note that *Q* is very sparse as it describes transitions between nearest neighbors. Per row, *Q* has approximately *K* entries. To exploit the sparsity, iterative solvers are very appealing as their per-iteration cost applied to this problem is linear in cell number and in the number of nearest neighbors. To apply an iterative solver, we must however re-write Equation (19) such that the right hand side is vector valued,

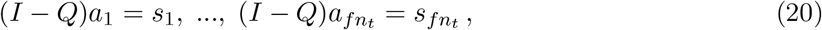

where *fn_t_* is the total number of cells which belong to approximately recurrent classes. To solve these individual problems, we use the iterative GMRES^34^ algorithm which efficiently exploits sparsity. For optimal performance, we use the PETSc implementation, which makes use of efficient message passing and other practical performance enhancements. Lastly, we parallelize solving the *fn_t_* linear problems. All of these tricks taken together allow us to compute absorption probabilities quickly even for large cell numbers (Fig. 5d and Supplementary Table 1).

##### 1.5 Propagating velocity uncertainty

So far, we have assumed that individual velocity vectors are deterministic, i.e. they have no measurement error. However, this is not correct because RNA velocity is estimated on the basis of spliced and unspliced gene counts, which are noisy quantities. Hence, the velocity vectors *v_i_* themselves should be treated as random variables which follows a certain distribution. Ultimately, our aim is to propagate the distribution in *v_i_* into our final quantities of interest, i.e. state assignments and fate probabilities. However, this is difficult as these final quantities of interest depend on *v_i_* in non-analytical ways, i.e. we cannot write down a closed-form equation which relates the final quantities to *v_i_*. A possible solution to this is to use a Monte Carlo scheme where we draw velocity vectors, compute final quantities based on the draw and repeat this many times. In the limit of infinitely many draws, this will give us the distribution over final quantities, given the distribution in *v_i_*. However, this has the disadvantage that we need to repeat our computation many times, which will get prohibitively expensive for large datasets. To get around this problem, and to allow CellRank to scale to large datasets, we construct an analytical approximation to the Monte Carlo based scheme. This analytical approximation will only have to be evaluated once and we can omit the sampling. We show in a practical example that the analytical approximation gives very similar results to the sampling based scheme and improves over a deterministic approach by a large margin.

#### Modeling the distribution over velocity vectors

Before we can propagate uncertainty, we need to describe the distribution over velocity vectors, i.e. we need to model the uncertainty present in the velocity vectors which are estimated by scVelo^9^ or velocyto^1^. Ideally, we would like these packages themselves to model uncertainty in the raw spliced and unspliced counts and to propagate this into a distribution over velocity vectors. However, as that is currently not being done, we will make an assumption about their distribution and use the KNN graph to fine-tune expectation and variance by considering neighboring velocity vectors. To ease notation and to illustrate the core ideas, we will drop the subscript *i* in this section and just focus on one fixed cell and it’s velocity vector *v*. Let’s assume that *v* follows a multivariate normal (MVN) distribution,

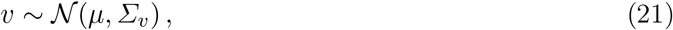

with mean vector *μ* ∈ ℝ^*G*^ and covariance matrix *Σ_v_* ∈ ℝ^*G×G*^. The MVN is a reasonable choice here as velocities can be both positive and negative and for most genes, as we expect to see both up-and down-regulation, velocity values will be approximately symmetric around their expected value. Let’s further assume the covariance matrix to be diagonal, i.e. gene-wise velocities are independent. This is a reasonable assumption to make as gene-wise velocities in both velocyto^1^ and scvelo^9^ are computed independently. To compute values for *μ* and *Σ_v_*, consider velocity vector *v* and its *K* nearest neighbors. To estimate *μ* and the diagonal elements of *Σ_v_*, we compute first and second order moments over the velocity vectors of these neighboring cells.

#### Propagating uncertainty into state assignments and fate probabilities

We seek to approximate the expected value of the final quantities of interest (state assignments and fate probabilities), given the distribution in the velocity vectors. Let *q* be a final quantity of interest. There are two major steps involved in computing *q*,

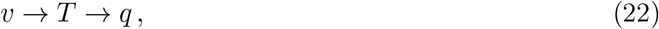

where *v* stands for our inputs, i.e. the velocity vectors, and *T* is the transition matrix defining the Markov chain. To get from *v* to *T*, we evaluate an analytical function which computes correlations and applies a softmax function. We can approximate this first part of the mapping with a Taylor series, which allows us to analytically propagate the distribution in *v* into *T*. For the second part of the mapping, we use the expected transition matrix to compute *q*. This yields an approximation to the expectation of the final quantity we can then compare with the approximation we obtain from a Monte Carlo scheme, which we treat as our ground truth.

#### Approximating the expected transition matrix

In the first step, we compute the expected value of the transition matrix, given the distribution of the velocity vectors. Given a particular draw *v* from the distribution in Equation (21) and a set of state-change vectors *s_k_*, we compute a vector of probabilities *p*, which lives on a *K*-simplex in ℝ^*K*^. Let’s denote the mapping from *v* to *p* by *h*,

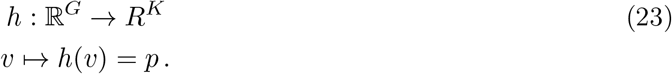

We can then formulate our problem as finding the expectation of *h* when applied to *v*, i.e.

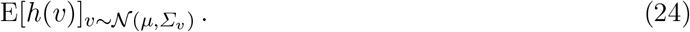

To approximate this, expand the i-th component of *h* in a Taylor-series around *μ*,

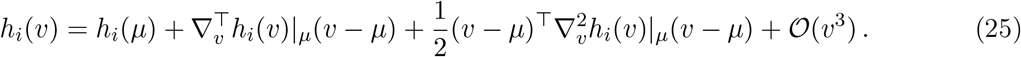

Define the Hessian matrix of *h_i_* at *v = μ* as

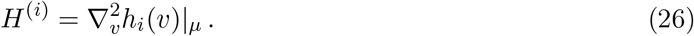

Taking the expectation of *h_i_* and using the Taylor-expansion,

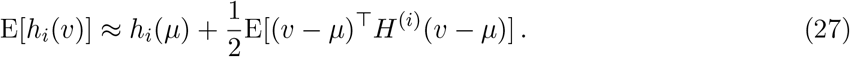

Note that the first order term cancels as E[*v – μ*] =0. The second order term can be further simplified by explicitly writing out the matrix multiplication,

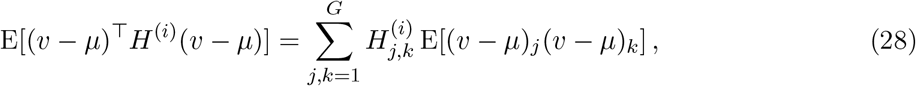

where we took the expectation inside the sum and the matrix elements outside the expectation as it does not involve *v*. For *j* ≠ *i*, the two terms inside the expectation involving *v* are independent given our distributional assumptions on *v* and the expectation can be taken separately. Using again the fact that E[*v* – *μ*] = 0, the sum equals zero for *j* ≠ *i*. It follows

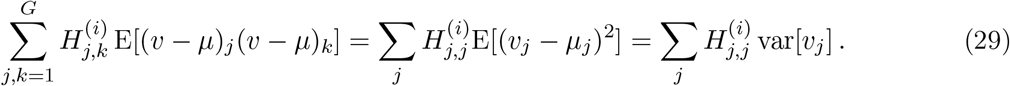

To summarize, our second order approximation to the transition probabilities given the distribution in *v* reads

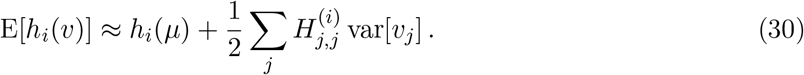

We use automatic differentiation as implemented in JAX^35^ to compute the Hessian matrices *H*^(*i*)^, which ensures they are highly accurate and can be computes in a scalable manner. Further, because we do not hard-code the derivatives, our approach is highly flexible to future changes in the way we compute transition probabilities. If for example it turns out at a later point that an alternative metric works better than Pearson correlation, this is automatically taken care of in the propagation of uncertainties and no changes need to be made, apart from changing the forwards function which computes the transition probabilities themselves. The above procedure can be repeated for all components *i* and for all cells to yield the second order approximation to the expected transition matrix *T*, given the distribution over each velocity vector.

#### Approximating the expected final quantities

To arrive at the final quantities of interest, i.e. state assignments and absorption probabilities, we use the expected transition matrix and proceed as in the deterministic case. We validate that this approximation gives very similar results to a fully stochastic approach based on Monte Carlo sampling (Supplementary Fig. 14a,b).

##### 1.6 The CellRank software package

The CellRank software package implements two main modules:

- `kernels` are classes that provide functionality to compute transition matrices based on (directed) single cell data.
- `estimators` are classes that implement algorithms to do inference based on `kernels`. For example, `estimators` compute macrostates and fate probabilities.

This modular and object oriented design allows CellRank to be extended easily into two directions. On the one hand, including more kernels to take into account further means of directional single cell data such as metabolic labeling or experimental time. On the other hand, including more estimators to learn new abstractions of cellular dynamics. The kernel module currently implements a

- `VelocityKernel` which computes a transition matrix on the basis of a KNN graph and RNA velocity information.
- `PalantirKernel` which mimicks the original routine outlined in the Palantir^8^ paper to compute a directed transition matrix on the basis of a KNN graph and a pseudotime.
- `ConnectivityKernel` which takes the adjacency matrix underlying the KNN graph and row-normalizes it to obtain a valid transition matrix. This is essentially the transition matrix used in the DPT^10^ algorithm.
- `PrecomputedKernel` which accepts any pre-computed transition matrix and allows for easy interfacing with the CellRank software.

All `kernel` classes are derived from a `base kernel` class which implements density normalization as implemented in ref.^10^. Instances of `kernel` classes can be combined by simply adding them up using the + operator, potentially including weights. A typical code snippet to compute a transition matrix will look like this:

~~~
from cellrank.tools.kernels import VelocityKernel, ConnectivityKernel
vk = VelocityKernel(adata).compute_transition_matrix()
ck = ConnectivityKernel(adata).compute_transition_matrix()
combined_kernel = 0.9*vk + 0.1*ck
~~~

The estimator module currently implements a

- `CFLARE` estimator. `CFLARE` stands for *Clustering and Filtering of Left and Right Eigenvectors*. This estimator computes terminal states directly by filtering cells in the top left eigenvectors and clustering them in the top right eigenvectors, thereby combining ideas of spectral clustering and stationary distributions.
- `GPCCA` estimator. The GPCCA estimator.

All `estimator` classes are derived from a `base estimator` class which allows to compute fate probabilities, regardless of how terminal/intermediate states have been computed. A typical code snippet to compute macrostates and fate probabilities will look like this:

~~~
from cellrank.tools.estimators import GPCCA
# initialise the estimator
gpcca = GPCCA(combined_kernel)
# compute macrostates and identify the terminal states among them
gpcca.compute_macrostates()
gpcca.compute_terminal_states()
# comptue fate probabilities
gpcca.compute_absorption_probabilities()
~~~

Both `kernels` and `estimators` implement a number of plotting functions to conveniently inspect results. We designed CellRank to be highly scalable to ever increasing cell numbers, widely applicable and extendable to problems in single cell dynamical inference, user friendly with tutorials and comprehensive documentation and robust with 88 % code coverage. CellRank is open source, fully integrated with SCANPY and scVelo and freely available at https://cellrank.org.

### 2 Computing a directed PAGA graph

Partition-based graph abstraction (PAGA)^36^ provides an interpretable graph-like connectivity map of the data manifold. It is obtained by associating a node with each manifold partition (e.g. cell type) and connecting each node by weighted edges that represent a statistical measure of connectivity between the partitions. The model considers groups/nodes as connected if their number of inter-edges exceeds what would have been expected under random assignment. The connection strength can be interpreted as confidence in the presence of an actual connection and allows discarding spurious, noise-related connections.

Here, we extend PAGA by directing the edges as to reflect the RNA velocity vector field rather than transcriptome similarity. The connectivity strengths are defined based on the velocity graph. That is, for each cell correlations between the cell’s velocity vector and its potential, cell-to-cell transitions are computed (Subsection 1.2). Inter-edges are considered whose correlation passes a certain threshold (default: 0.1). The number of inter-edges are then tested against random assignment for significance.

To further constrain the single cell graph, compute a gene-shared latent time using scVelo^37^. In short, this aggregates the per-gene time assignments computed in scVelo’s dynamical model to a global scale which faithfully approximates a single-cells internal clock. Once we have computed the initial states using CellRank, we can use these as a prior for latent time to force it to start in this state. All of latent time, initial and terminal states can in turn be used as a prior to regularize the directed graph. At single-cell level, we use latent time as a constraint to prune the cell-to cell transition edges to those that match the ordering of cells given by latent time. For the initial and terminal states, the edges are further constrained to only retain those cell-to-cell transitions that constitute outgoing flows for cells in initial cellular populations, and to incoming flows for cells in terminal populations.

Finally, a minimum spanning is constructed for the directed abstracted graph. It is obtained by pruning node-to-node edges such that only the most confident path from one node to another is retained. If there are multiple paths to reach a particular node, only the path with the highest confidence is kept.

### 3 Computing gene expression trends along lineages

CellRank computes fate probabilities which specify how likely each individual cell is to transition towards each identified terminal state (Subsection 1.4). Combined with any pseudo-temporal measure like DPT^10^, scVelo’s latent time^2^ or Palantir’s pseudotime^38^, this allows us to compute and to compare gene expression trends towards specific terminal populations. In contrast to other methods, we do not partition the set of all cells into clusters and define lineage each lineage via an ordered set of clusters. Instead, we use all cells to fit each lineage but we weigh each cell according to its fate probability, our measure of lineage membership. This means that for cells uncommitted between two or more fates, we allow them to contribute to each one of these, weighted by the fate probabilities. For cells committed towards any particular fate, their fate probabilities towards the remaining fates will be zero or almost zero which naturally excludes them when fitting these other lineages.

#### Imputing gene expression recovers trends from noisy data

To improve the robustness and resolution of gene expression trends, we adapt two strategies. First, we use imputed gene expression values and second, we fit Generalized Additive Models (GAMs). For gene expression imputation, we use MAGIC^39^ by default, however, any imputed gene expression matrix can be supplied. MAGIC is based on KNN imputation and makes use of the covariance structure among neighboring cells to estimate expression levels for each gene. The KNN graph is computed globally, based on the expression values of all genes and not just the one we are currently considering.

#### Generalized Additive Models (GAMs) robustly fit gene expression values

While sliding window approaches are known to be sensitive to local density differences and only take into account the current gene when determining gene expression trends, we fit GAMs to gene expression values which have been imputed borrowing information from neighboring cells via a KNN graph. Using GAMs allows us to flexibly model many different kinds of gene trends in a robust and scalable manner. We fit the gene expression trend for branch *t* in gene *g* via

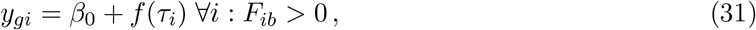

where *y_gi_* is gene expression of gene *g* in cell *i, τ_i_* is the pseudotemporal value of cell *i* and *F* is the fate matrix of Subsection 1.4. By default, we use cubic splices for the smoothing functions *f* as these have been shown to be effective in capturing non-linear relationships in trends^40^.

To visualize the smooth trend, we select 200 equally spaced testing points along pseudotime and we predict gene expression at each of them using the fitted model of Equation (31). To estimate uncertainty along the trend, we use the standard deviation of the residuals of the fit, given by

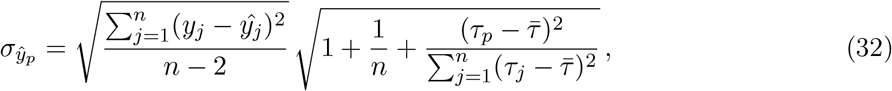

where *ŷ_p_* denotes predicted gene expression at test point 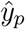 denotes average pseudo-time across all cells and n is the number of test points^41^. For the fitting of Equation (31), we provide interfaces to both the R package `mgcv`^42^ as well as the Python package `pyGAM`^43^. We parallelize gene fitting to scale well in the number of genes, which is important when plotting heatmaps summarizing many gene expression trends.

#### Visualizing gene expression trends for the pancreas example

For CellRank’s gene expression trends of lineage-associated genes along the alpha, beta, epsilon and delta fates, we used Palantir’s pseudotime^38^, MAGIC imputed data^39^ and the `pyGAM`^43^ package to fit GAMs. We used default values to fit the splines, i.e. we place 10 knots along pseudotime and we use cubic splines. For the delta lineage, fate probabilities among early cells were very low (0.01 average fate probability among Ngn3 high EP cells, see Fig. 2e). This reflects the small size of the delta population (70 cells or 3% of the data, see Supplementary Fig. 10a,b) as well as the fact that delta cells are produced mostly at later stages in pancreatic development^44^. To still be able to reliably fit gene expression of early cells along the delta lineage, we thresholded weights at 0.05, i.e. weights smaller than this value were clipped to this value. This was done only for the fitting of gene expression trends.

### 4 Clustering gene expression trends

CellRank allows gene expression trends along a particular lineage to be clustered, thus recovering the major patterns of regulation towards a specific terminal state like (transient) up-or down-regulation. For the set of genes we are interested in, we recover their regulation along a specific lineage by fitting GAMs in pseudotime where we supply fate probabiliteis as cell-level lineage weights (Section 3). In the next step, we cluster the GAM-smoothed gene expression trends. For this, we z-transform expression values and we compute a PCA representation of the trends. By default, we use 50 PCs. We then compute a KNN graph in PC space as outlined in Subsection 1.2 and we cluster the KNN graph using the louvain^45^ or leiden^46^ algorithms. For each recovered cluster, we compute its mean and standard deviation (point-wise, for all testing points that were used for smoothing) and visualize them, together with the individual, smoothed trends per cluster. As gene-trend fitting is efficiently parallelized in CellRank, such an analysis can be performed in an unbiased fashion for large gene sets. For 10k genes, the run-time is about 6 min on a 2019 Mac book pro with 2,8 GHz Intel Core i7 processor and 16 GB RAM.

#### Clustering gene expression trends towards the delta fate

To cluster gene expression trends towards the delta fate in Fig. 3e, all genes which were expressed in at least 10 cells were included (12,987 genes). Smooth gene expression trends along the delta lineage were determined using Palantir’s pseudotime^38^. We used *K* = 30 nearest neighbors for the gene-trend KNN graph and the louvain algorithm with resolution parameter set to 0.2 to avoid over-clustering the trends.

### 5 Uncovering putative driver genes

To find genes which are expressed at high levels in cells that are biased towards a particular fate, we compute Person’s correlation between expression levels of a set of genes and fate probabilities. We sort genes according to their correlation values and consider high-scoring genes as candidate driver genes. By default, we consider all genes which have passed pre-processing gene filtering thresholds. The computation of correlation values can be restricted to a set of pre-defined clusters if one is interested in driver genes which act in a certain region of the phenotypic manifold.

#### Uncovering putative driver genes for delta development

To uncover putative driver genes towards the delta fate in Fig. 3d,e, we considered 12,987 genes which were expressed in at least 10 cells. We computed correlation of total-count normalized, log transformed gene expression values with the probability of becoming a delta cell. We restricted correlation computation to the Fev+ cluster, where we expected the fate decision towards delta to occur.

### 6 Robustness analysis

We were interested in evaluating how much CellRank’s fate probabilities change in response to changes in the following key pre-processing parameters:

- the number of neighbors *K* used for KNN graph construction (Subsection 1.2)
- scVelo’s gene-filtering parameter `min_shared_counts` which determines how many counts a gene must have in both spliced and unspliced layers
- scVelo’s gene filtering parameter `n_top_genes` which determines the number of most highly variable genes used for the velocity computation
- the number of principal components `n_pcs` used for KNN graph construction (Subsection 1.2)

In addition to the 4 key pre-processing parameters, we were interested to see how much CellRank’s results change when we randomly sub-sample the number of cells to 90% of the original cell number and when we vary the number of macrostates. We used the pancreas example^17^ in all of the following comparisons.

#### Robustness with respect to key pre-processing parameters

To evaluate robustness with respect to changes in the pre-processing parameters, we varied one parameter at a time (keeping the others fixed), computed macrostates and fate probabilities towards them. We then compared the fate probabilities for different values of the parameter by computing pairwise Pearson correlation among all possible pairs of values we used for the parameter. We did this separately for each lineage, i.e. for the alpha, beta, epsilon and delta lineages. For each lineage, we recorded the median and minimum correlation achieved across all the different comparisons. We always computed enough macrostates so that the alpha, beta, epsilon and delta states were included. Naturally, the precise location of the terminal states changed slightly across parameter combinations. For this reason, the correlation values we recorded reflect robustness of the entire CellRank workflow, including both the computation of terminal states as well as fate probabilities. In a separate comparison, we were interested in evaluating the robustness of just the last step of the CellRank algorithm, i.e. the computation of fate probabilities. For this, we kept the terminal states fixed across parameter variations and proceeded as above otherwise, computing pairwise Pearson correlations among fate probabilities per lineage across all parameter value combinations. Furthermore, we were interested to see whether CellRank’s robustness changes when we propagate uncertainty. For this, we repeated all the aforementioned computations using our analytical approximation to propagate uncertainty.

#### Robustness with respect to random sub-sampling of cells

We subsampled the data to 90% of cells, computed macrostates and fate probabilities towards the alpha, beta, epsilon and delta states. We repeated this 20 times, recorded all computed fate probabilities and compared them pairwise per lineage using Pearson’s correlation for all possible pairs of random draws. As in the above evaluation for the key pre-processing parameters, we recorded minimum and median correlation per lineage across all pairs and we repeated this for fixed terminal states and for propagated uncertainty.

#### Robustness with respect to the number of macrostates

To evaluate sensitivity with respect to this parameter, we varied the number of macrostates between 10 and 16 and confirmed that within this range, the key terminal and initial states exist and remain in the same location.

### 7 Pancreas data example

We used a scRNA-seq time-series data set comprising embryonic days 12.5 — 15.5 of pancreatic development in mice assayed using 10x Genomics^17^. We restricted the data to the last time point (E15.5) and to the Ngn3 low EP, Ngn3 high EP, Fev+ and endocrine clusters to focus on the late stages of endocrinogenesis where all of alpha, beta, epsilon and delta fates are present. We further filtered out cycling cells to amplify the differentiation signal. Our final subset contained 2531 cells. We kept the original cluster annotations which were available on a coarse level and on a fine level. On the fine level, the Fev+ cluster was sub-clustered into different populations which are biased towards different endocrine fates (Supplementary Fig. 10c).

#### Data pre-processing and velocity computation

For the following processing, we used scVelo^9^ and SCANPY^14^ with mostly default parameters. Loom files containing raw spliced and unspliced counts were obtained by running the velocyto^1^ command-line pipeline. We filtered genes to be expressed in at least 10 cells and to have at least 20 counts in both spliced and unspliced layers. We further normalized by total counts per cell, log transformed the data and kept the top 2000 highly variable genes. We then computed a PCA representation of the data and used the top 30 PCs to compute a KNN graph with *K* = 30 nearest neighbors. For velocity computation, we used scVelo’s dynamical model of splicing kinetics. We evaluate robustness of CellRank’s results to changes in these pre-processing parameters (Section 6).

#### Embedding computation

We used the KNN graph to compute a PAGA^7^ representation of the data. The PAGA graph was used to initialize the computation of a UMAP^12,47^ representation of the data. Note that UMAP was only used to visualize the data and was not supplied to CellRank to compute the transition matrix or any downstream quantities.

#### CellRank parameters

We use CellRank’s analytical stochastic approximation to compute transition probabilities and include a diffusion kernel with weight 0.2 (Subsection 1.2 and Subsection 1.5). We compute 12 macrostates and automatically detect the terminal alpha, beta and epsilon states. The delta population is picked up automatically as a macrostate. We manually assign it the terminal label.

#### Statistical testing

To check whether Fev+ delta cells were assigned significantly higher delta fate probability compared to other Fev+ clusters by CellRank, we applied a two-sided Welch unequal variances t-test. The test assumes two independent normally distributed samples with unequal variances and checks whether their means are significantly different.

### 8 Lung data example

We used a scRNA-seq time-series data-set of lung regeneration past bleomycin injury in mice assayed using Dropseq^16,48^. The data-set contained 18 time points comprising days 0-54 past injury. There was daily sampling from days 2-13 and wider time-lags between the following time-points. Two replicate mice were used per time point. We restricted the data to days 2-15 to make sure that the sampling is dense enough for velocities to be able to meaningfully extrapolate gene expression. If time points are too far apart, then RNA velocity cannot be used to predict the next likely cellular state because the linear extrapolation is only meaningful on the time scales of the splicing kinetics. Our final subset contained 24,882 cells. We kept the original cluster annotations.

#### Data pre-processing and velocity computation

For the following processing, we used scVelo^9^ and SCANPY^14^ with mostly default parameters. Loom files containing raw spliced and unspliced counts were obtained by running the velocyto^1^ command-line pipeline. We filtered genes to be expressed in at least 10 cells and to have at least 20 counts in both spliced and unspliced layers. We further normalized by total counts per cell, log transformed the data and kept the top 2000 highly variable genes. We kept the PCA coordinates from the original study and computed a KNN graph with K = 30 nearest neighbors using the top 50 PCs. For velocity computation, we used scVelo’s dynamical model of splicing kinetics.

#### Embedding computation

The lung data was processed in three separate batches. We used BBKNN^49^ to compute a batch corrected KNN graph with 10 neighbors within each batch. The corrected KNN graph was used to compute a UMAP^12,47^ representation of the data. Note that UMAP was only used to visualize the data and was not supplied to CellRank to compute the transition matrix or any downstream quantities. We did not use BBKNN to correct the graph we used for velocity computation as it is an open question how to do batch correction for velocity computation. We used uncorrected data for velocity computation.

#### CellRank parameters

We use CellRank’s analytical stochastic approximation to compute transition probabilities and include a diffusion kernel with weight 0.2 (Subsection 1.2 and Subsection 1.5). On the full data of Fig. 6a, we compute 9 macrostates. On the reduced data of Fig. 6d, we compute 3 macrostates.

#### Statistical testing

Analogous to the corresponding paragraph in the pancreas example above (Section 7), we used the two-sided Welch unequal variances t-test to check whether goblet cells were assigned significantly higher basal probability than other secretory airway cells.

#### Defining stages of the differentiation trajectory

We sub-setted cells to goblet and basal cells and re-run CellRank on the subset to investigate the trajectory at higher resolution. CellRank automatically detected initial and terminal states and computed fate probabilities towards the terminal states (Supplementary Fig. 24a-c). Further, we applied Palantir^38^ to the subset to compute a pseudotime (Supplementary Fig. 24d,e). We combined the pseudotime with CellRank’s fate probabilities to define three stages of the dedifferentiation trajectory by requiring cells to have at least 0.66 basal probability. Cells passing this threshold were assigned to three bins of equal size along the pseudotemporal axis. We used this binning to define the three stages of the trajectory.

### 9 Methods comparison

We compared CellRank with the following similarity-based tools that compute probabilistic fate assignments on the single cell level: Palantir^38^, FateID^50^ and STEMNET^51^. We compared these methods in terms of the identification of initial and terminal states, fate probabilities, gene expression trends and run-time.

#### The Palantir algorithm

Palantir^38^ computes a KNN graph in the space of diffusion components and uses this graph to compute a pseudotime via iteratively updating shortest path distances from a set of waypoints. Palantir required us to provide a number of *waypoint cells* – essentially a smaller number of cells that the system is reduced to in order to make it computationally feasible. We set this number to 1200 cells for the pancreas data and to 15% of the total cell number of the runtime and memory benchmarks on the reprogramming dataset^52^ below. For pseudotime computation, an initial cells needs to be supplied by the user. The pseudotime is used to direct edges in the KNN graph by removing edges that point from later cells to earlier cells in pseudotime. The stationary distribution of the resulting directed transition matrix is combined with extrema in the diffusion components to identify terminal cells. Absorption probabilities towards the terminal cells serve as fate probabilities. Gene expression trends are computed similarly to CellRank, by fitting GAMs in pseudotime where each cell contributes to each lineage according to its fate probabilities.

#### The FateID algorithm

FateID ^50^ either requires the user to provide terminal populations directly or through a set of marker genes. Terminal populations are used to train a random forest classifier. The classifier is applied to a set of cells in the neighborhood of each terminal cluster where it predicts the likely fate of these cells. The training set is iteratively expanded and the Random forest is re-trained on expanding populations, thus moving from the committed populations backward in time, classifying the fate of increasingly earlier cells. Two key parameters here are the size of the training and test sets used for the Random forest classifier, which we set to 1% of the data in all benchmarks. Gene expression trends are computed by selecting a (discrete) set of cells which pass a certain threshold for fate bias towards a specific terminal population. A principal curve is fit to these cells in a low dimensional embedding and pseudotime is assigned via projection onto this curve. Alternatively, the authors recommend to compute diffusion pseudotime^10^ (DPT) on the set of cells selected for a particular lineage. Gene expression values are then normalised and a local regression (LOESS) is performed to obtain mean trends. In contrast to CellRank and Palantir, this approach does not provide confidence intervals for the expression trends, it is dependent on low dimensional embeddings (principal curve fit) and it discreetly assigns cells to lineages, thereby ignoring the gradual nature of fate commitment when visualizing gene expression trends. Since different cells are selected for different lineages, the computed pseudo-temporal orderings are incompatible and gene trends along different lineages cannot be visualized jointly.

#### The STEMNET algorithm

STEMNET^51^ requires the user to provide the terminal populations directly as input to the algorithm. It then trains an elastic-net regularized generalized linear model on the terminal populations to predict state membership. This first step serves as feature selection – it selects a set of genes which are specific to their terminal populations. In the next step, the classifier uses expression of these genes to predict fate bias for the remaining, transient cells. STEMNET uses the computed fate probabilities to place cells on a simplex in 2 dimensions as a dimensionality reduction method. It does not offer a method to visualize gene expression trends.

#### Fate probabilities

In order to enable a fair comparison across methods, we supplied all methods with CellRank’s identified terminal states and compared predicted fate probabilities. Methods differed in the format they require terminal state information to be passed: for Palantir, we passed individual cells, i.e. 4 cells, one for each of the three terminal states and one initial cell from the initial state. For STEMNET, we passed populations of cells defined through the underlying transcriptomic clusters. We passed the alpha, beta, epsilon and delta clusters defined through the sub-clustering of Supplementary Fig. 10c. For FateID, we passed marker genes to identify the terminal populations. For each terminal state, we passed its corresponding hormone-production associated gene, i.e. *Ins1* for beta, *Gcg* for alpha, *Ghrl* for epsilon and *Sst* for delta^53^. We checked whether methods correctly predicted beta to be the dominant fate among early cells by computing the average fate prediction among Ngn3 high EP cells.

#### Gene expression trends

To visualize gene expression trends, we used the functionalities that each method provided. STEMNET did not have an option to compute gene expression trends. For CellRank, we visualized gene expression trends as described in Section 3. For Palantir, we used default parameters. For FateID, it was difficult to find a good threshold value to assign cells to lineages. If this value is too high, then early cells in the trajectory are not selected and the terminal states are isolated. If this value is too low, then for a subset of the lineages, very unlikely cells are assigned and the trends are very unspecific. The default value is 0.25, which was too high in our case. We decided to set the threshold at 0.15, which was a compromise between trying to have early cells in every lineage and making sure that irrelevant cells are not assigned. We computed DPT^10^ on the set of the selected cells, as recommended in the original publication. To identify a root cell for each lineage, we first computed DPT on the entire data-set, then subsetted to a lineage-specific set of cells and picked the cell with the earliest original DPT value as the root cell for the second DPT computation. We visualized expression trends for the key lineage drivers *Pax4*^54^ and *Pdx1*^55–57^ (beta), *Arx*^54^ (alpha), *Ghrl*^53^ (epsilon) and *Hhex*^58^ and *Cd24a*^59,60^ (delta) as well as the lineage accociated genes *Peg10*^61,62^ (alpha) and *Irs4*^62^ (epsilon). In Fig. 5c, we checked whether methods correctly predicted upregulation of *Pdx1* along the beta fate.

#### Runtime

We compared run-time of the four methods applied to a scRNA-seq dataset comprising 100k cells undergoing reprogramming from mouse embryonic fibroblasts to induced endoderm progenitor cells^52^. We randomly subsampled the data-set to obtain 10 data-sets of increasing size, starting from 10k cells in steps of 10k until 100k cells. For each sub-sampled dataset, we applied each method 10 times and computed the mean runtime as well as the standard error on the mean.

We used CellRank to compute 3 terminal states and we supplied these to all other methods to ensure that the number of terminal states is consistent across methods. Methods differed in the format they require terminal state information to be passed: for Palantir, we passed individual cells, i.e. three terminal cells and one initial cell (taken from the earliest time point of the reprogramming data). For STEMNET, we passed a set of cells for each terminal state by choosing the cells which have been most confidently assigned to each terminal state by CellRank. For each terminal state, we passed a number of cells that was equal to 1% of the total cell number. FateID requires marker genes to identify the terminal populations, so we computed the top 3 lineage drivers per CellRank-identified terminal state and passed these.

For CellRank, we separately recorded the time it took to compute the terminal states and fate probabilities. For terminal states in CellRank, we included in this benchmark the entire workflow from computing the transition matrix via decomposing it into macrostates to identifying the terminal states among the macrostates. For fate probabilities, we benchmarked the `compute_absorption_probabilities()` method (CellRank), the `run_palantir()` function (Palantir), the `fateBias()` function (FateID) and the `runSTEMNET()` function (FateID).

Comparisons were run on an Intel(R) Xeon(R) Gold 6126 CPU @ 2.60GHz and 32 cores. Each job was allocated at least 90 GiB RAM and we recorded the actual peak memory usage (see below). FateID did not finish on 100k cells because of a memory error due to densification of a large matrix.

#### Peak memory usage

The setup was identical to the setup for the runtime comparison above, only that we recorded peak memory usage of each method (Supplementary Table 2). For the Python-based methods CellRank and Palantir, we used the `memory-profiler`^63^ package whereas for the R-based packages STEMNET and FateID, we used the peakRAM^64^ profiler. CellRank and Palantir efficiently parallelize their computations across several cores which increases their peak memory consumption. We repeated our evaluation for these two methods on 100k cells using just a single core to estimate the size of this effect (Supplementary Table 3).

### 10 Immunofluorescence stainings and microscopy on airway epithelial cells

Formalin-fixed paraffin-embedded lung sections (3.5 *μm* thick) from bleomycin-treated mice at day 10 (n=2) and day 22 (n=2) after bleomycin instillation, and from phosphate-buffered saline (PBS)-treated controls (n=2) were stained as previously described^16^. In brief, after deparaffinization, rehydration and heat-mediated antigen retrieval with citrate buffer (10 mM, pH = 6.0), sections were blocked with 5% bovine serum albumin for 1 h at room temperature and then incubated with the following primary antibodies overnight at 4°C: rabbit anti-Bpifb1 (kindly provided by C. Bingle^65^, 1:500), mouse anti-Trp63 (abcam, ab735, clone A4A, 1:50) and chicken anti-Krt5 (BioLegend, Poly9059, 1:1,000).

For visualization of stainings the following secondary antibodies were used: Goat anti-rabbit Alexa Fluor® 488 (Invitrogen, A11008, 1:250), Goat anti-chicken Alexa Fluor® 568 (Invitrogen, A11041,1:250) and goat anti-mouse Alexa Fluor® 647 (Invitrogen, A21236, 1:250). Cell nuclei were visualized with 4’,6-diamidino-2-phenylindole (DAPI).

Immunofluorescent images were acquired with an AxioImager.M2 microscope (Zeiss) using a PlanApochromat 20x/0.8 M27 objective. For quantification of immunofluorescence stainings, five different intrapulmonary regions were recorded per mouse and the percentage of positively stained cells normalized to the total number of airway cells was manually quantified using Fiji software (ImageJ, v. 2.0.0).

