## Supplementary Figures for "CellRank for directed single-cell fate mapping"

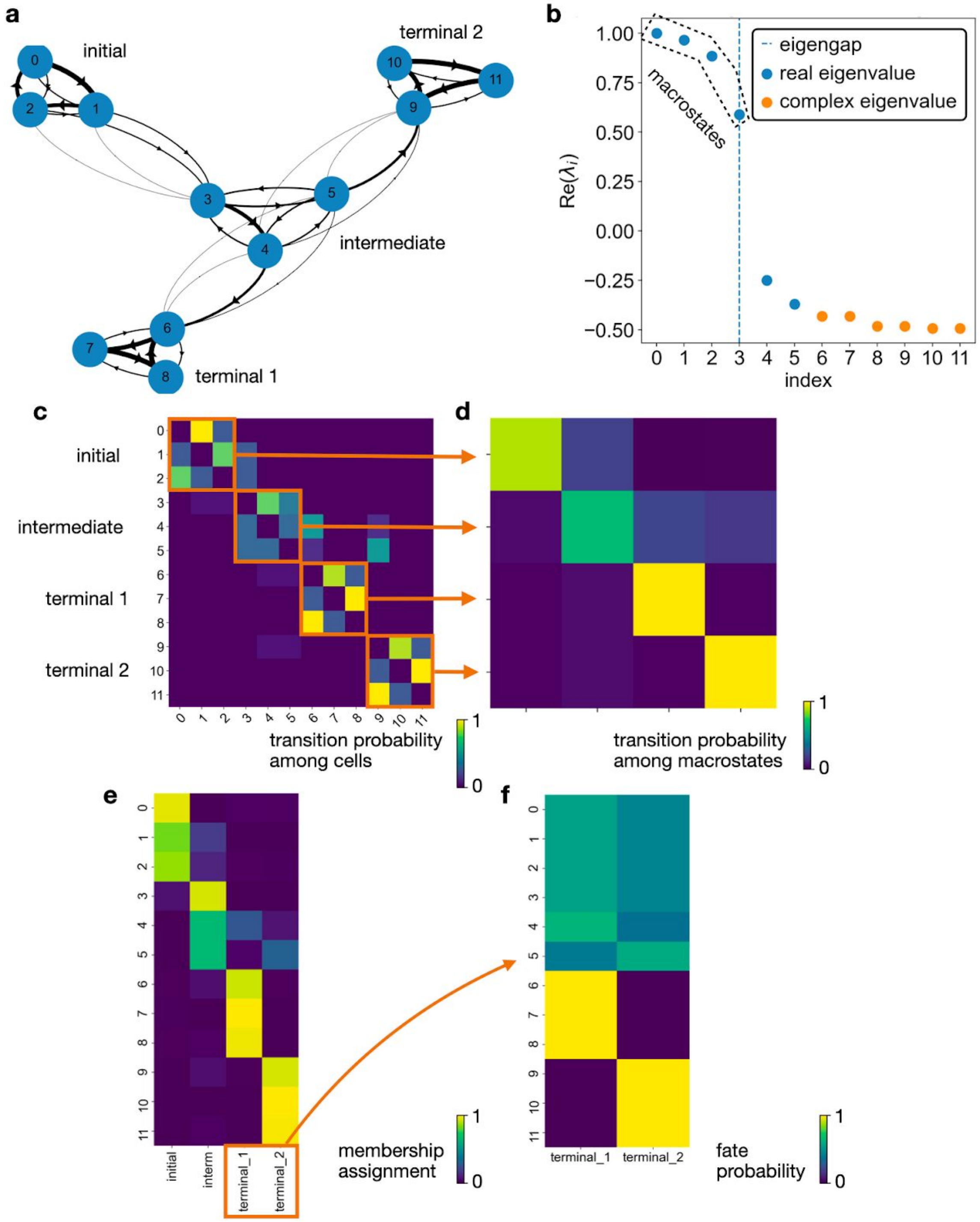


**Supplementary Fig. 1: GPCCA and fate probabilities extract the essence of cellular state transitions**

**a.** Markov transition graph of a toy example of cellular state changes. Starting from a cyclic initial state, cells transition via an intermediate state into either one of two terminal states, both of which are cycling again. Note that cell number 3 is more likely to go to cell number 4 than 5, which results in a global fate bias towards the first terminal state. **b.** The corresponding transition matrix can be decomposed into real Schur vectors, each corresponding to one eigenvalue. The 4 eigenvalues close to one are associated with the initial, terminal and intermediate states. Complex eigenvalues appear because the transition matrix is non-symmetric. **c.** The original transition matrix. The block structure shows the separation into the 4 macrostates and the possible transitions between them. **d.** The coarse-grained transition matrix, identifying the different macrostates and their transition probabilities relative to one-another. The initial state is the macrostate with almost no incoming but large outgoing transition probability. The intermediate state is the state with both large incoming and large outgoing transition probability, and relatively little self-transition probability. The terminal states are the states with large incoming, but almost no outgoing and large self-transition probability. **e.** Each macrostate is associated with a membership vector that assigns cells to the state in a soft fashion, i.e. using weights that sum to one. We show the 4 membership vectors in a heatmap. **f.** Fate probabilities towards the two terminal states. We correctly recover the global bias towards the first terminal state.

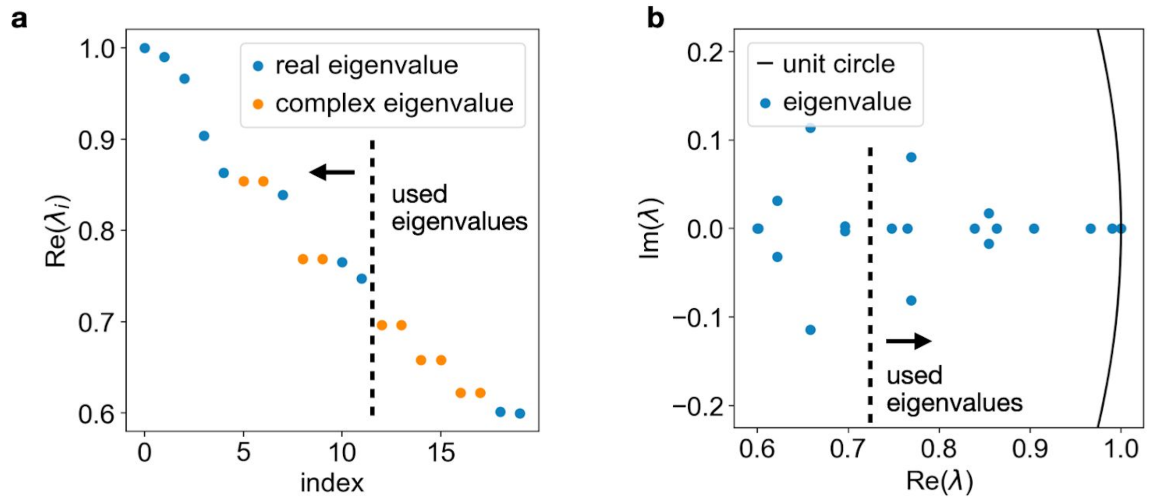


**Supplementary Fig. 2: Spectrum of the pancreas transition matrix**

**a.** Real part of the top 20 eigenvalues. Purely real eigenvalues are shown in blue. Complex eigenvalues come in pairs of complex conjugates for real matrices and are shown in orange. Dashed line highlights the first 12 eigenvalues, which we use to compute macrostates in Fig. 2. **b.** Eigenvalues from (**a**) in the complex plane.

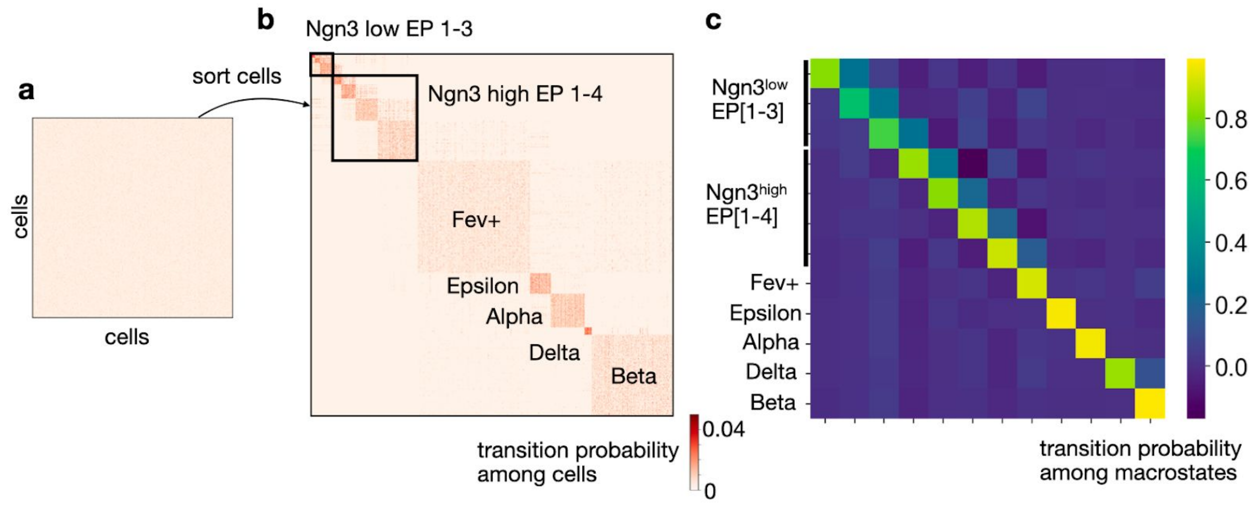


**Supplementary Fig. 3: Recovering structure in the transition matrix**

**a.** Heatmap of the transition matrix for the pancreas dataset from Fig. 2. The ordering of cells (rows and columns) in the matrix is arbitrary. The colorbar has been adjusted such that values larger than the 90th percentile are clipped to the 90th percentile to avoid skewing the colorbar towards extreme values. However, there is still no visible structure in the matrix because of sparsity, noise and the random order of cells. **b.** Same matrix as in (**a**), just re-ordered such that cells which likely belong to the same macrostate are next to each other. This recovers the structure of the developmental dynamics. Note that the sparsity structure of the matrix is symmetric (KNN graph is symmetric) while the actual values are not (RNA velocity infused directionality). **c**. Coarse-grained transition matrix from Fig. 2. Macrostates defined in this matrix were used to reorder cells in (**b**).

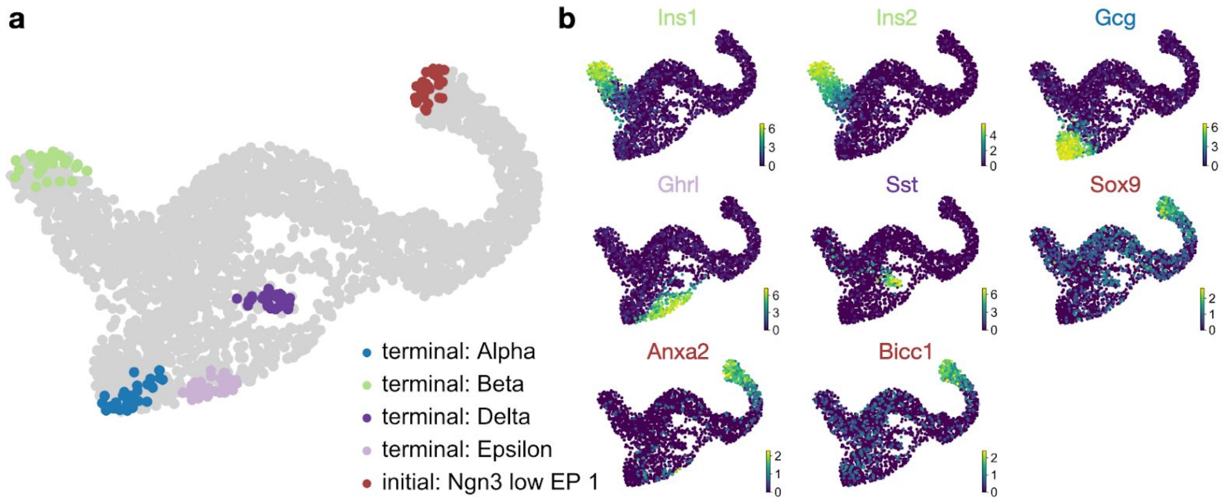


**Supplementary Fig. 4: Marker genes confirm CellRanks initial and terminal state annotations in the pancreas data**

**a.** CellRanks initial and terminal states from Fig. 2 **b.** Cells are colored based on the expression level of the indicated gene in each UMAP. The terminal states express the key marker genes relevant for each respective cell type. Showing for beta: Ins1 and Ins2 (insulin), alpha: Gcg (glucagon), epsilon: Ghrl (ghrelin), delta: Sst (somatostatin)[38]. For the initial state, we show expression of ductal cell markers Sox9, Anxa2 and Bicc1[29,38].

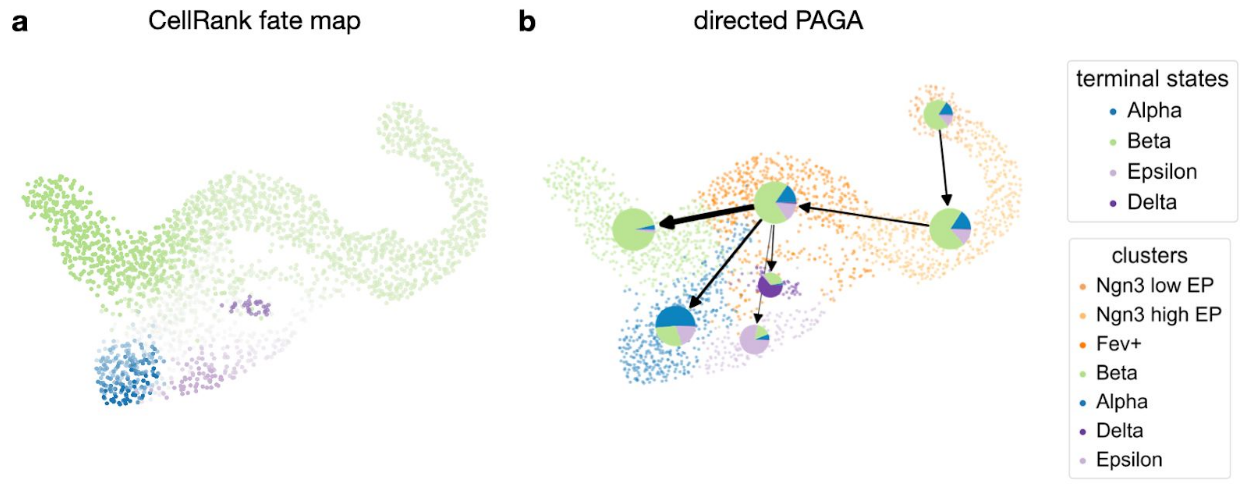


**Supplementary Fig. 5: Visualising fate probabilities in a new directed PAGA graph**

**a.** Lineage probabilities for the pancreas data from Fig. 2, visualised as a fate map where each cell is colored according to its most likely fate. Color intensity reflects the degree of lineage priming. **b.** Probabilistic approximate graph abstraction (PAGA)[11] in a new directed flavour, combined with CellRank's lineage probabilities, shown as pie charts. Arrows represent aggregated velocity flow (Online Methods).

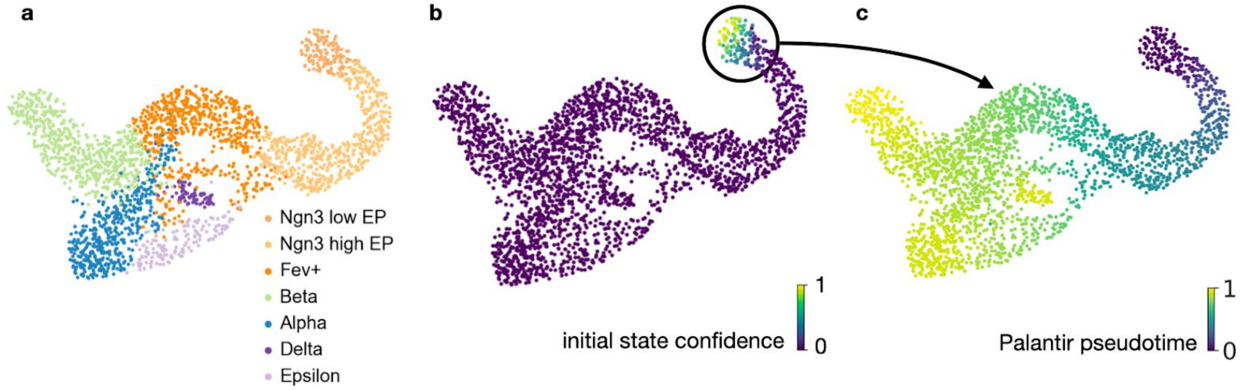


**Supplementary Fig. 6: Palantir pseudotime for the pancreas data**

**a.** UMAP of the pancreas data with cluster annotations from the original publication[29]. **b.** Membership vector corresponding to the Ngn3$^{low}$ EP_1 macrostate which we identified as an initial state. **c.** We selected one of the cells which had high initial state confidence in (**b**) and supplied it to Palantir to compute a pseudotemporal ordering of all cells[25].

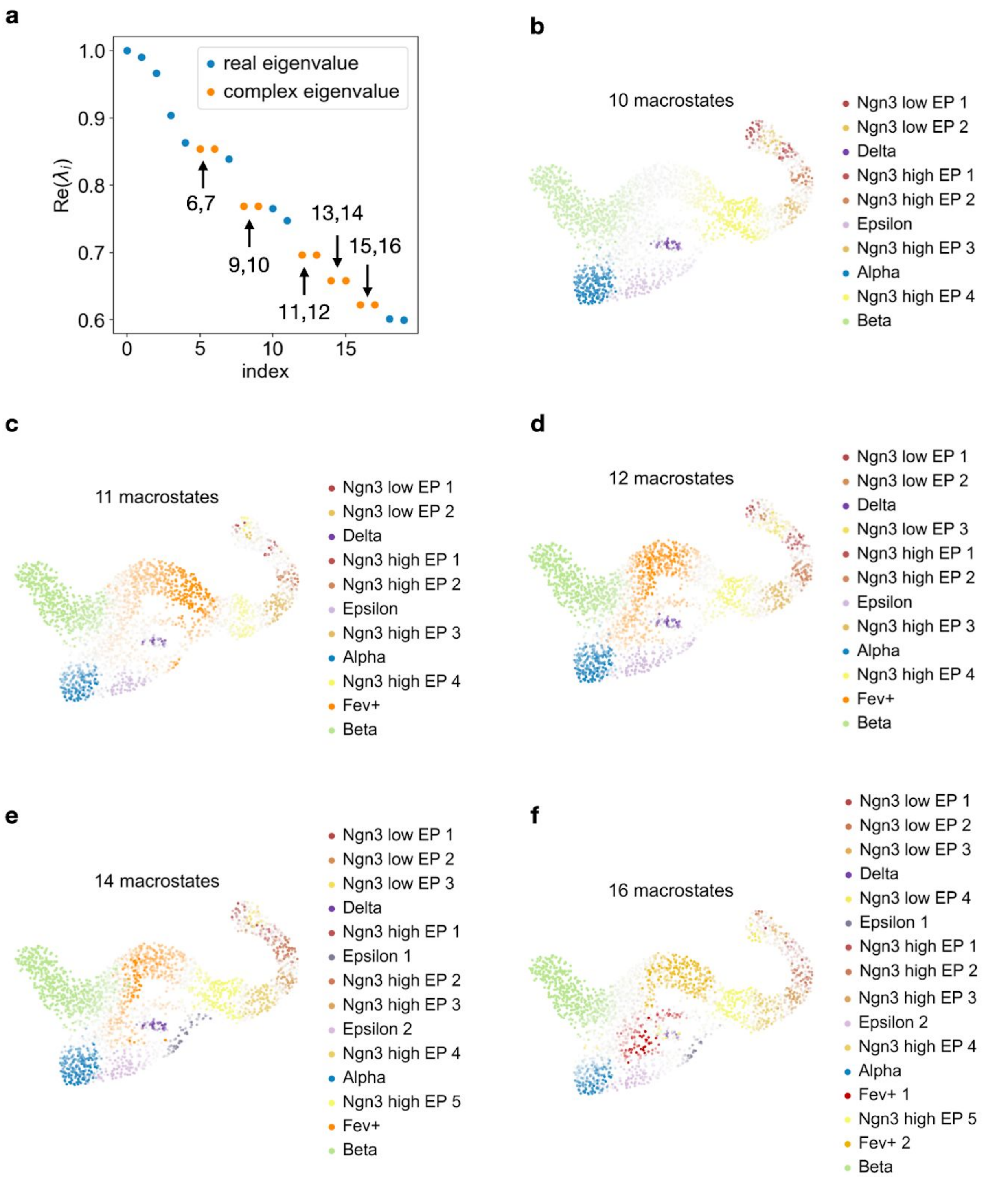


**Supplementary Fig. 7: Varying the number of macrostates for the pancreas does not change biological interpretation**

**a.** Real part of the 20 eigenvalues with the largest real part for the pancreas transition matrix of Fig. 2. We highlight eigenvalues that come in pairs of complex conjugates. Splitting pairs of complex conjugates leads to non-invariant subspaces (Online Methods), therefore, we choose a number of states which always includes both eigenvalues from pairs of complex conjugates. **b-f.** When varying the number of macrostates, we consistently recover the alpha, beta, epsilon, delta and Ngn3$^{low}$ EP_1 states. Increasing the number of macrostates increases the resolution at which we interpret the data. However, for the findings we report in Fig. 2, any of the number of macrostates presented here would lead to similar results and near identical biological interpretation.

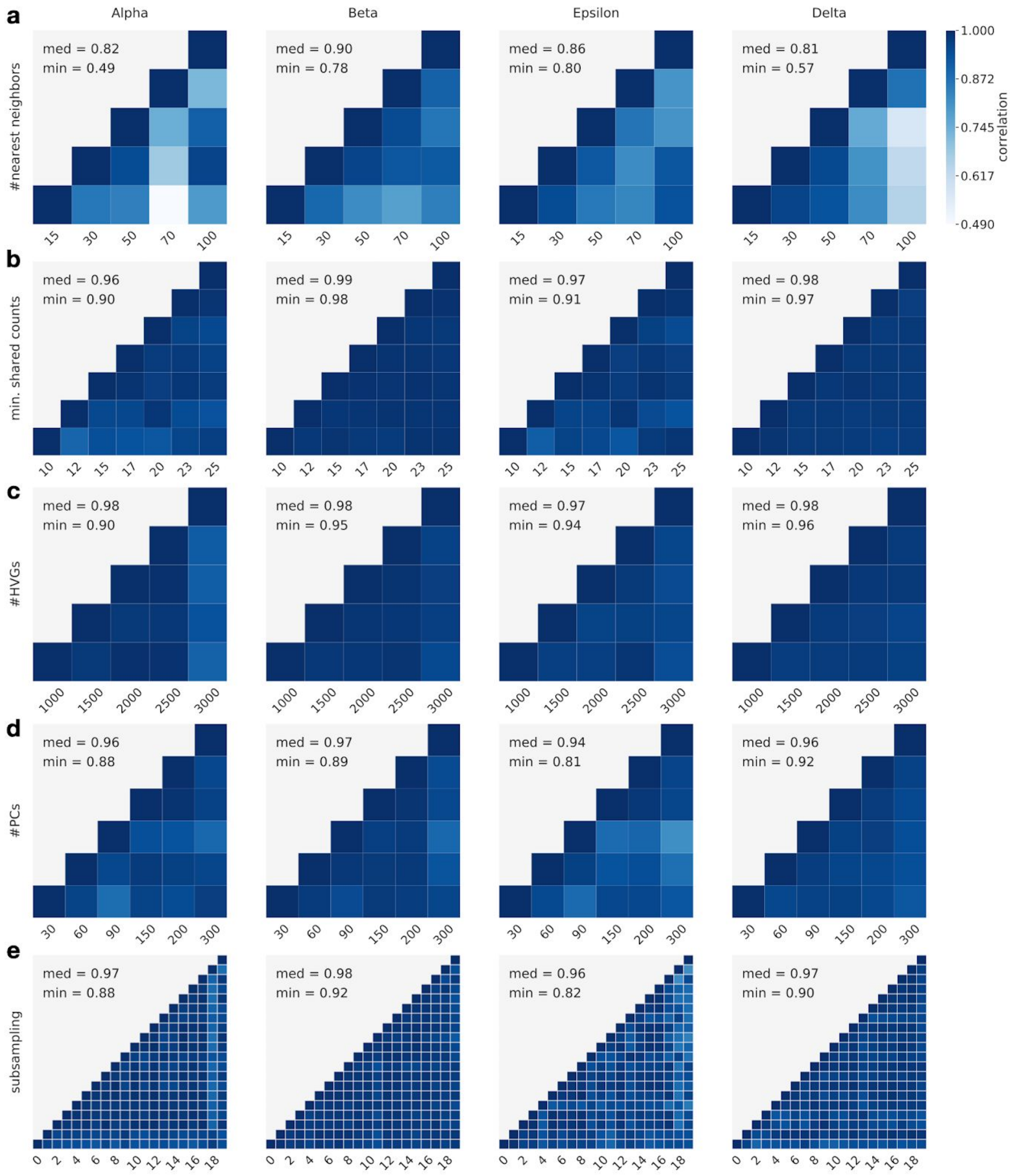


**Supplementary Fig. 8: CellRank is robust to parameter choice and random subsampling**

**a-d** Pairwise correlations of fate probabilities per lineage when varying (**a**) the number of nearest neighbors in KNN graph construction, (**b**) the gene filtering parameter “min_shared_counts” which determines the minimum required number of spliced and unspliced counts, (**c**) the number of highly variable genes, (**d**) the number of principal components used for KNN graph construction. Across the 4 parameters, we achieve a minimum median correlation of 0.81, highlighting CellRanks robustness to a wide range of parameter choices. **e.** Pairwise correlations of fate probabilities per lineage when randomly subsampling the data to 90% of cells. CellRank is very robust to subsampling with a minimum median correlation of 0.96.

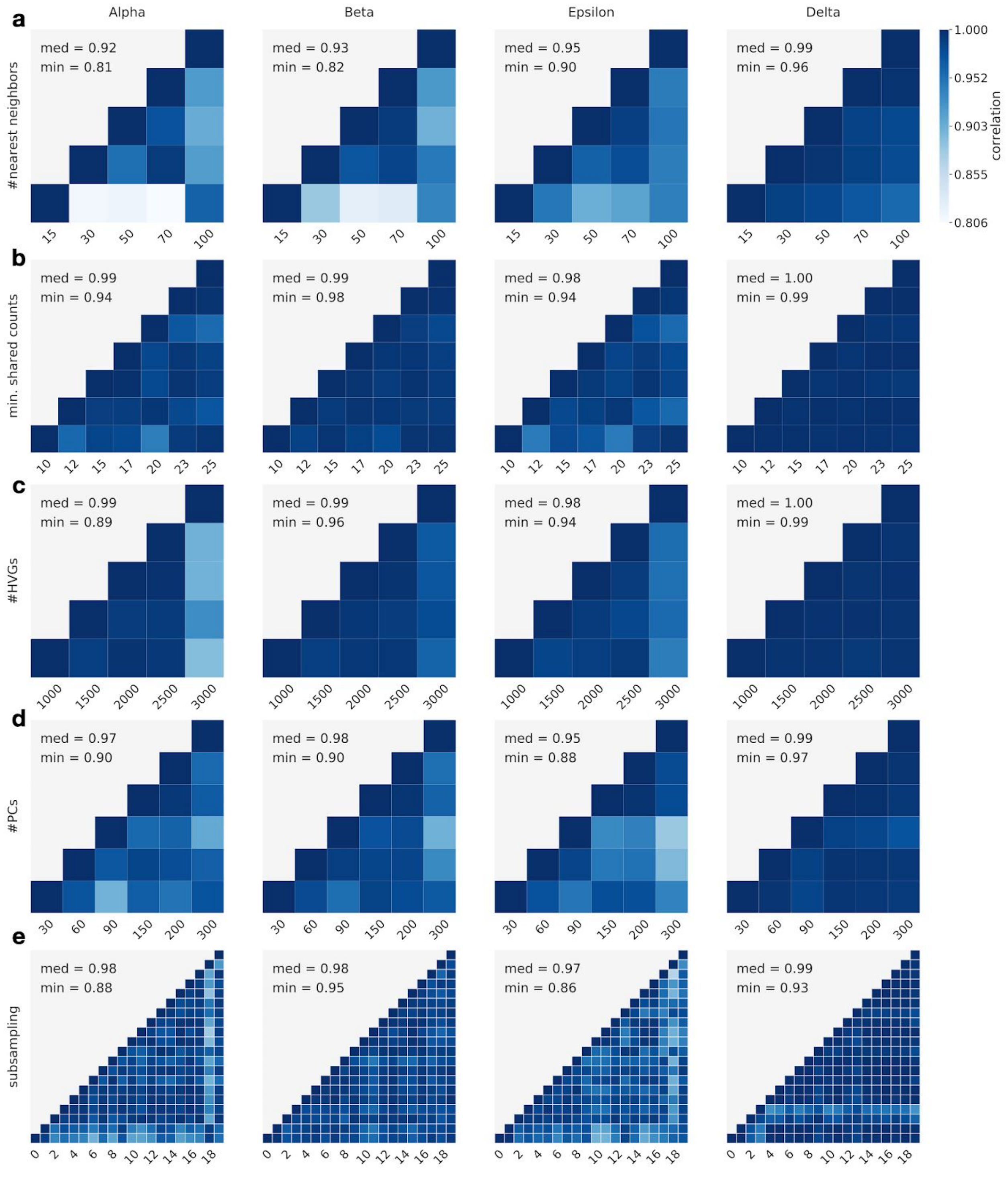


**Supplementary Fig. 9: Fixing the terminal states further increases robustness**

**a-e** Like Suppl. Fig. 8, only that we fix the terminal states to restrict the robustness comparison to the computation of fate probabilities, i.e. the terminal states have been computed once and were fixed across all parameter variations and subsampling of cells. Note that the color scale changed. This increases the minimum median correlation for parameter variations (**a-d**) from 0.81 with recomputed terminal states to 0.92 here and for subsampling (**e**) from 0.96 with recomputed terminal states to 0.97 here.

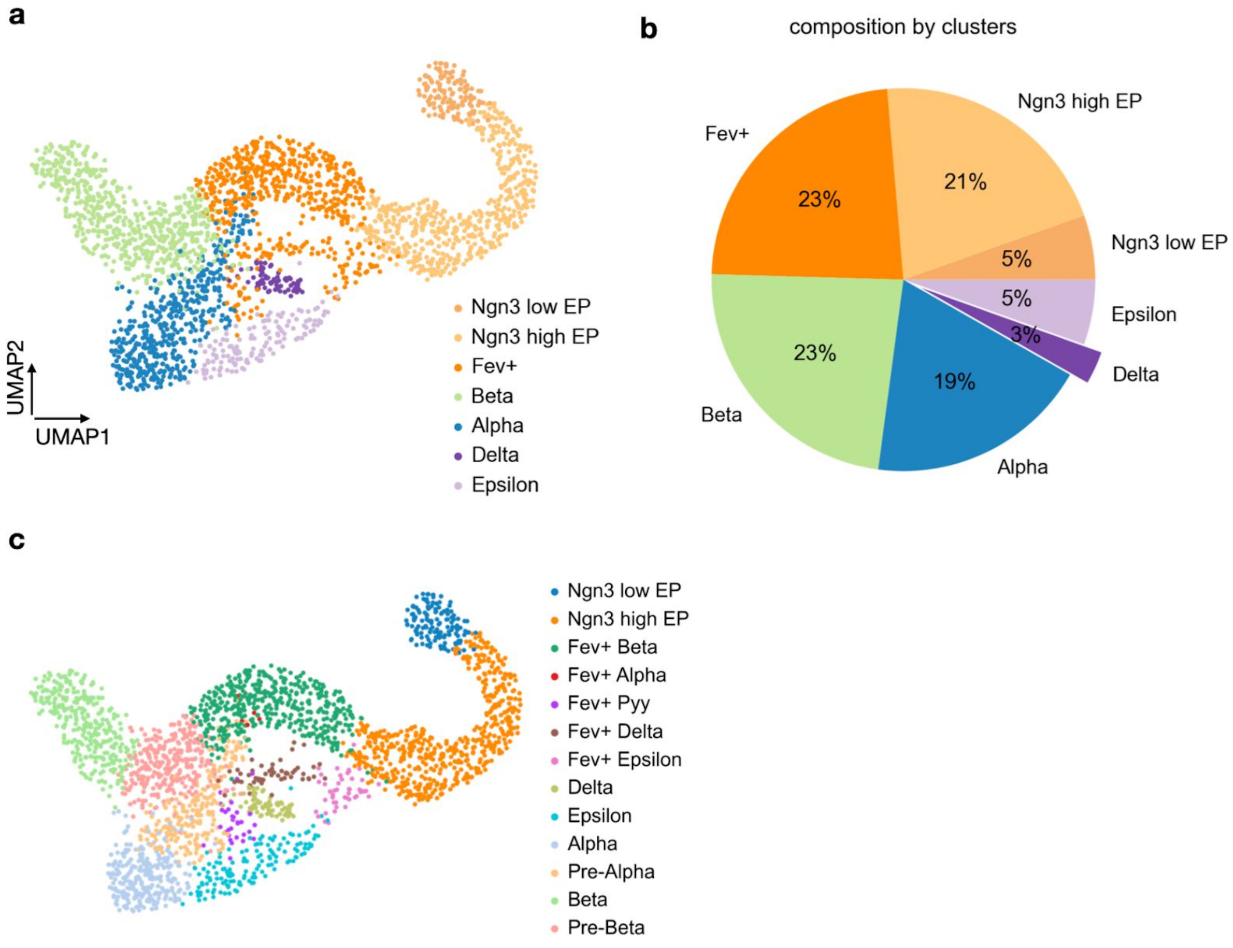


**Supplementary Fig. 10: Cell type proportions in the pancreas data**

**a.** UMAP representation of the pancreas data from Fig. 2 with original cluster annotations[29]. **b.** Cell type proportions. Delta cells are the rarest cell type in this data with only 3% abundance. **c.** Sub-clustering of the data from the original publication[29]. Alpha and beta cells have been sub-clustered by us.

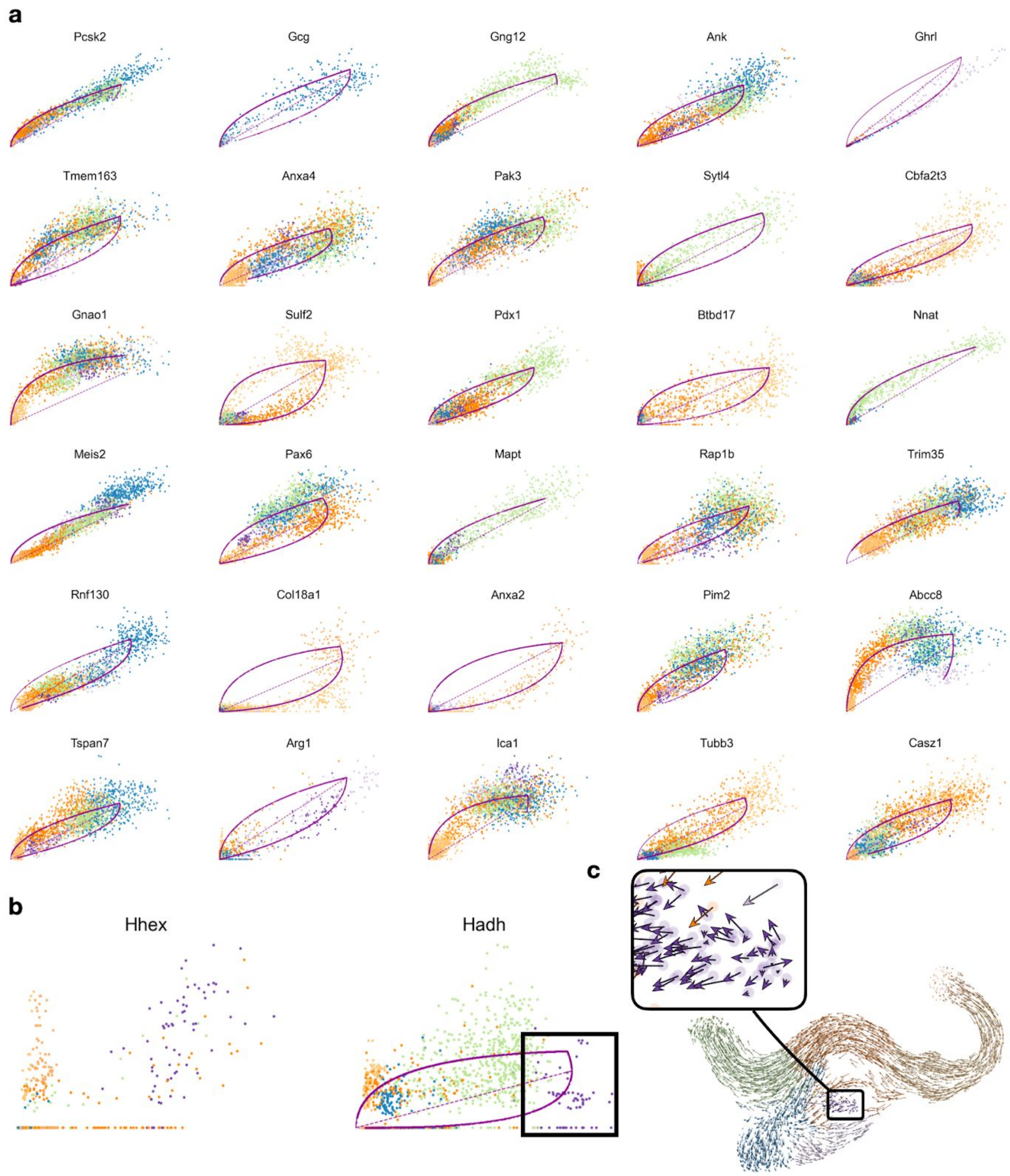


**Supplementary Fig. 11: Splicing kinetics do not capture delta cell development**

**a.** Phase portraits of the top 30 genes which are assigned the highest likelihoods by scVelo's dynamical model of the mRNA lifecycle. Unspliced counts are on the x-axis, spliced counts are on the y-axis. Cells are colored according to the clusters from Fig. 2. The solid purple curve is scVelo's dynamical fit and the dashed purple line is scVelo's inferred steady-state ratio. The top 30 genes are dominated by drivers for the alpha (*Gcg*[38]), epsilon (*Ghrl*[38]) and beta (*Gng12*[89], *Pdx1*[40,87]) lineages while delta drivers are not present. **b.** *Hhex*[41], which is the major known delta-driver, could not be fit by scVelo because of too little expression and too large noise levels. *Hadh*, another likely delta driver, could be fit. However, delta cells are an outlier in this fit (see inlet) and were not correctly assigned to the steady-state. *Cd24a*, another likely delta driver, did not make it through scVelo's filters to be considered a "velocity gene", possibly because of too little expression and too high noise levels. **c.** UMAP with projected scVelo velocity vectors, inlet highlights delta cells and their noisy velocity vectors. Splicing kinetics do not reveal development towards this state.

a

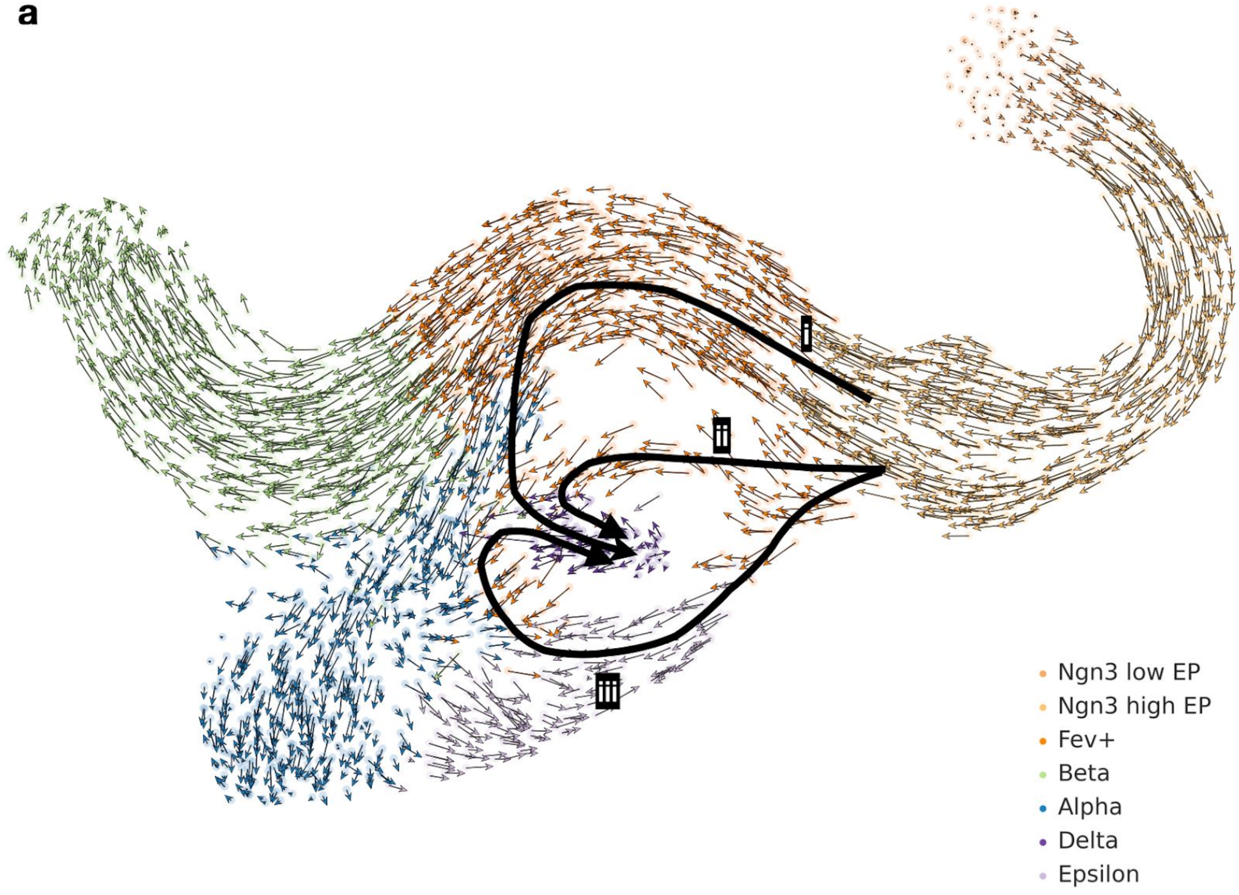


**Supplementary Fig. 12: Projected velocities do not reveal how delta cells are generated**

**a.** scVelo velocities projected onto the UMAP show three possible paths for delta cell differentiation (i-iii). Velocities reveal short-range fate relationships but cannot be combined to give long-range fate predictions from looking at an embedding.

**a**

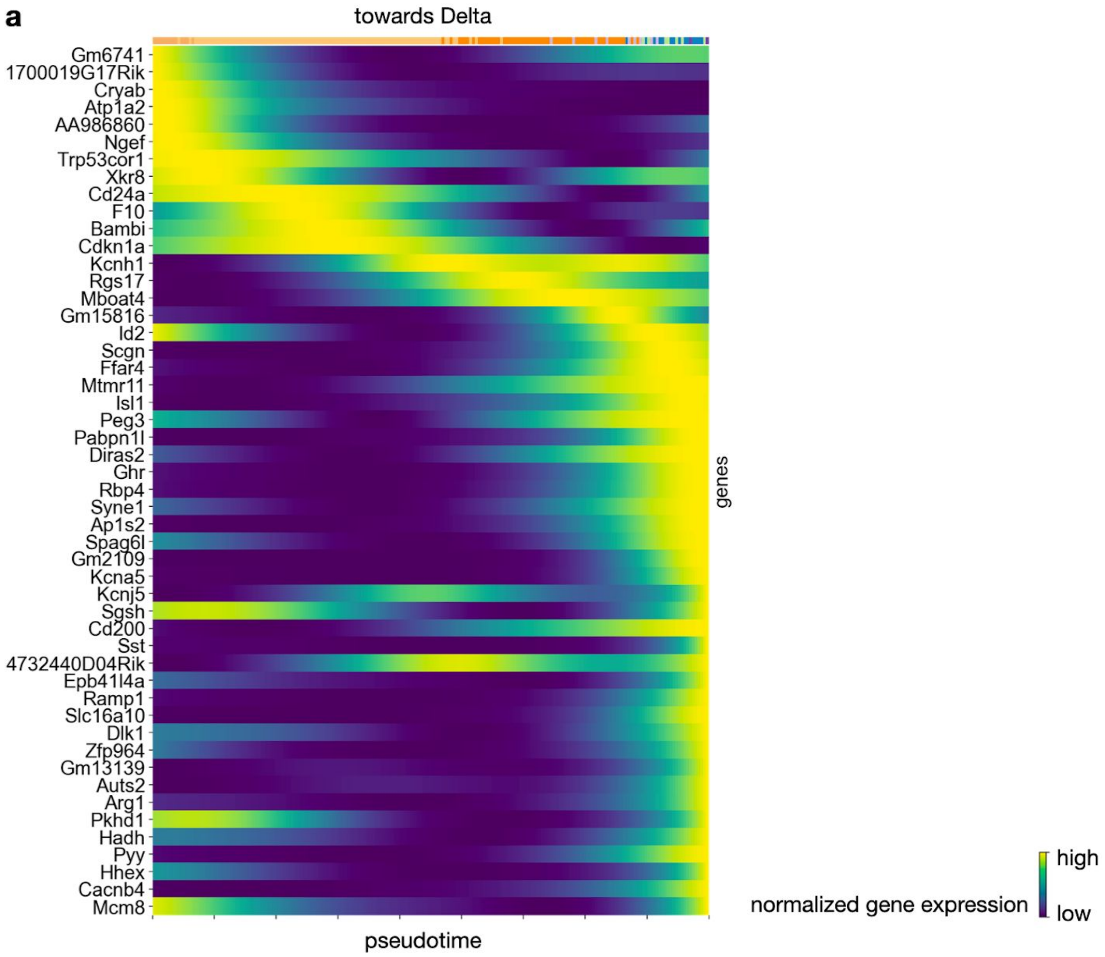


**Supplementary Fig. 13: Heatmap of genes whose expression correlates well with delta fate**

**a.** Heatmap of Fig. 3d with all gene names shown.

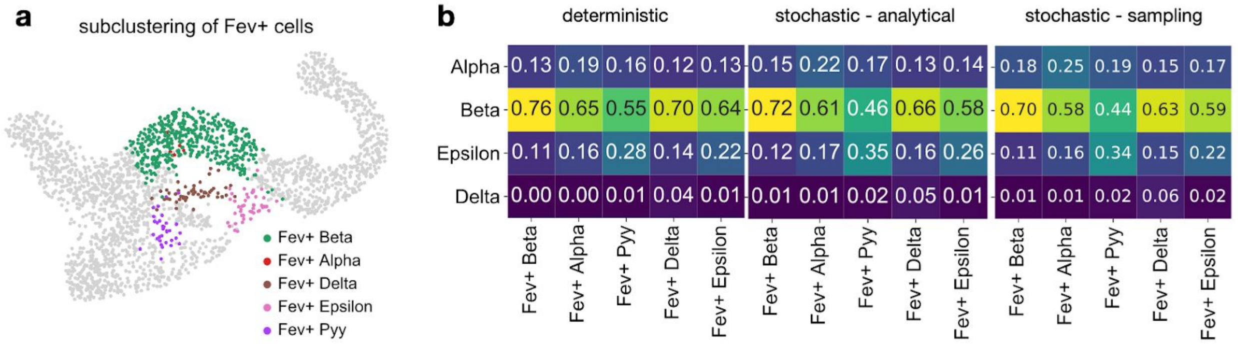


**Supplementary Fig. 14: Comparing fate probabilities between deterministic and stochastic modes**

**a.** Sub-clustering of the Fev+ cluster in the pancreas data[29]. **b.** Comparing average fate probabilities per sub-cluster. These were obtained from not propagating ("deterministic") or propagating ("stochastic - analytical" and "stochastic - sampling") velocity uncertainty. Both stochastic approaches agree in down-weighting probability towards the dominant beta fate and up-weighting probability towards the alpha, delta and epsilon fates.

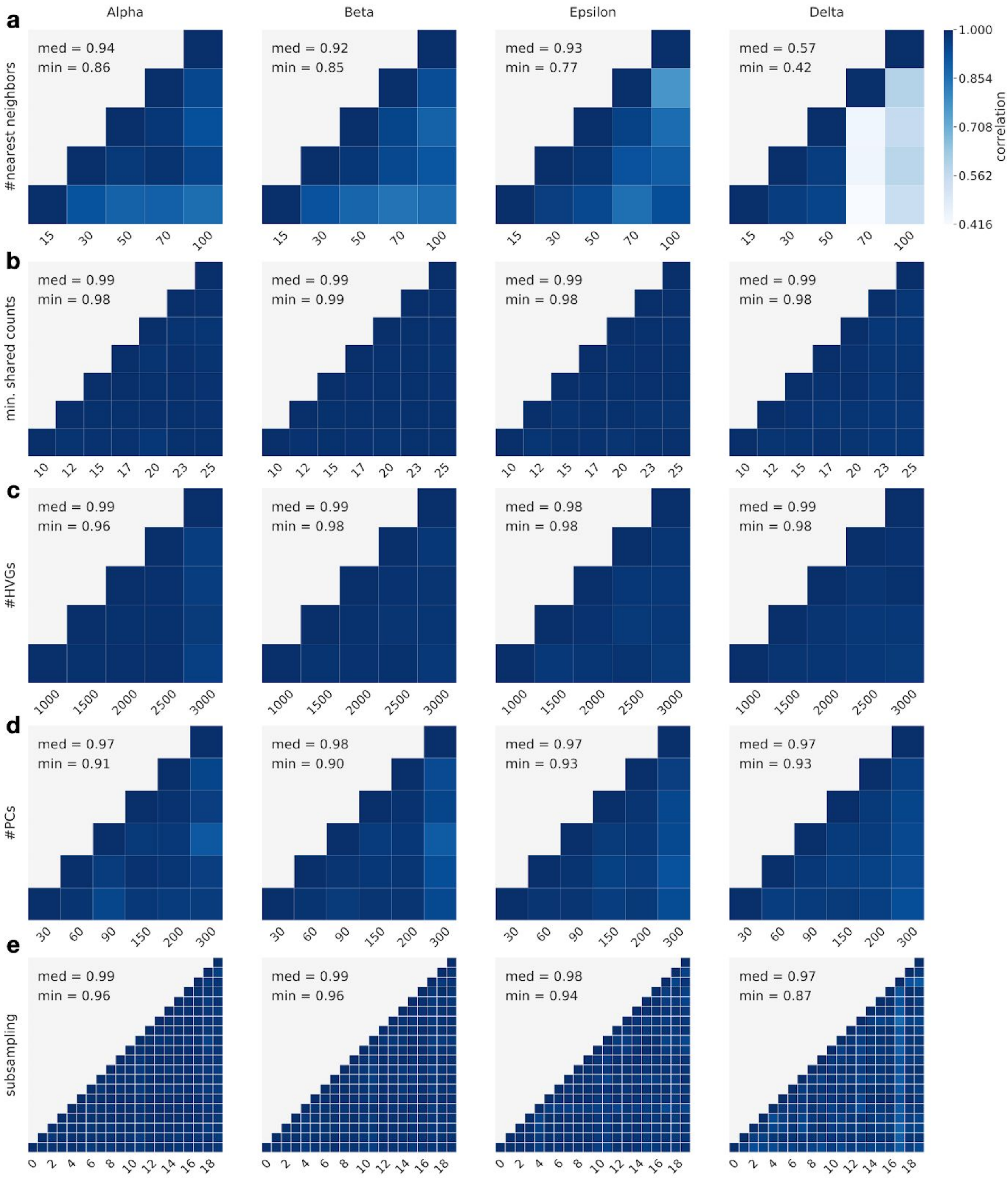


**Supplementary Fig. 15: Robustness increases when propagating uncertainty**

**a-e.** Like Suppl. Fig. 8, only that we use the analytical approximation to propagate uncertainty into the transition probabilities. Apart from the delta lineage for the number of nearest neighbors, this increases the minimum median correlation for parameter variations (**a-d)** from 0.81 in the deterministic case to 0.92 here. When we vary the number of nearest neighbors for the delta lineage, we observe outlier terminal states when using 70 nearest neighbors. This effect disappears when we fix the terminal states (Suppl. Fig. 16a). **e.** For subsampling, using the stochastic approximation increases the minimum median correlation from 0.96 in the deterministic case to 0.97 here.

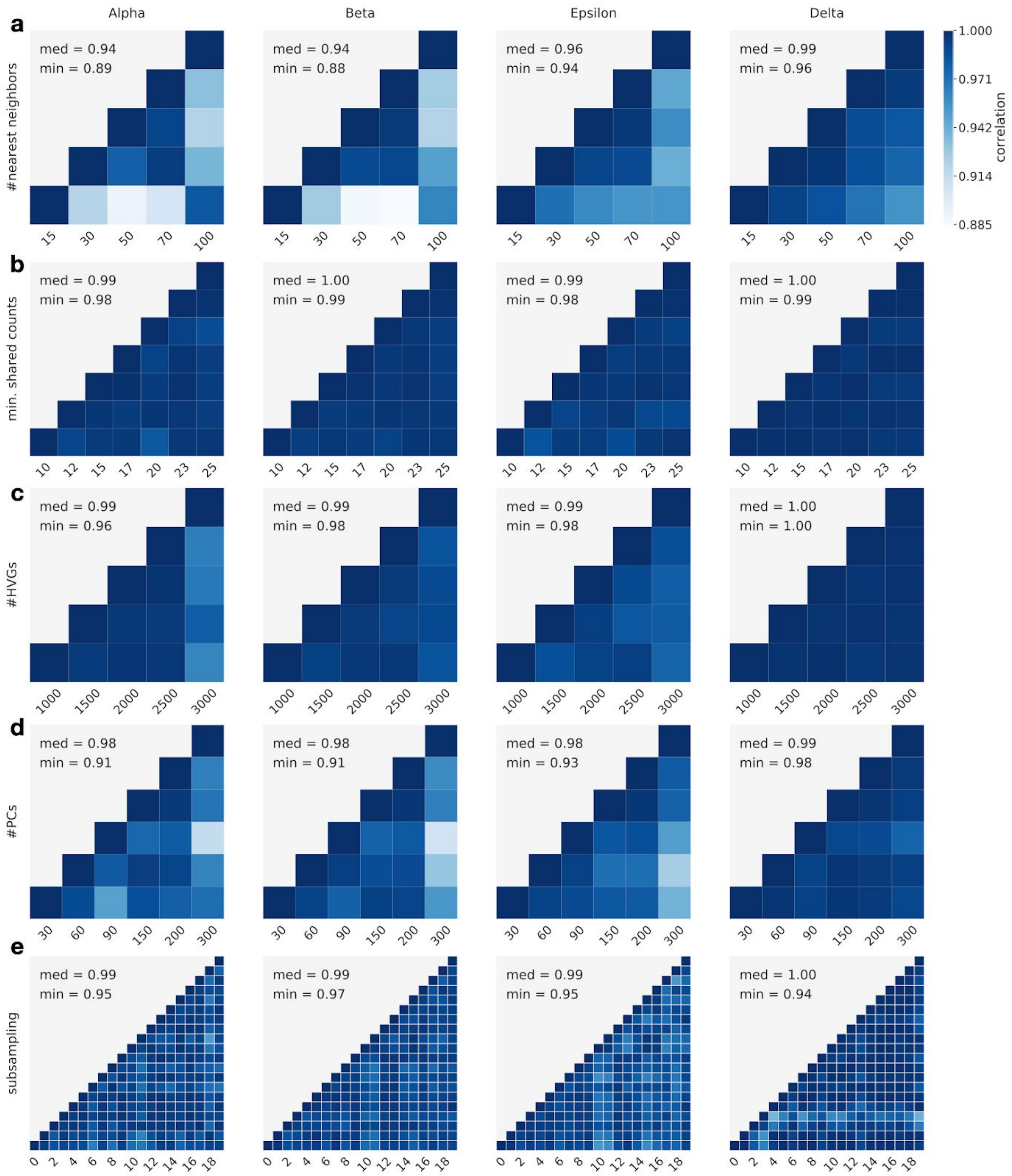


**Supplementary Fig. 16: Robustness further increases when propagating uncertainty and fixing the terminal states**

**a-e.** Like Suppl. Fig. 15, only that we fix the terminal states to restrict the robustness comparison to the computation of fate probabilities, i.e. the terminal states have been computed once and were fixed across all parameter perturbations and subsampling of cells. This increases the minimum median correlation for parameter variations (**a-d)** from 0.57 with recomputed terminal states to 0.94 here and for subsampling (**e)** from 0.97 with recomputed terminal states to 0.99 here.

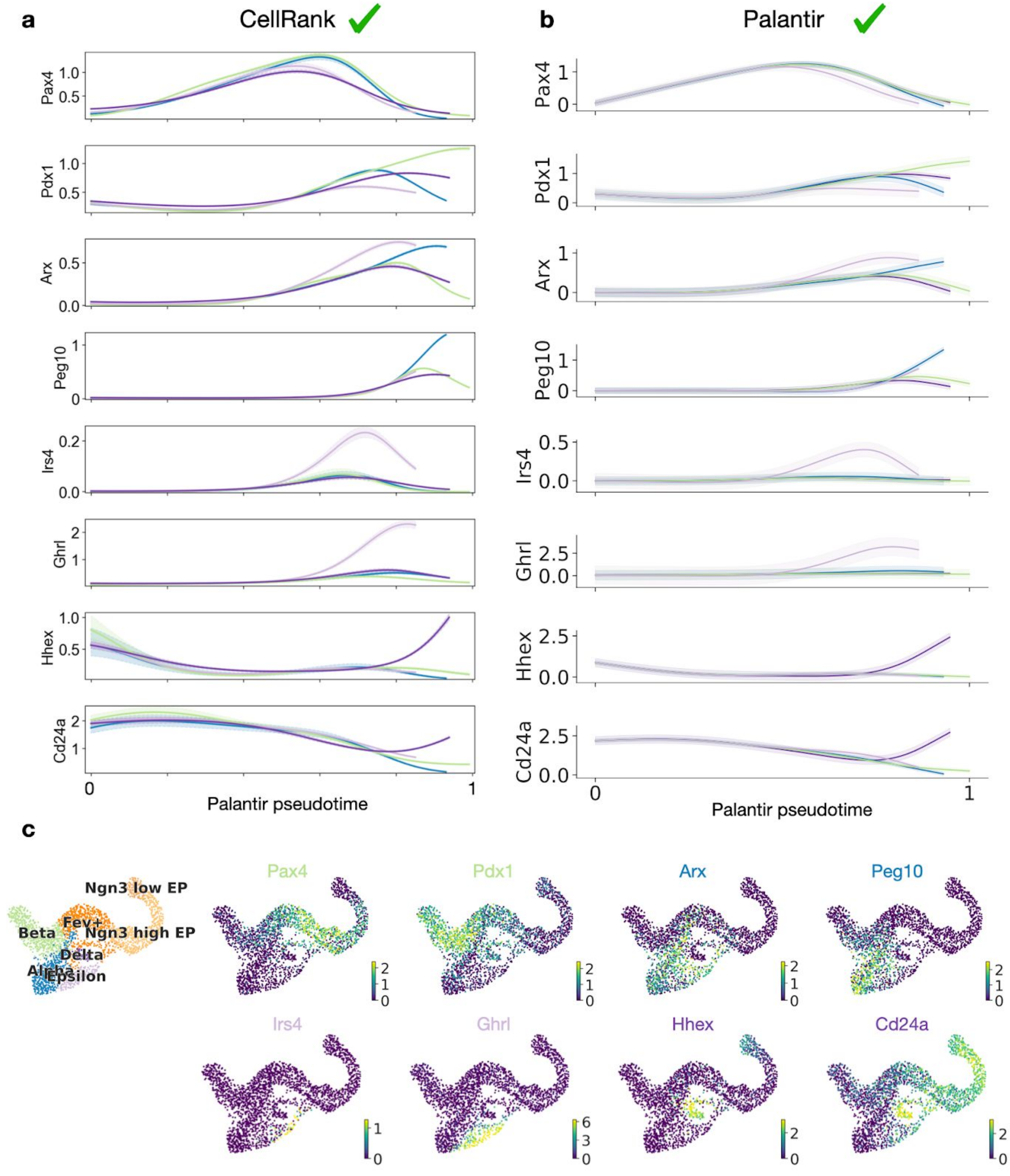


**Supplementary Fig. 17: Gene expression trends for CellRank and Palantir**

**a-b.** Gene expression trends for key regulators *Pax4*[39] and *Pdx1*[40,87,88] (beta), *Arx*[39] (alpha), *Ghrl*[38] (epsilon), *Hhex*[41] and *Cd24a*[44,45] (delta) as well as lineage associated genes *Peg10*[42,89] (alpha) and *Irs4*[42] (epsilon) for CellRank (**a**) and Palantir (**b**). The x-axis is given by Palantir's pseudotime (Suppl. Fig. 6c). Expression values were imputed using MAGIC[85]. Green ticks indicate that methods correctly predicted lineage-specific gene regulation. CellRank and Palantir give similar results because both methods were supplied with CellRank's terminal states. **c.** Cluster labels from ref.[29] as well as expression of the genes from (**a**) and (**b**) in the UMAP.

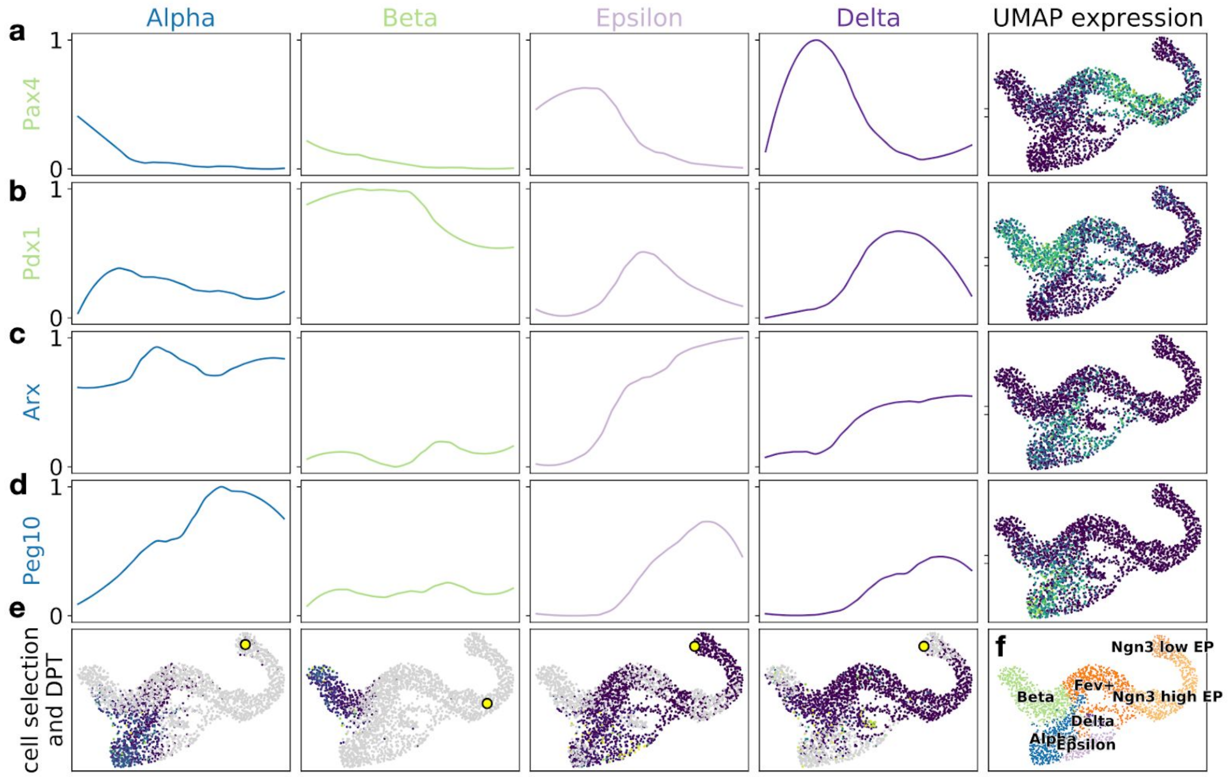


**Supplementary Fig. 18: FateID gene expression trends for alpha- and beta-fate associated genes**

**a-d.** Gene expression trends for key regulators *Pax4*[39] and *Pdx1*[40,87,88] (beta), *Arx*[39] (alpha) as well as the lineage associated gene *Peg10*[42,89] (alpha). We color each gene by its associated lineage. Expression trends computed using FateID towards the alpha, beta, epsilon and delta fates for these genes are shown in the first four columns. On the x-axis are the indices of the pseudo-temporally ordered cells assigned to the given lineage, on the y-axis is normalised gene expression. The line represents a local regression of z-transformed gene expression values (Online methods). In the last column, we show gene expression values in the UMAP. Yellow denotes high expression, blue denotes low expression. **e.** For each lineage, we show the cells assigned to it by FateID, colored by diffusion pseudotime[10] (DPT) which was used for gene-trend smoothing (Online methods). The yellow dot denotes the root cell used for DPT computation in the corresponding lineage. Cells not assigned to a lineage are colored in grey. **f.** Cluster annotations from the original publication[29].

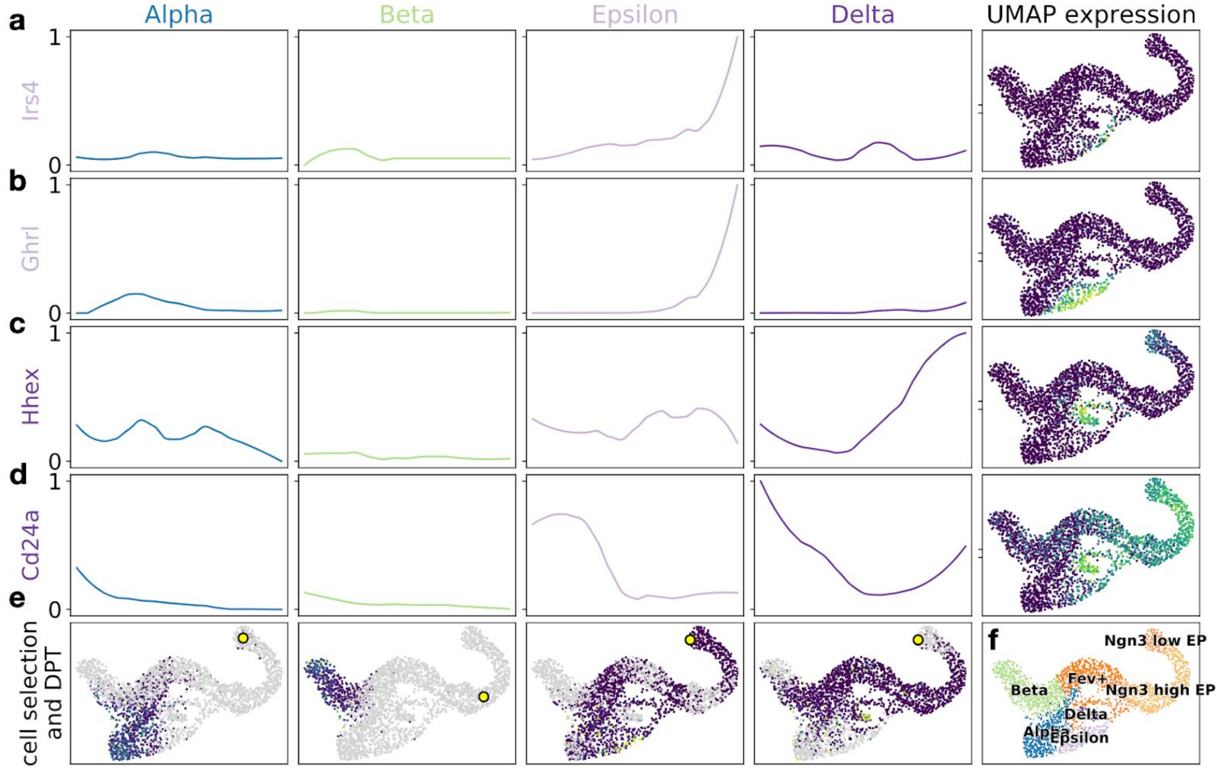


**Supplementary Fig. 19: FateID gene expression trends for epsilon- and delta-fate associated genes**

**a-d.** Like Suppl. Fig. 18, only that we show trends for the key lineage drivers *Ghrl*[38] (epsilon), *Hhex*[41] and *Cd24a*[44,45] (delta) as well as for the lineage associated gene *Irs4*[42] (epsilon). Panels **e** and **f** remain unchanged.

**a**

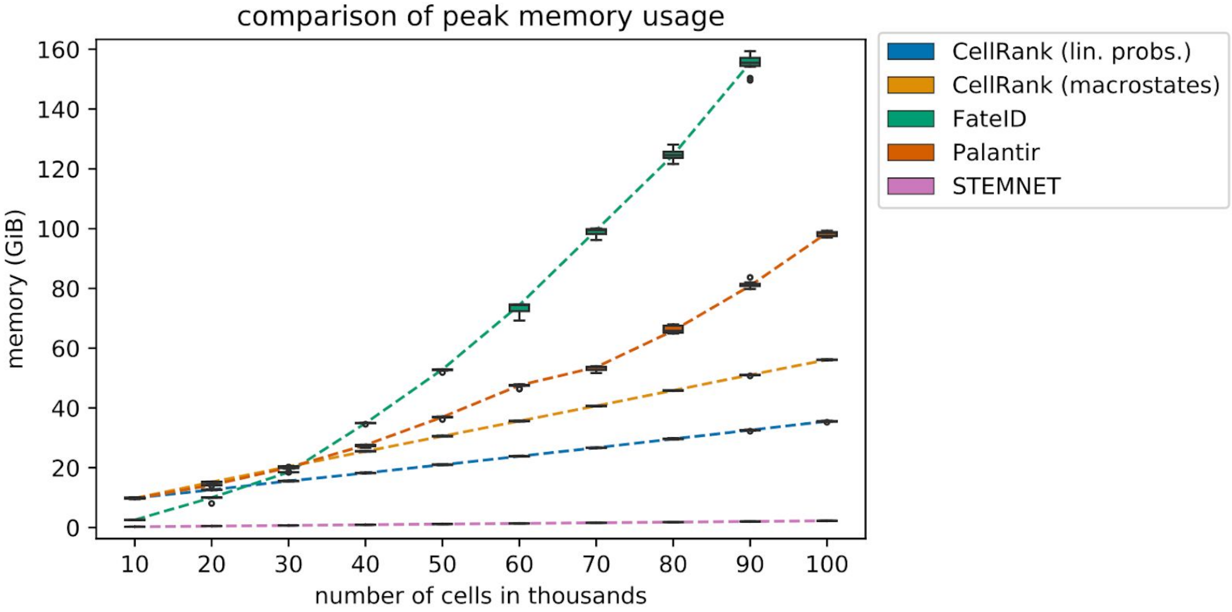


**Supplementary Fig. 20: Comparing peak memory usage across methods**

**a.** Boxplot comparing peak memory usage of CellRank to compute macrostates and fate probabilities with FateID, Palantir and STEMNET to compute fate probabilities, given CellRank's terminal states on the reprogramming dataset[63] of Fig. 5f (Online methods). Box plots show the median, the box covers the 25 to 75% quantiles, whiskers extend up to 1.5 times the interquartile range above and below the box. Outliers are shown as dots and the dashed lines connect the medians. FateID did not finish on 100k cells because of memory constraints. Note that parallelisation across 32 cores increases peak memory usage for Palantir and CellRank, the only two methods that make use of parallelisation. We report decreased peak memory usage on 100k cells using a single core for Palantir and CellRank in Supplementary Table 3.

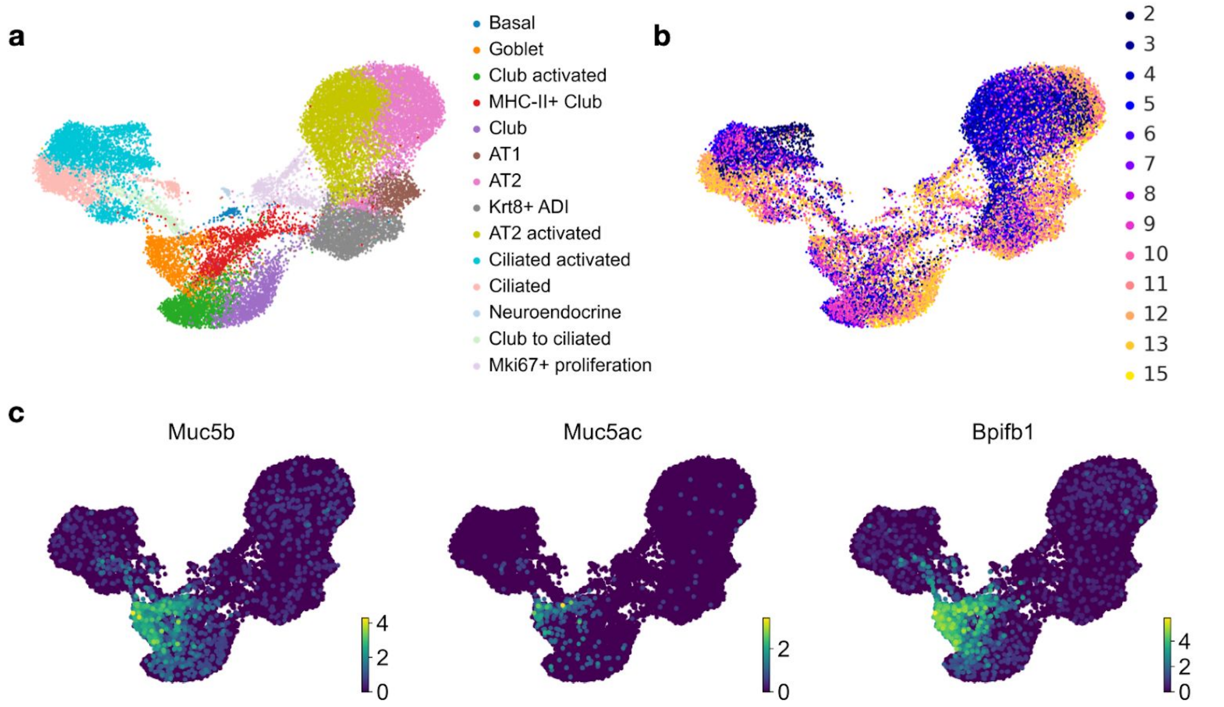


**Supplementary Fig. 21: Cluster labels and time point annotations for the lung data**

**a.** Original cluster labels for the lung regeneration data[64] in a UMAP projection. The data contains 24,051 murine lung epithelial cells sequenced using the Dropseq workflow[90] at 13 time points spanning days 2-15 past bleomycin injury. The 'activated' label refers to cell states that emerge after bleomycin injury. **b.** Same as (**a**) with time points colored in. Time points refer to time passed since bleomycin injury. **c.** Expression of goblet cell markers *Muc5b, Muc5ac and Bpifb1* agrees with the goblet annotation of (**a**).

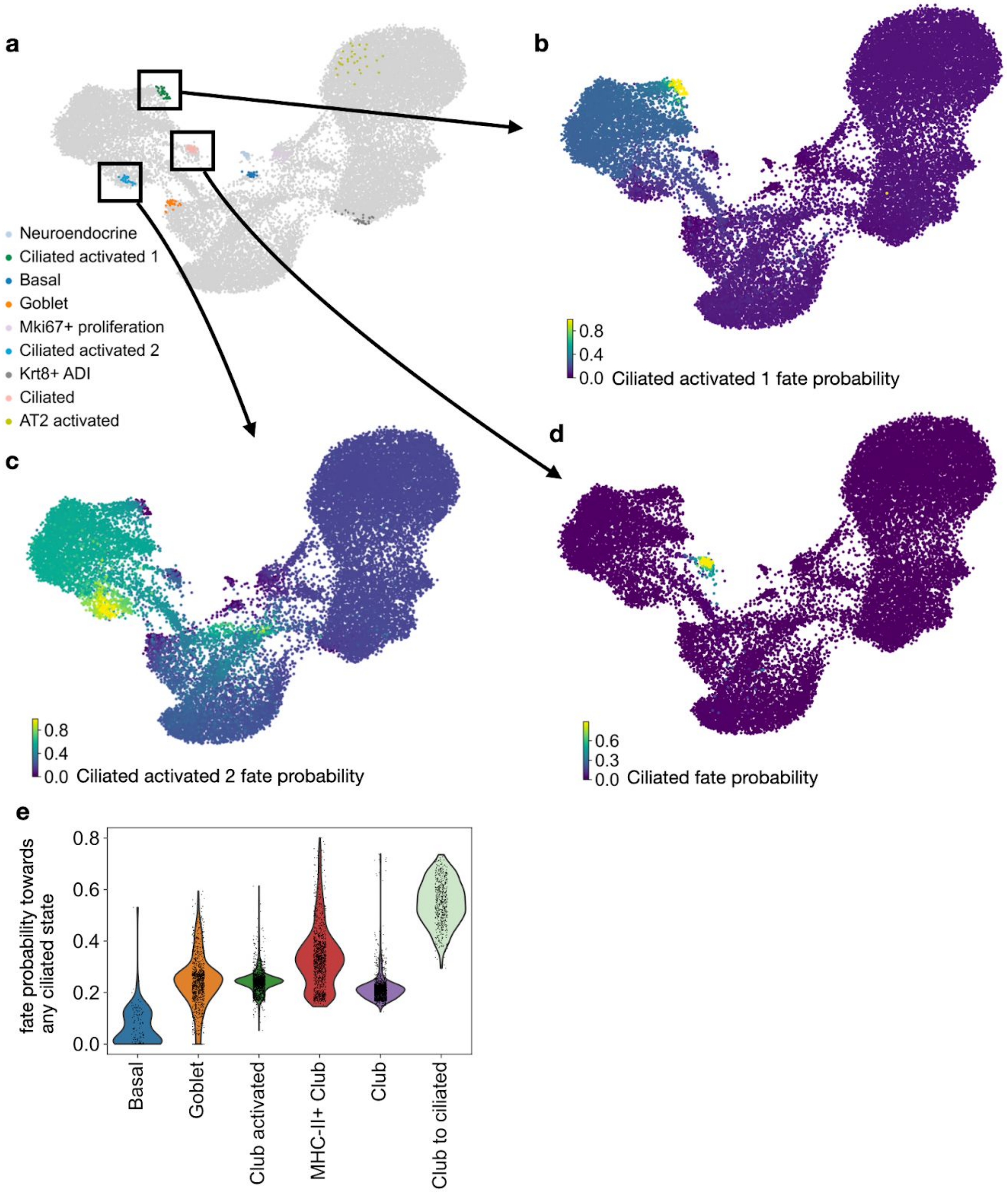


**Supplementary Fig. 22: CellRank correctly recovers club to ciliated trajectory**

**a.** CellRank identified macrostates in the UMAP. For each macrostate, we show the 30 most confidently assigned cells. We color macrostates according to the cluster from Suppl. Fig. 21a that they mostly overlap with. We highlight 3 macrostates that overlap with the ciliated clusters:

the ‘Ciliated activated 1’, ‘Ciliated activated 2’ and ‘Ciliated’ macrostates. Note that the ‘activated’ label has been assigned in the original publication to denote populations that appear upon injury[64]. **b-d.** Same UMAP as in (**a**), colored by fate probabilities towards the Ciliated activated 1, Ciliated activated 2 and Ciliated macrostates. Among the genes that correlated best with fate probabilities towards the Ciliated macrostate were *Mcidas, Deup1* and *Ccno,* all of which are involved in the normal development of ciliated cells[91–94]. **e.** Violin plots showing the distribution over fate probabilities to transition towards any of the three ciliated states within the club, goblet and basal cell clusters shown in Suppl. Fig. 21a. We summed over fate probabilities towards the three individual ciliated macrostates. The cluster annotated ‘Club to ciliated’ in the original publication[64] is assigned the highest probability by CellRank. Interestingly, goblet cells are assigned a small probability to transition towards the ciliated population, an observation that was also made by others recently[95].

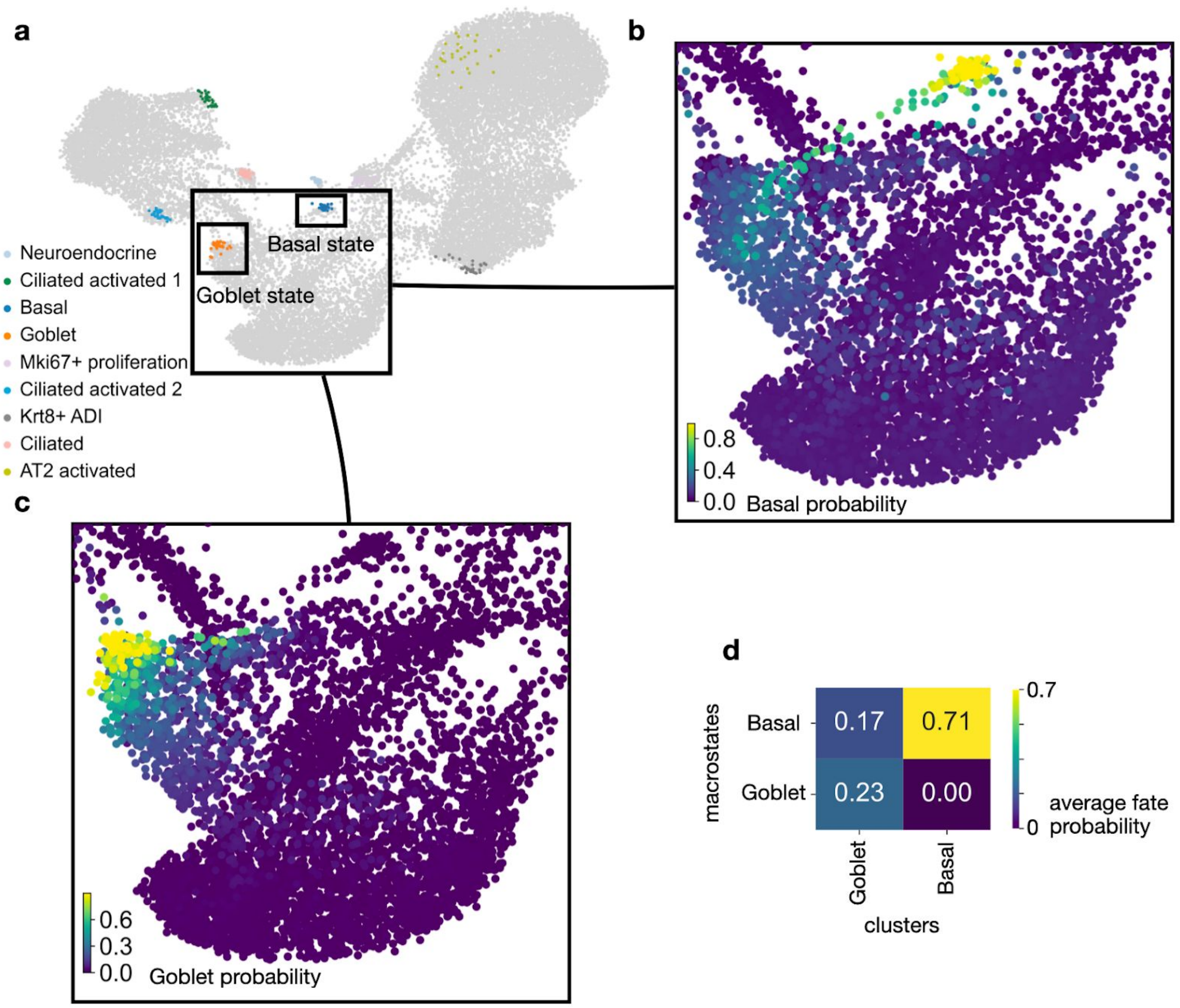


**Supplementary Fig. 23: CellRank predicts a goblet to basal dedifferentiation trajectory**

**a.** 9 macrostates, computed using CellRank. We highlight a subset of airway cells, composed of club, goblet and basal cells. **b.** Single-cell fate probabilities of transitioning towards the basal state. We see a 'band' of cells within the goblet cluster which has high basal probability. **d.** Single-cell fate probabilities of transitioning towards the goblet state. Basal cells do not show any probability of transitioning towards the goblet state. **e.** Quantification of the results from (**c**) and (**d**). Goblet cells have a large probability of transitioning towards basal cells, but basal cells have no probability of transitioning towards the goblet state. This confirms that the direction of the recovered trajectory is goblet -> basal.

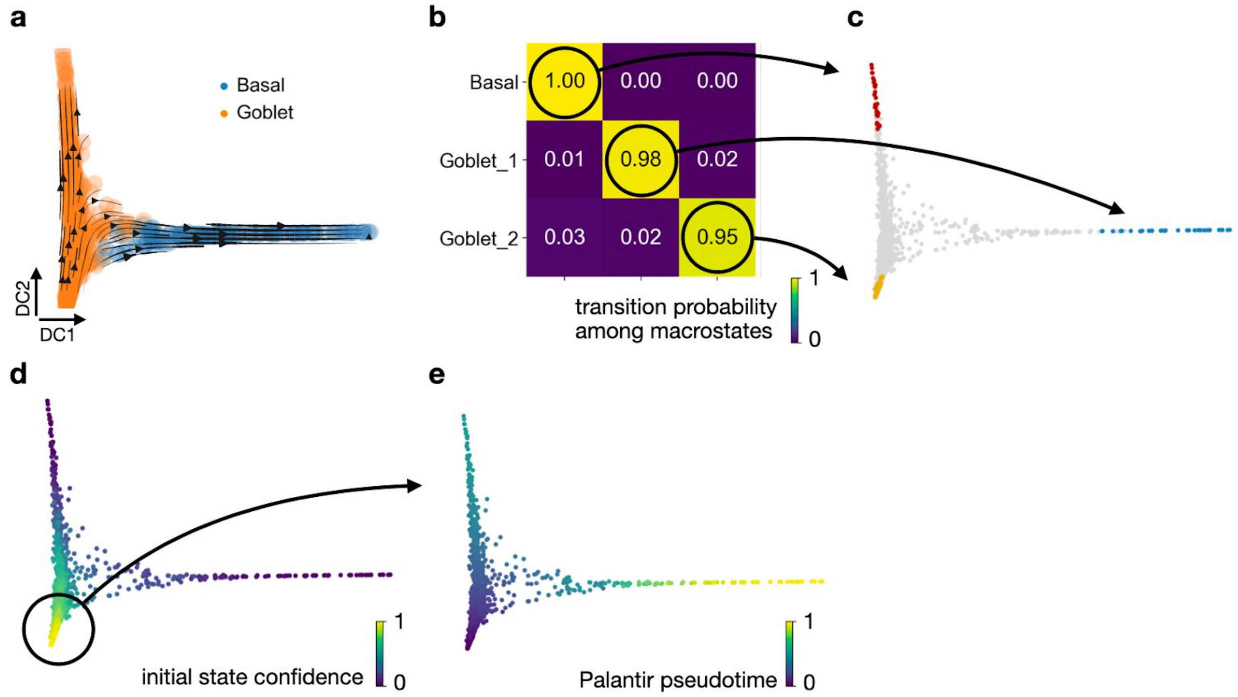


**Supplementary Fig. 24: Computing a pseudotime for the goblet to basal transitions**

**a.** Diffusion map of a subset of the cells from the lung data of Fig. 6 labelled as "Goblet" and "Basal" in the original publication[64]. **b** Coarse-grained transition matrix, computed for three macrostates. The macrostate labelled as 'Goblet_2' was automatically detected as initial by CellRank because it had the smallest value in the CGSD. **c.** Showing the 30 cells most confidently assigned to their macrostate in the diffusion map. We kept the color for the basal state but created two new colors for the initial and terminal goblet states because they both overlap with the same transcriptomic goblet cluster and hence would both get the same color. **d.** Membership vector corresponding to the initial 'Goblet_2' state, here labelled as 'initial state confidence'. The cell which had the maximum value in the initial state confidence was used as initial cells to compute Palantir's pseudotime. **e.** Palantir pseudotime.

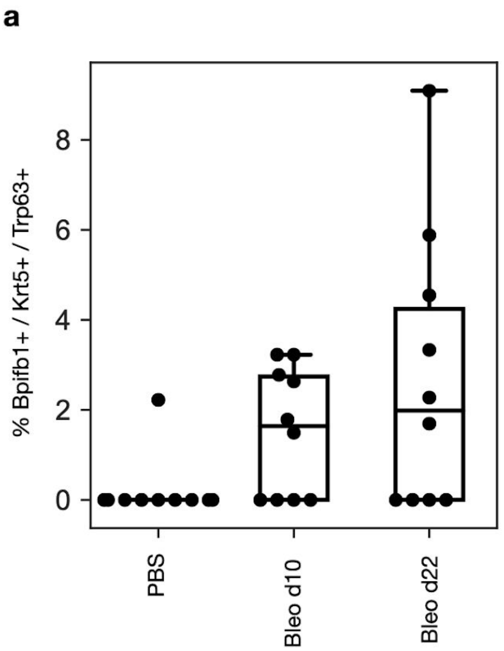


**Supplementary Fig. 25: Quantifying the abundance of triple positive cells**

**a.** We quantify the abundance of cells positive for the goblet cell marker *Bpifb1* as well as the basal cell markers *Krt5* and *Trp63* in the three stages in control mice treated with PBS (n=2), ten days past bleomycin injury (Bleo d10, n=2), and 22 days past bleomycin injury (Bleo d22, n=2) in ten different intrapulmonary airway regions (n=5 per mouse). We find triple positive cells in bleomycin-injured lungs and rarely in PBS treated control mice.
