## Supplementary Tables for "CellRank for directed single-cell fate mapping"

| #cells (thousands) | CR (macro states) [mean] | CR (macro states) [std] | CR (lin. probs.) [mean] | CR (lin. probs.) [std] | P [mean] | P [std] | S [mean] | S [std] | F [mean] | F [std] |
|---|---|---|---|---|---|---|---|---|---|---|
| **10** | 2.8 | 0.9 | 1.5 | 2.2 | 27.3 | 3 | 6.5 | 0.3 | 160.5 | 8.8 |
| **20** | 5 | 0.1 | 4.2 | 0.8 | 102.5 | 10.4 | 11.9 | 0.4 | 531.1 | 31.7 |
| **30** | 7.6 | 0.1 | 8.8 | 0.8 | 217.6 | 19.8 | 17.7 | 0.4 | 1029.6 | 57.8 |
| **40** | 11.1 | 0.5 | 18.3 | 1.6 | 447.7 | 52.6 | 22.9 | 0.3 | 1840.8 | 89.5 |
| **50** | 15 | 0.3 | 32.1 | 2.9 | 750.1 | 66.1 | 28.9 | 0.3 | 2690.5 | 117.5 |
| **60** | 18.8 | 2 | 35.9 | 6.2 | 1092.4 | 119.5 | 35 | 0.6 | 3772.8 | 147 |
| **70** | 21.8 | 0.5 | 48.5 | 1.8 | 1524.1 | 158.9 | 42.1 | 0.5 | 5151.4 | 217.9 |
| **80** | 26.2 | 1.8 | 65.7 | 2.5 | 2352.1 | 292.4 | 48.4 | 0.5 | 7414.3 | 454.8 |
| **90** | 29.7 | 1.3 | 96.9 | 11.5 | 3424.8 | 130.4 | 54.3 | 0.6 | 9752.7 | 380.9 |
| **100** | 32.5 | 0.9 | 124.8 | 4.1 | 4371.5 | 226.3 | 60.3 | 1.2 | NA | NA |

**Supplementary Table 1: Comparing compute times**

Compute times for CellRank (CR), Palantir (P), STEMNET (S) and FateID (F) for the 100k reprogramming dataset of ref.[63] to compute fate probabilities (Online methods). Values represent time in seconds. For CellRank, we additionally recorded the time it took to compute three macrostates. We repeated each computation 10 times to compute mean and standard error on the mean. FateID did not finish on 100k cells due to memory constraints.

| #cells (thousands) | CR (macro states) [mean] | CR (macro states) [std] | CR (lin. probs.) [mean] | CR (lin. probs.) [std] | S [mean] | S [std] | P [mean] | P [std] | F [mean] | F [std] |
|---|---|---|---|---|---|---|---|---|---|---|
| **10** | 9.78 | 0.2 | 9.9 | 0.09 | 0.23 | 0 | 9.74 | 0.06 | 2.55 | 0 |
| **20** | 15.21 | 0.07 | 12.64 | 0.1 | 0.46 | 0 | 14.15 | 0.09 | 9.81 | 0.62 |
| **30** | 20.36 | 0.06 | 15.53 | 0.08 | 0.68 | 0 | 20.02 | 0.19 | 18.5 | 0.01 |
| **40** | 25.49 | 0.08 | 18.26 | 0.07 | 0.91 | 0 | 27.43 | 0.34 | 34.88 | 0.12 |
| **50** | 30.56 | 0.13 | 21.02 | 0.1 | 1.14 | 0 | 36.89 | 0.28 | 52.72 | 0.35 |
| **60** | 35.63 | 0.09 | 23.83 | 0.09 | 1.36 | 0 | 47.38 | 0.55 | 73.43 | 1.85 |
| **70** | 40.71 | 0.05 | 26.68 | 0.06 | 1.59 | 0 | 53.24 | 0.81 | 98.92 | 1.28 |
| **80** | 45.86 | 0.08 | 29.63 | 0.13 | 1.81 | 0 | 66.28 | 1.2 | 124.65 | 1.96 |
| **90** | 51.01 | 0.11 | 32.55 | 0.15 | 2.04 | 0 | 81.19 | 1.08 | 155.1 | 3.07 |
| **100** | 56.13 | 0.11 | 35.52 | 0.14 | 2.26 | 0 | 98.26 | 0.83 | NA | NA |

**Supplementary Table 2: Comparing peak memory usage**

Peak memory usage for CellRank (CR), STEMNET (S), Palantir (P) and FateID (F) for the 100k reprogramming dataset of ref.[63] to compute fate probabilities (Online methods). Values represent peak memory usage in GiB. For CellRank, we additionally recorded the time it took to compute three macrostates. We repeated each computation 10 times to compute mean and standard error on the mean. FateID did not finish on 100k cells due to memory constraints. CellRank and Palantir make use of parallelisation to speed up their computations which increases peak memory usage (Online methods). We report decreased peak memory usage on 100k cells using a single core for Palantir and CellRank in Supplementary Table 3.

| #cells (thousands) | CR (macrostates) [mean] | CR (macrostates) [std] | CR (lin. probs.) [mean] | CR (lin. probs.) [std] | P [mean] | P [std] |
|---|---|---|---|---|---|---|
| **100** | 22.56 | 0.06 | 14.22 | 0.05 | 82.5 | 4.35 |

**Supplementary Table 3: Comparing peak memory usage on a single core**

Running the memory benchmark of Suppl. Tab. 2 on a single core on 100k cells for the methods which make use of parallelisation, CellRank (CR) and Palantir (P). Values represent peak memory usage in GiB. We repeated each computation 10 times to compute mean and standard error on the mean.
